# Specialized enteric glial cells coordinate intestinal motility

**DOI:** 10.1101/2023.06.07.544052

**Authors:** Marissa A. Scavuzzo, Katherine C. Letai, Jeyashri S. Rameshbabu, Alka Tomar, Isha K. Shah, Naya S. Alsouss, William J. Wulftange, Yuka Maeno-Hikichi, Jesse J. Zhan, Bruce W. Armstrong, Haleigh Paolucci, Begüm Aydin, H. Elizabeth Shick, Aakash K. Shah, Calia P. Thompson, Alsedeaq M. Hawamdeh, Aura Perez, Ying Xiong, Erin F. Cohn, Kevin C. Allan, Benjamin L.L. Clayton, Paul J. Tesar

**Affiliations:** Institute for Glial Sciences, Case Western Reserve University School of Medicine, Cleveland, Ohio 44106, USA; Department of Genetics and Genome Sciences, Case Western Reserve University School of Medicine, Cleveland, Ohio 44106, USA; Laboratory of Mucosal Immunology, The Rockefeller University, New York, New York 10065, USA

## Abstract

The enteric nervous system is a complex network of neurons and glia within the gut that coordinate gut motility. By optimizing single nucleus RNA-sequencing methods and spatial transcriptomics, we generated maps of the mouse duodenum and identified distinct molecular classes of enteric glia across the intestine with unique morphological and spatial identities. Here we show enteric glial functional specialization, with one myenteric subtype directly sensing force and expressing the mechanosensory ion channel PIEZO2. Genetic reduction of PIEZO2 in enteric glial populations enriched for this mechanosensory subtype led to defects in gastrointestinal motility. These results provide insight into the multifaceted functions of distinct enteric glial cell subtypes in maintaining gut health, and emphasize the importance of considering subtype-specific roles of enteric glia in a wide range of diseases and disorders.

## Main

The enteric nervous system (ENS) is often referred to as the ’second brain’ due to its complexity and independence from central control. The ENS is a complex network of neurons and glia embedded within the gastrointestinal tract, spanning the entire length from the esophagus to the rectum. The overwhelming majority of neurons and glial cells that exist outside of the brain are housed inside of the gut, where they populate every layer of tissue from the outside in. Enteric glial cells are found intramuscularly, in the mucosa beneath the epithelium, and organized into two ganglionated plexus layers, the myenteric and submucosal plexus (**Fig. 1a**).

**Figure 1.**
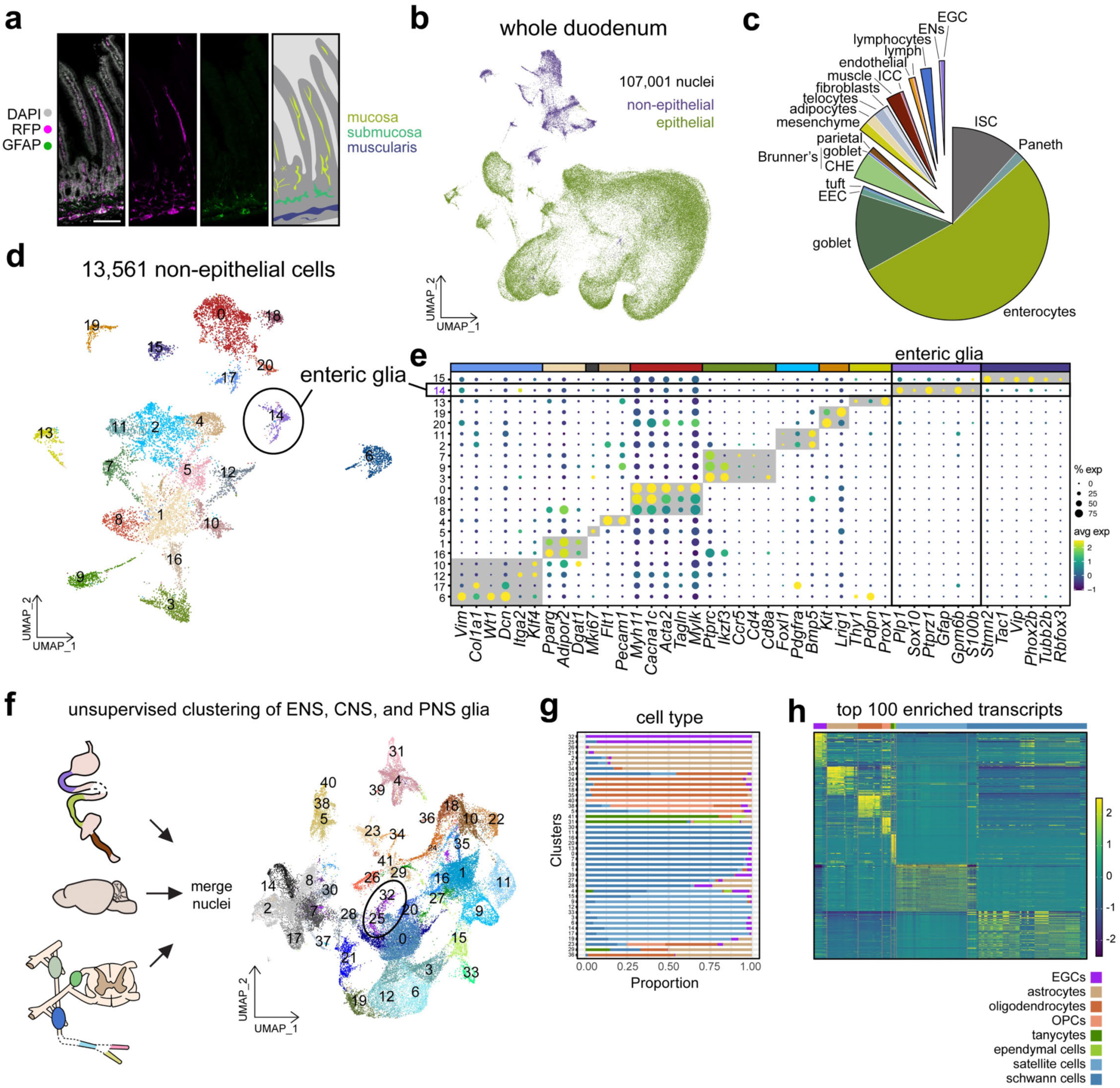
Enteric glia are a unique population of cells. **a**, Labeling of the enteric nervous system in the *Sox10-*cre; Ai14(RCL-tdT) mouse duodenum. RFP marks enteric nervous system cells, GFAP is shown in green, and nuclei are marked in white by DAPI. Scale bar = 100 um. **b**, snRNAseq of duodenal nuclei visualized by Uniform Manifold Approximation and Projection (UMAP), colored by cellular compartment (N=8 mice, n=107,001 nuclei). Unsupervised clusters shown in Extended Data Fig. 2. **c**, Proportion of cell types in the mouse duodenum from *CitraPrep* and snRNAseq. **d**, Unsupervised clustering of non-epithelial cells in the duodenum shown by UMAP. **e**, Cell type specific marker expression (columns) in clusters (rows) as shown in (d). Broad cell type classification is shown with dark gray boxes indicating from left to right mesenchymal cells, adipocytes, endothelial cells, musculature, lymphocytes, telocytes, interstitial cells of Cajal, lymphatic vasculature, enteric glia, and enteric neurons. Differentially expressed transcripts are detailed in Supplementary Table 1. The size of each circle indicates percentage of cells in the cluster that express the marker (≥1 UMI) while the color shows the average expression of transcript in cells. **f**, On left, schematic of study design. Enteric glia from cluster 14 represented duodenal glial cells. Individual datasets shown in Extended Data Fig. 2 and 3. On right, snRNAseq of glial cells spanning the central, peripheral, and enteric nervous systems colored by unsupervised clusters and visualized by UMAP (N=41 mice, n=47,998 nuclei). **g**, Proportion mapping of cell types to unsupervised clusters. Cell types are shown by color with key shown to the bottom right. Proportion of cells are shown on the y-axis while columns binned in the x-axis show unsupervised clusters. **h**, Top 100 differentially expressed transcripts enriched in each glial cell type. Individual transcripts are shown in rows, cell types are clustered together in columns with color annotation above. Datasets are provided in Supplementary Table 2.

Glial cells in the brain are known for their functional diversity. The gut is a highly complex and dynamic tissue, where cells are exposed to circulating factors, microbial metabolites, mechanical cues, and constant environmental changes. The glial cells in this tissue are vulnerable to a variety of signals and can interact with a wide assortment of cell types to execute a range of digestive functions. However, their diversity and physiological roles are just beginning to be explored. A comprehensive classification of all cells in the intestine is vital to delineate how enteric glial cells impact intestinal health and physiology.

In many other organs, single cell technologies have increased our understanding of cellular diversity in complex tissues and enabled the identification of specialized cell types with unique roles in tissue function^1–10^. However, efforts to understand the cellular diversity of the ENS have thus far focused on specific layers of the tissue or neuronal diversity^1,11–19^. The technical challenges inherent to profiling a digestive, nucleic acid-degrading environment have prevented the generation of a comprehensive transcriptional analysis of enteric glial cells across the gut from every layer of tissue at the single cell level (**Extended Data Fig. 1a**). Here, we set out to decode the global identity and diversity of enteric glial cells throughout the entire proximal-to-distal intestine. Functional interrogation of enteric glial subtypes demonstrates that subtypes of glia execute unique functions to regulate gastrointestinal physiology.

### A global atlas of the mouse duodenum

The biochemistry inherent to the adult mouse duodenum has hindered global transcriptional analysis at the single cell level due to an abundance of RNAses, ions, and a variable pH. Here, we successfully overcome these obstacles with an optimized iodixanol gradient and citric acid-based method we call *CitraPrep* that enabled us to obtain high quality nuclear RNA from single cells in the mouse duodenum (**Extended Data Fig. 1b-f**)^1,20–28^. Our protocol produced a high proportion of nuclei passing quality control metrics (**Extended Data Fig. 1g-i**). After filtering for low quality nuclei and doublets, we recovered 107,001 nuclei with an average of 7,249 UMIs per cell (**Fig. 1b**). We found differential expression of cell type specific genes across unsupervised clusters that spanned all major intestinal cell types (**Supplementary Table 1)**. Non-epithelial cell types, which made up 12.67% of total nuclei profiled, were selected for subclustering for higher resolution of cell types, including enteric neurons and glia (**Fig. 1c-e, Supplementary Table 1**)^29–32^.

### Enteric glial cells are regionally unique

We set out to assess whether individual enteric glial cells are transcriptionally analogous to other glial cell types in the body^2,33–37^. We performed snRNAseq on mouse cortical brain tissues to compare to our duodenal data set as well as our analyses of publicly available snRNAseq data (**Fig. 1f**)^24,38–41^. Isolating glial cell types from each tissue showed 7 glial cell types across 9 tissue regions extending throughout the body from the cortex (head) to the sural nerve (feet) (**Extended Data Fig. 2a-f**). Within this population of glial cells, we identified unsupervised clusters that were largely defined by cell type rather than study, sample, or tissue origin (**Extended Data Fig. 3a-g**). Enteric glia clustered mostly apart from all other cell types (**Fig. 1g)**^42,43^. We further identified and validated highly enriched, differentially expressed glial cell type specific markers (**Fig. 1h, Extended Data Fig. 3h-i, Supplementary Table 2**). Our data support that enteric glia are a transcriptionally distinct glial cell type.

We next analyzed snRNAseq from the ileum and colon to subset out enteric glial cells (**Extended Data Fig. 4a**)^1^. After merging duodenum, ileum, and colon glial subsets together, we found that unsupervised clusters separated by regional identity indicating enrichment for unique molecular profiles (**Extended Data Fig. 4b-e, Supplementary Table 3**)^44^. To validate our snRNA-seq results with an orthogonal approach, we optimized multiplexed error-robust fluorescence in situ hybridization (MERFISH) for parallel probing of duodenal and colon tissue (**Extended Data Fig. 4f-l**, **Supplementary Table 3**). We selected 140 transcripts that spanned regional diversity, confirming genes to be differentially expressed between the duodenum and colon (**Extended Data Fig. 4f-l, Supplementary Table 3**).

### Transcriptionally defined subtypes of enteric glia are spatially arranged

In our duodenal enteric glial dataset, we identified 6 molecularly distinct enteric glia subpopulations that cluster apart from enteric neurons and are differentiated by enriched expression of specific transcripts (**Fig. 2a-b**). For example, EGC1 is defined by enriched expression of the nuclear hormone receptor *Ppard,* EGC5 is defined by enriched expression of WNT ligands *Wnt6, Wnt4,* and *Wnt5a,* while EGC3 expressed intermediate markers of neurons and glia (**Fig. 2c**, **Extended Data Fig. 5a-b**)^45^. Immunostaining using tissue from *Sox10*-cre; RCL-tdTomato mice validated expression of these glial cell markers in neural crest derived cells (SOX10-positive cells; **Extended Data Fig. 5c**). Equipped with a transcriptional map of individual cells in the duodenum, we projected these markers across all major cell types to identify genes preferentially expressed in enteric glial cell subpopulations (**Fig. 2c, Extended Data Fig. 5d**). Using publicly available small intestinal spatial transcriptomics data, we identified the primary location of EGC subtypes (**Fig. 2d-g, Extended Data Fig. 5e-h, Supplementary Table 4)**. Using *Sox10*-cre; RCL-tdTomato mouse tissue, we show that the EGC1 subtype, marked by enriched PPARd, was overwhelmingly localized in the submucosa with very few in the tunica muscularis (**Fig. 2h-j, Extended Data Fig. 5i**). In our analysis of publicly available single-cell ENS datasets, we observed that mucosal-associated glia, such as those marked by *Ppard*, were often absent (**Extended Data Fig. 6a-j**, **Supplementary Table 4)**, highlighting that profiling glia only from the tunica muscularis leads to subtype specific dropouts in the datasets.

**Fig. 2.**
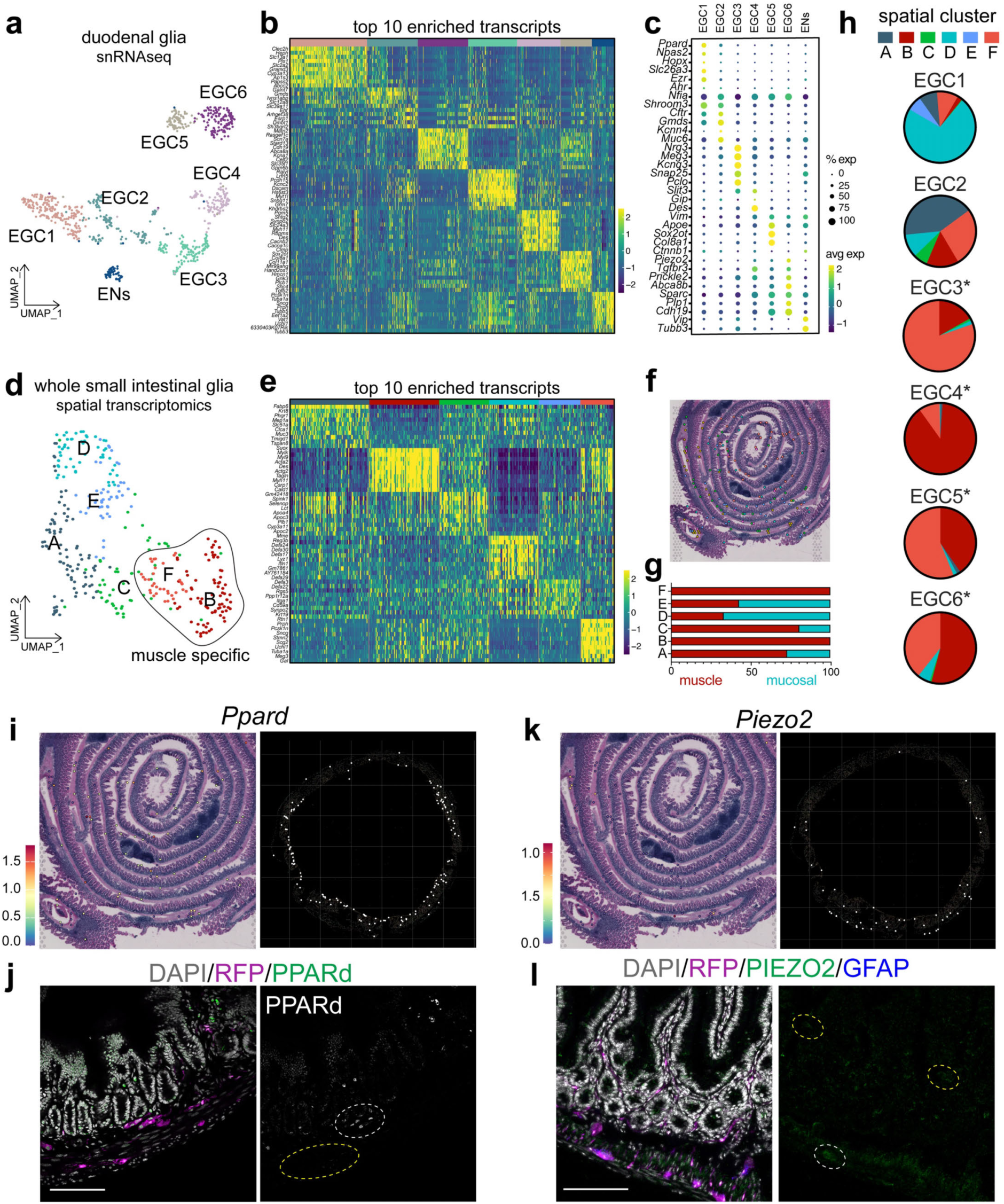
Transcriptionally unique glial cell types are spatially arranged. **a**, UMAP plot of subclustered duodenal ENS cells. Nuclei are colored by unsupervised clustering and annotated post hoc. **b**, Heatmap of the top 10 differentially expressed genes (rows) in each cluster (columns). Cluster identity is denoted by color above correlating with Fig. 2a. Datasets are provided in Supplementary Table 4. **c**, Cell type specific marker expression (rows) in clusters (columns). The size of each circle indicates percentage of cells in the cluster that express the marker (≥1 UMI) while the color shows the average expression of transcript in cells. **d**, UMAP of enteric glial cells that express *Plp1, Gfap, Sox10,* or *Gfra2* > 0.5 from small intestinal spatial transcriptomics from GSE227742. Nuclei are colored by unsupervised clustering. **e**, Heatmap of the top 10 differentially expressed genes (rows) in each cluster (columns) from spatial transcriptomics. Cluster identity is denoted by color above correlating with Fig. 2d. Datasets are provided in Supplementary Table 4. **f**, Spatial distribution of unsupervised clusters shown in Fig. 2d. **g**, Proportion mapping of enteric glia unsupervised cluster spatial identity. Regions are annotated by color and binned into mucosal or muscle. **h**, Mapping snRNA-seq unsupervised clusters to cluster identity from spatial transcriptomics by comparing differentially expressed genes between groups. Asterisk denotes groups that were majority associated with the muscle layer. **i**, Spatial distribution of EGC1 marker *Ppard* in enteric glial clusters from spatial transcriptomics small intestine (left) and whole MERFISH colon (right). Spatial transcriptomics retains expression data, with high expression red, moderate expression yellow, and low to no expression blue/purple. MERFISH shows binary expression data, with each dot representing a cell with *Ppard* transcript. **j**, Representative immunostaining of EGC1 in the adult *Sox10-*cre; RCL-tdT mouse duodenum. tdTomato is marked by RFP in magenta while nuclei are marked in white by DAPI. White circle shows co-expression of PPARd/RFP in the ENS while yellow circle shows myenteric RFP without PPARd. Scale bars = 100 um. **k**, Spatial distribution of EGC6 cell marker *Piezo2* in enteric glial clusters from spatial transcriptomics small intestine (left) and whole MERFISH colon (right). Spatial transcriptomics retains expression data, with high expression red, moderate expression yellow, and low to no expression blue/purple. MERFISH shows binary expression data, with each dot representing a cell with *Piezo2* transcript. **l**, Representative immunostaining of myenteric associated EGC6 cell markers in the adult *Sox10-*cre; RCL-tdT mouse duodenum. Intestinal compartments are labeled; MP is myenteric plexus while SMP denotes submucosal plexus. tdTomato is marked by RFP in magenta while nuclei are marked in white by DAPI. White circle shows co-expression of PIEZO2/GFAP in the ENS while yellow circles show PIEZO2 without GFAP outside of the ENS in the villi where known expression is observed in enterochromaffin cells. Scale bars = 100 um.

We identified a specific population of glial cells, EGC6, that exhibited enriched expression of *Piezo2,* with low expression in enteric neurons relative to glia (**Fig. 2c**). Visium spatial transcriptomic analysis, MERFISH, and immunostaining showed that PIEZO2+ neural crest derived cells were restricted to the myenteric plexus (**Fig. 2h, k-l, Extended Data Fig. 5i**)^1,17^. Analysis of *in vitro* and *in vivo* cell morphology showed the majority of EGC6 cells were intraganglionic in morphology (**Extended Data Fig. 7a-l).** Interactome analysis predicted that this population interacts with interstitial cells of Cajal (ICCs) and enteric neurons, two cell types also located in the myenteric plexus (**Extended Data Fig. 8a-b, Supplementary Table 5**). Functional enrichment analysis on the top 100 differentially expressed genes in this population compared to other enteric glia subtypes included transcripts associated with neuronal action potentials, further supporting a role for these cells in regulating neuronal activity (**Extended Data Fig. 8b-d**). Collectively, given the spatial location of this population, their predicted interactions, and their enriched expression of the mechanosensitive ion channel *Piezo2* we hypothesized these cells sense force to regulate physiology.

### Myenteric glial cells regulate gut motility via PIEZO2

Humans with *PIEZO2* loss-of-function mutations report a wide range of bowel dysfunctions including constipation and hardened stools^46,47^. Mechanical sensing via PIEZO2 from DRG somatosensory neurons that innervate the colon assist in sensing and reducing transit^46^; previous literature also suggests that enteric glia may promote transit through Ca^2+^ signaling^48–54^. Based on the significantly enriched PIEZO2 expression in EGC6, we set out to determine whether EGC6 cells directly sense force through the PIEZO2-Ca^2+^ axis to regulate aboral gut motility.

We found that the *Piezo2+* enteric glial subtype was present across the gastrointestinal tract (**Fig, 3a-c**). We next confirmed expression of PIEZO2 at the protein level in myenteric glial cells by its co-expression with SOX10 (marking all myenteric glia) and GFAP (marking a subset of myenteric glia including EGC6) in both mouse and human tissue (**Fig. 3d, Extended Data Fig. 8e-i**).

**Fig. 3.**
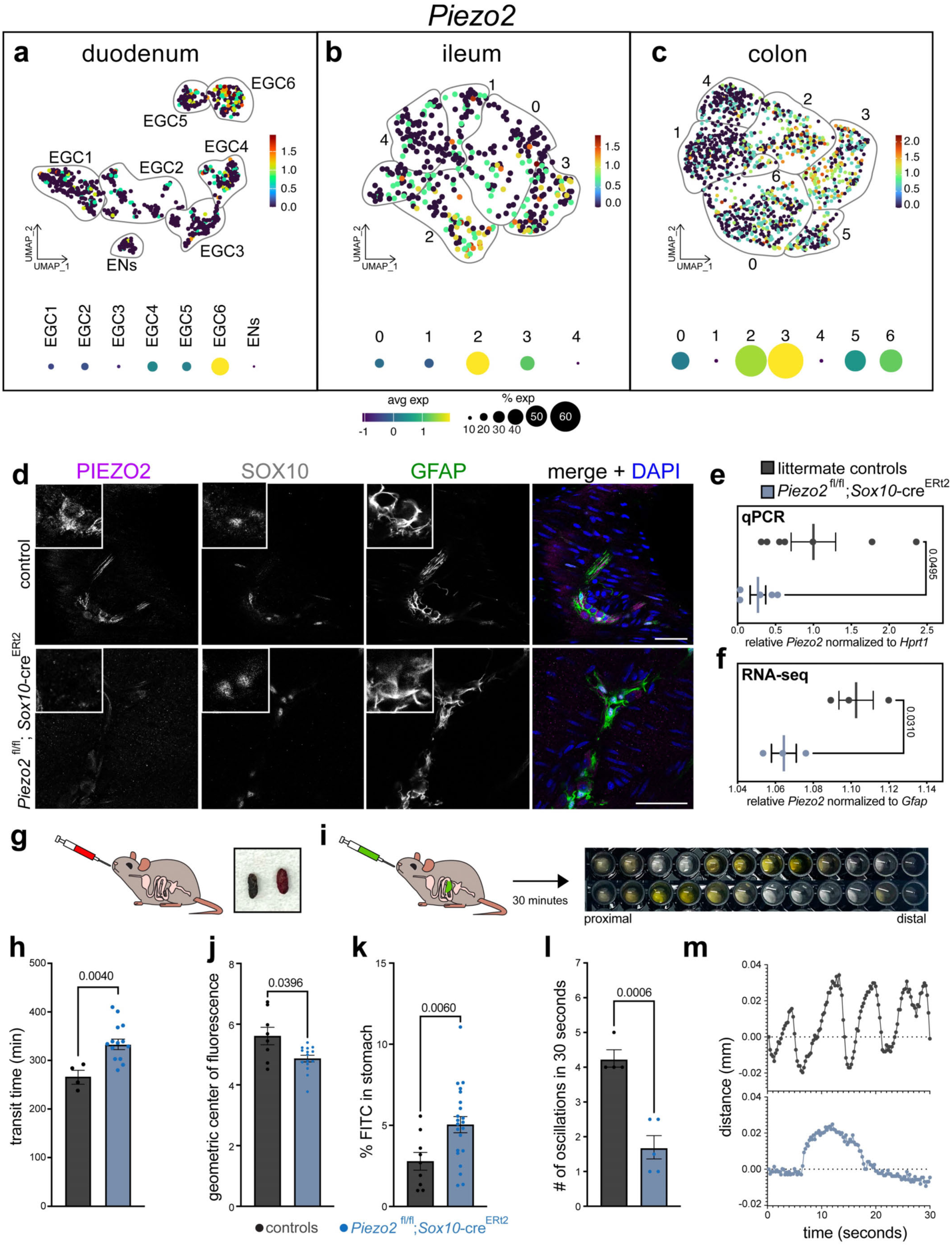
Mechanosensory enteric glial cells regulate gut motility. **a-c**, *Piezo2* expression in (**a)** duodenal, (**b**) ileal, and (**c**) colonic enteric glial cells, with expression projected upon the UMAP plot above a dot plot showing expression across unsupervised clusters. The size of each circle indicates percentage of cells in the cluster that express the marker (≥1 UMI) while the color shows the average expression of transcript in cells. **d**, Immunostaining of PIEZO2 (in magenta) in the tunica muscularis of the duodenum overlapping with pan-enteric glia marker SOX10 (in white) and subtype enteric glia marker GFAP (in green) with enteric glia specific adult PIEZO2 knockout shown on the bottom in Piezo2^fl/fl^; *Sox10*-cre^ERt2^ mice. Scale bar = 50 um. Inset shows a PIEZO2+ enteric glial cell co-expressing GFAP and SOX10, with PIEZO2 depletion in the cre/lox mice. Note positive staining of PIEZO2 in GFAP-cells with sensory fiber morphology. **e**, qPCR of *Piezo2* normalized to *Hprt1* from the longitudinal muscle / myenteric plexus of Piezo2^fl*/fl*^; *Sox10-*cre^ERt2^ and littermate controls in biological replicate relative to control. Center line shows mean with standard error and individual animals represented as dots. **f**, RNA-seq of *Piezo2* normalized to *Gfap* from the longitudinal muscle / myenteric plexus of Piezo2^fl*/fl*^; *Sox10-*cre^ERt2^ and littermate controls in biological replicate relative to control. Center line shows mean with standard error and individual animals represented as dots. **g,** Schematic showing oral gavage of carmine red dye. Carmine red dye in fecal pellets marks transit time. **h**, Total gastrointestinal transit time measuring time from gavage of carmine red dye to the appearance of red stool in littermate controls (N=4) and Piezo2^fl/fl^; *Sox10*-cre^ERt2^ mice (N=14). **i**, Schematic and gastrointestinal transit time assay with representative results from the stomach and small intestine. Each well represents contents from one equal sized segment of the small intestine, with the first well showing the stomach and the last well the cecum. **j**, The geometric center of fluorescence shows the location of the bolus in the small intestine in littermate controls (N=8) and Piezo2^fl/fl^; *Sox10*-cre^ERt2^ mice (N=15). **k**, Gastric emptying in littermate controls (N=9) and Piezo2^fl/fl^; *Sox10*-cre^ERt2^ mice (N=22). **l**, Quantification showing the number of oscillations over a 30 second time frame in littermate controls (N=4) and Piezo2^fl/fl^; *Sox10*-cre^ERt2^ (N=5) mouse tissue stimulated with 1dyn/cm^2^ shear stress *ex vivo*. **m**, Representative oscillations recorded from *ex vivo* longitudinal muscle with myenteric plexus under shear stress stimulation.

To determine whether PIEZO2 conferred functional specialization to mechanically sensitive EGC6, we crossed *Piezo2^fl/fl^* mice^55–61^ to the *Sox10-*cre^ERt2^ driver line to deplete PIEZO2 in all myenteric plexus glia, including EGC6 cells. We treated mice with tamoxifen at three weeks of age to selectively deplete PIEZO2 in adult glial cells. Three weeks after tamoxifen administration, we confirmed *Piezo2* depletion by immunostaining, qPCR, and bulk RNA-seq, which showed reduction of PIEZO2 from the tunica muscularis and from SOX10+ myenteric glia in the mouse intestine (**Fig. 3d-f**).

To understand how PIEZO2 loss in enteric glia affects gastrointestinal functions, we gavaged mice with a non-absorbable dye and measured the length of time for the colored fecal pellet to appear (**Fig. 3g**). In mice lacking PIEZO2, we observed a significant delay in overall transit time compared to co-housed, littermate controls (**Fig. 3h**), along with a decrease in fecal water content (**Extended Data Fig. S9a**). PIEZO2 deficient mice did not exhibit any differences in epithelial permeability, weight, or intestinal length compared to controls (**Extended Data Fig. 9b-e**). To gain resolution and locate where contents were getting stuck, we gavaged mice with a non-absorbable fluorescent dye and collected luminal contents from equally sized segments of the gut after 30 minutes later (**Extended Data Fig. 9f)**. We observed a shift in the distribution of dye in the gastrointestinal tract of *Piezo2 ^fl/fl^; Sox10-*cre^ERt2^ mice, with a significant defect in gastric emptying (**Fig. 3i-k**). To determine whether the delay was due to changes in gut motility, we performed *ex vivo* contractility assays after loss of PIEZO2 using an intestine-on-a-chip platform. In response to shear stress, we observed a regular oscillating contractile pattern that was completely abolished upon treatment with the voltage-dependent L-type calcium channel inhibitor nifedipine (**Extended Data Fig. 9g-h**). In contrast, while *Piezo2 ^fl/fl^; Sox10-*cre^ERt2^ tissue had comparable amplitude or strength of contractions, these tissues exhibited a decreased frequency and arrhythmicity compared to littermate controls (**Fig. 3l-m**). Finally, we broadly measured acetylcholine and nitric oxide (two neurotransmitters known to directly modulate peristalsis) in the muscle layer of the small intestine in *Piezo2 ^fl/fl^; Sox10-*cre^ERt2^ and found disorganized levels of nitric oxide and acetylcholine (**Extended Data Fig. 9i-j**).

### A subset of enteric glial cells act as biomechanical sensors

To see if enteric glia were competent to respond to mechanical force, we engineered a microfluidic biochip to mechanically stimulate intestinal tissue during simultaneous calcium imaging (**Fig. 4a, Extended Data Fig. 9k**). As expected, we observed differential calcium responses across enteric glia^62^, however a subset of myenteric glial cells responded only after stimulation with force (**Fig. 4b-c**). In the majority of these enteric glia, activation was reversed upon treatment with D-GsMTx4, a peptide inhibitor of PIEZO2 and TRPC1/6 (**Fig. 4c, Supplementary Video 1**). Immunostaining of these mechanosensory cells post-hoc showed that they expressed PIEZO2 and GFAP (**Extended Data Fig. 9l**), and that enteric glia express low levels of *Trpc1* and *Trpc6* (**Extended Data Fig. 9m**). To increase the throughput of this analysis, we cultured enteric glial-enriched primary mouse cells to image before, during, and after a pulse of shear stress stimulation at 1 dyn/cm^2^, which showed a significant response and rapid return to baseline across biological replicates (**Fig. 4d-h, Extended Data Fig. 9n-o, Supplementary Video 2**). We identified the first and last responding cells in our imaging of gut tissue and found that the first responding cells were PIEZO2+, GFAP+, SOX10+ (**Fig. 4i-j, Supplementary Video 3**). We then mapped the activity of these cells throughout space-time to demonstrate the precise temporal dynamics of individual cells as their activity peaks over time (**Fig. 4k**), illustrating a spreading phenomenon that starts at a hub of mechanosensitive, PIEZO2+ enteric glial cells. These results support that a specialized mechanosensory subpopulation of enteric glia exists in the myenteric plexus of the small intestine.

**Fig. 4.**
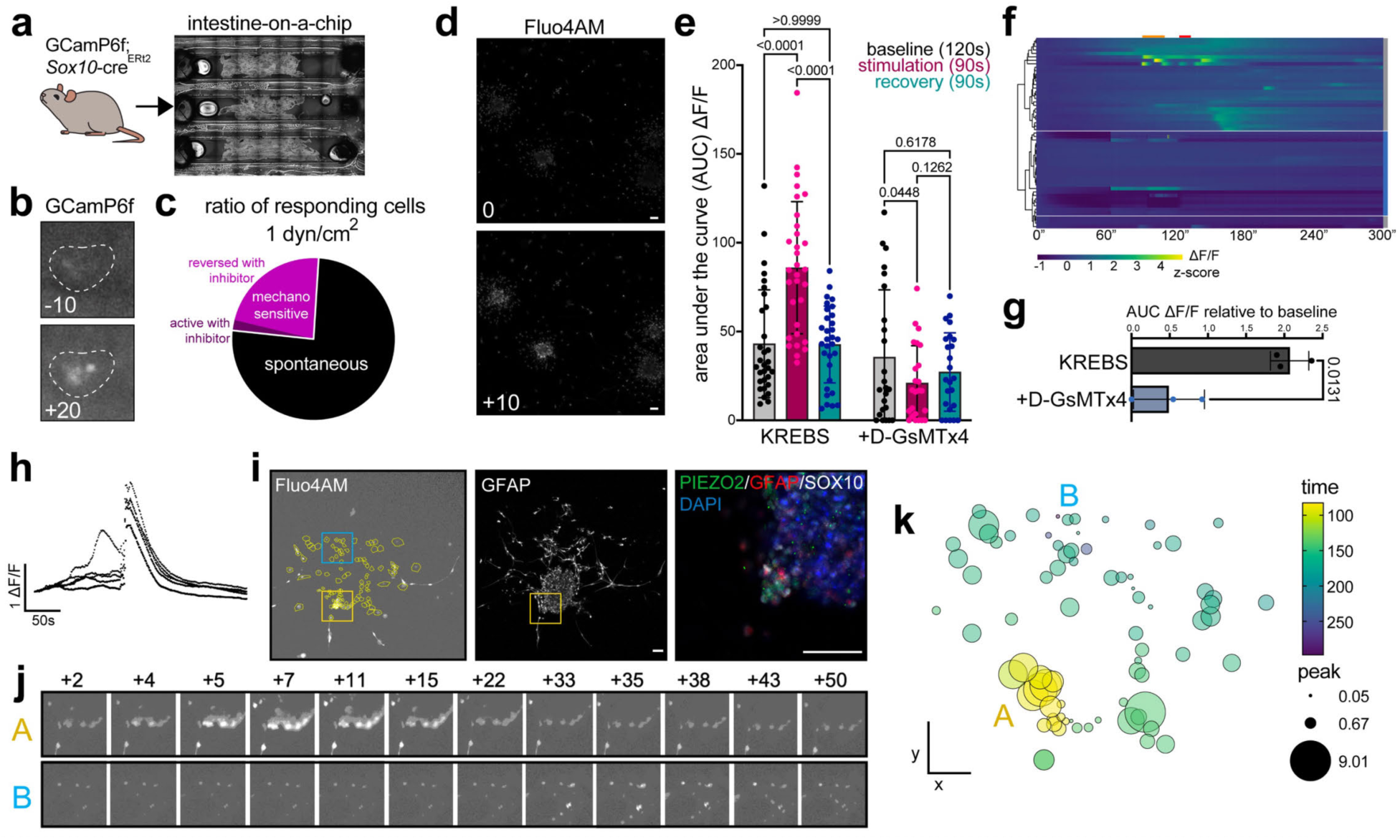
PIEZO2+ enteric glial cells initiate force-induced calcium waves. **a**, Intestine-on-a-chip platform engineered on coverslips for high resolution, live Ca^2+^ imaging under shear stress with microfluidics. Longitudinal muscle with myenteric plexus was dissected from mice with the calcium indicator GCamP6f expressed in enteric glia via tamoxifen driven expression of Cre in *Sox10*+ cells. Brightfield image shows an overview of the coverslip with tissues in chambers. **b**, Representative image of mechanoresponsive enteric glia in intact tissue 10 seconds before and 20 seconds after 1dyn/cm^2^ shear stress is applied. **c**, Pie chart shows quantification of cells responding only at baseline (without shear stress), only after application of shear stress at 1dyn/cm^2^ and reversal of this response with 10μM D-GsMTx4, which inhibits PIEZO2. Cells that are responsive in both conditions are annotated, while cells active in the presence of inhibitor D-GsMTx4 at rest, after stimulation, or in both conditions are shown in a bracket. **d**, Representative image of mechanoresponsive enteric glia in culture at the onset of and 10 seconds after 1dyn/cm^2^ shear stress is applied. **e**, Quantification of enteric glial responses in different phases of stimulation as shown in Extended Data Fig. 8h measuring area under the curve. Each dot is an individual cell from three biological replicates; individual animal data shown in Fig. 4g. Bar plot shows mean, error bars show SEM, and P values are from two-way ANOVA mixed-effects model with multiple comparisons. **f**, Heatmap showing temporal calcium dynamics of individual cells. Time is plotted on the x axis in seconds, with cells shown in rows. Cells are grouped via hierarchical clustering with the majority of KREBS (gray) and 10μM D-GsMTx4 (blue) grouping together. The same cells are shown with and without 10μM D-GsMTx4. The orange bar above the plot depicts the removal of the clip and the red bar depicts the 10 seconds of 1 dyn/cm^2^ shear stress. **g**, Quantification of enteric glial cell responses upon stimulation, relative to KREBS control. Each dot is a biological replicate. **h**, Representative calcium traces from enteric glial cells. **i**, Left, image of cells during calcium experiments with a yellow box around early responders and a blue box around late responders. Center, GFAP immunostaining of the same cells. Right, expanded view of early responding cells within yellow box with immunostaining for PIEZO2 in green, GFAP in red, SOX10 in white, and nuclei marked by DAPI in blue. Scale bar = 50 um. **j**, Expanded view of early responding cells (A) and later responding cells (B) over a timecourse after shear stress application at 1 dyn/cm^2^. **k**, Calcium responses mapped over space-time. Spatial coordinates are mapped in 2D on the x and y axis, while peak time is illustrated colorimetrically from early (yellow) to late (purple). The size of the circle shows the peak ι1F/F response level.

### PIEZO2-mediated release of enteric gliotransmitters in response to force

To model physiological tension that glial cells in the tunica muscularis would experience *in vivo,* we used a cell stretching bioreactor system to determine what metabolites enteric glial cells release upon stimulation with force, and which of these were dependent on PIEZO2 (**Fig. 5a**)^47,63^. There was no significant change in the level of lactate dehydrogenase in control nor *Piezo2* deficient cells, with low absolute levels in both conditions, suggesting cell viability was not impacted by stretch (**Fig. 5b**). Sequencing the transcriptome of the tunica muscularis from littermate controls or *Piezo2*^fl/fl^; *Sox10-*cre^ERt2^ mice three weeks after tamoxifen induction showed 43 differentially expressed transcripts (*p*adj<0.05, **Extended Data Fig. 9p, Supplementary Table 6)**. These transcripts included many immunomodulatory genes including the inducible nitric oxide synthase 2, *Nos2,* which was decreased in depletion mice, but, as anticipated, no notable transcripts associated with neurotransmission or motility. After loss of PIEZO2 from enteric glial cells we did not observe changes in the levels of nitric oxide (**Fig. 5c**) or acetylcholine from glia themselves (**Fig. 5d**). Together, this indicates that the tissue level neurotransmitter changes we found and resulting motility differences were secondary to other post-transcriptional glial factors.

**Fig. 5.**
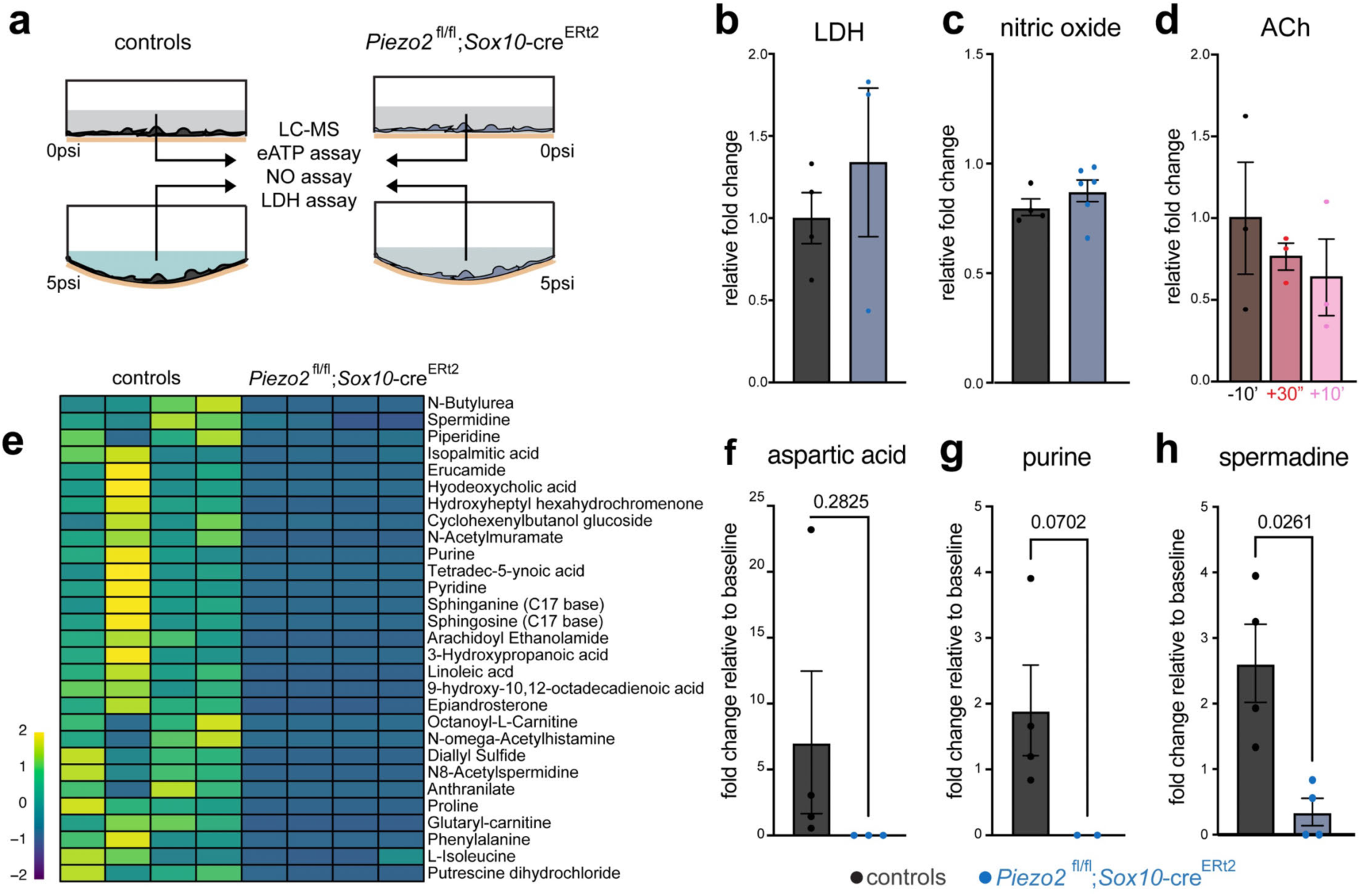
Mechanical activation via PIEZO2 drives purine and polyamine release from enteric glia. **a,** Schematic of cells plated in cell stretching bioreactor systems to collect conditioned media after 10 minutes at rest, application of tensile strain at 5psi, followed by 30 seconds of incubation. **b**, Fold change in lactate dehydrogenase (LDH) release after 5psi tensile strain in littermate control (N=3) and Piezo2^fl*/fl*^; *Sox10-*cre^ERt2^ mice (N=3). **c**, Fold change in nitric oxide release upon tensile strain at 5psi relative to baseline in enteric glial cells from littermate controls and Piezo2^fl*/fl*^; *Sox10-*cre^ERt2^ mice. **d**, Acetylcholine levels at baseline (-10m), after acute tensile strain (30s) and extended release (+10m) measured by untargeted liquid chromatography-mass spectrometry. Data is from three biological replicates. **e**, Untargeted metabolomics from 17 samples filtered for force-dependent compounds lost after PIEZO2 knockout. Heatmap showing release of compounds after stretch normalized to baseline in littermate controls (N=4) and Piezo2^fl/fl^; Sox10-cre^Ert2^ (N=4). Datasets in Supplementary Table 7. **f-h**, Fold change of (**f**) aspartic acid, (**g**) purines, and (**h)** spermadine relative to baseline. Each dot shows a biological replicate, error bars show SEM, and P values are from Welch’s two-sided t tests.

We performed untargeted metabolomics to identify factors released by glial cells before and after stretch in control and *Piezo2*^fl/fl^; *Sox10-*cre^ERt2^ cells. We identified metabolites with force-dependent release, with levels increasing in the media after stretch (**Supplementary Table 7**). We then identified a cohort of these whose force-dependent release was lost when PIEZO2 function was abolished (**Fig. 5e**). Enteric glia can directly release factors like ATP, nitric oxide, and y-Aminobutyric acid (GABA)^64,65,66^. After depletion of *Piezo2* from enteric glia, cells exhibit a diminished release of aspartic acid, purines (ATP), and polyamines (spermidine) upon mechanical stimulation (**Fig. 5f-h**). Thus, myenteric glia can sense force through PIEZO2 and release gliotransmitters upon mechanical stimulation.

Smooth muscle cells and enteric neurons are abundant in the tunica muscularis. To determine what cell types PIEZO2-dependent gliotransmitters can act upon, we generated primary cultures of smooth muscle cells and enteric neurons (**Fig. 6a,c**), two cell types predicted to interact with EGC6 (**Extended Data Fig. 8a-d**). We observed minimal calcium responses to candidate gliotransmitters in smooth muscle cells, with a slow and sustained approximately 2-fold increase in signal after administration with spermidine and BzATP, an agonist for purinergic receptors with high affinity for P2X1, P2X3, and P2X7^67^ (**Fig. 6b**). A small subset of enteric neurons exhibited robust activation (approximately 9-fold) upon stimulation with spermidine, while nearly all enteric neurons were activated by BzATP (approximately 5-fold; **Fig. 6d,e**). Enteric neurons are highly diverse. To gain insight into which enteric neurons were competent to respond to purines and polyamines, we analyzed the transcriptomes of nuclei from neurons in the tunica muscularis of the colon and small intestine (**Supplementary Table 8**)^1,68^. We identified 8 known enteric neuron subtypes based on marker expression (**Fig. 6f**). Intrinsic primary afferent neurons (IPANs), marked by expression of *Ano2, Calcb,* and *Nmu*, initiate intrinsic reflexes that activate ascending cholinergic circuits and recruit descending nitrergic circuits, ultimately modulating excitatory and inhibitor motor neurons and thus peristalsis^69–72^. We found enriched expression of putative polyamine receptors in neurons expressing IPAN-associated markers (**Fig. 6f**), including the inward rectifier potassium channel *Kcnj12* (KIR2.2)^73,74^. We immunostained the enteric neuron cultures used for calcium imaging and validated KIR2.2 expression at the protein level in ANO2+ cells (**Fig. 6g**). In contrast, putative purine receptors were expressed broadly across neuron classes, including IPANs, excitatory motor neurons, and inhibitory motor neurons (**Fig. 6f**). This shows that candidate PIEZO2-dependent glial mediators can activate enteric neurons.

**Fig. 6.**
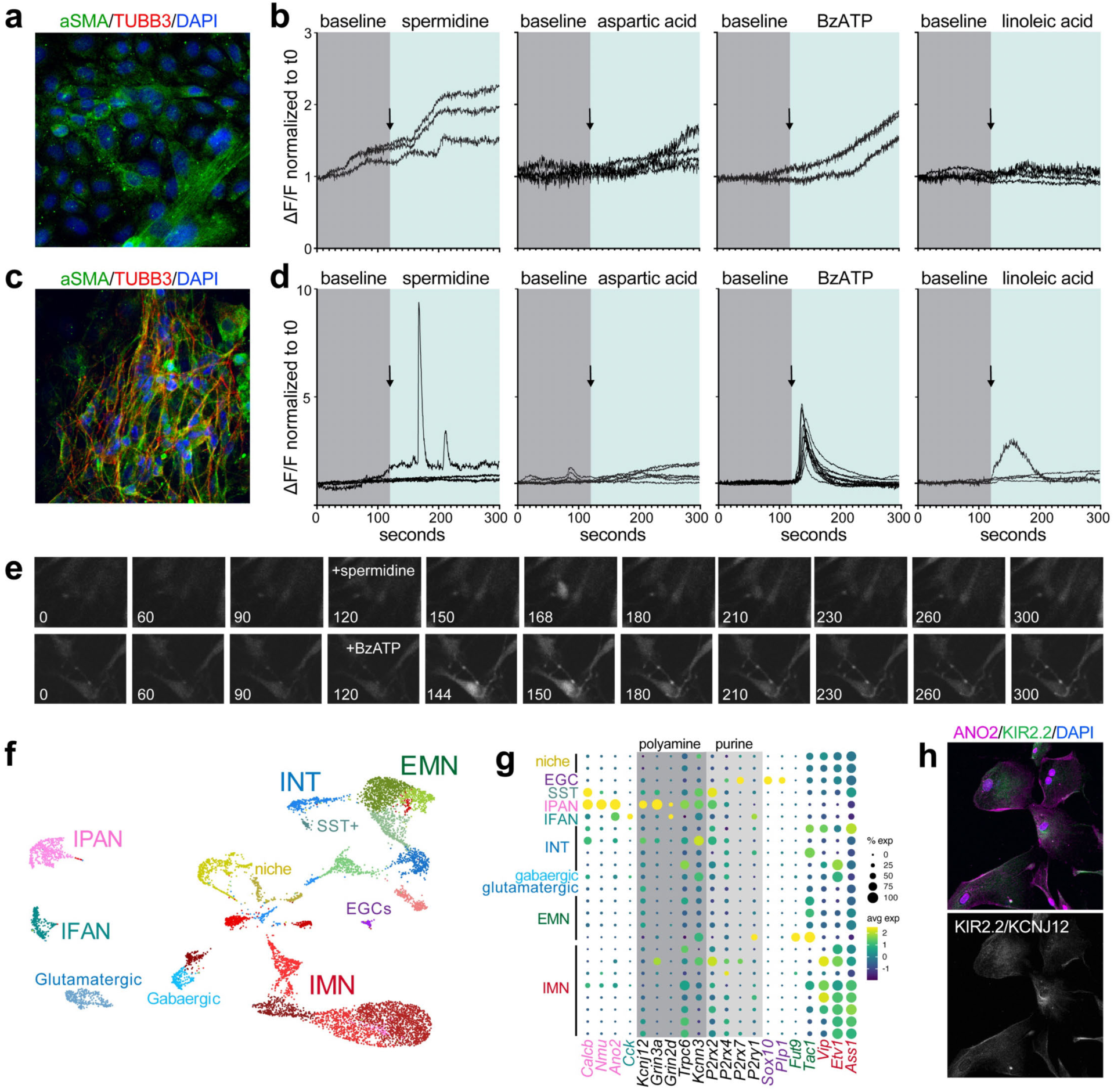
Purines and polyamines are force-dependent gliotransmitters that activate enteric neurons. **a,** Representative immunostaining of smooth muscle cultures used for live imaging, with aSMA marking smooth muscle in green and TUBB3 marking neurons in red. DAPI marks nuclei in blue. **b,** Calcium traces of smooth muscle cells before and after stimulation with candidate gliotransmitters. **c** Representative immunostaining of myenteric neuron cultures used for live imaging, with aSMA marking smooth muscle in green and TUBB3 marking neurons in red. DAPI marks nuclei in blue. **d,** Calcium traces of enteric neurons before and after stimulation with candidate gliotransmitters. **e,** Representative calcium images over a timecourse of enteric neurons before and after stimulation with spermidine (top) and BzATP (bottom). **f,** UMAP showing snRNA-sequencing of enteric neurons in the tunica muscularis. **g,** Dot plot showing transcript levels of marking enteric neuron subtypes (colors matching those in Fig. 6f) and putative purine and polyamine receptors (shaded gray and annotated above). Columns show transcripts while rows show enteric neuron subtypes. The size of each circle indicates percentage of cells in the cluster that express the marker (≥1 UMI) while the color shows the average expression of transcript in cells. **h,** Representative immunostaining validating protein level expression of polyamine receptor KIR2.2 (*Kcnj12)* in intrinsic primary afferent neurons (IPANs) marked in magenta by ANO2.

### A specialized enteric glial cell subpopulation regulates intestinal motility

Our expression data combined with tracing experiments suggested that *Sox10*-cre^ERt2^ targeted the majority of myenteric glia, while *Plp1-*cre^ERt2^ and *Gfap-*cre^ERt2^ mark partially overlapping, but not identical, enteric glial populations (**Fig. 7a-g**). With this in mind, we used these genetic drivers (*Plp1-*cre^ERt2^ and *Gfap-*cre^ERt2^) crossed with *Piezo2 ^fl/fl^* to specifically deplete *Piezo2* in subsets of enteric glial cells. In mice with loss of PIEZO2 from *Gfap+* cells, we detected a significant delay in transit time, and significant retention of dye in the stomach, phenocopying the global myenteric glia knockdown of PIEZO2 (**Fig. 7h**). In contrast, *Plp1-*driven loss of PIEZO2 had no phenotypic difference from littermate controls (**Fig. 7i**). Importantly, reducing PIEZO2 in *Aldh1l1*-positive astrocytes (**Extended Data Fig. 10a-c**), which are glial cell types outside the ENS, did not exhibit differences in whole gut transit time (**Extended Data Fig. 10d**), bolus location (**Extended Data Fig. 10e**), dye retained in stomach (**Extended Data Fig. 10f**), or body weight (**Extended Data Fig. 10g**). These data show that functionally specialized enteric glial cells exist in the gut and fine-tune intestinal physiology.

**Fig. 7.**
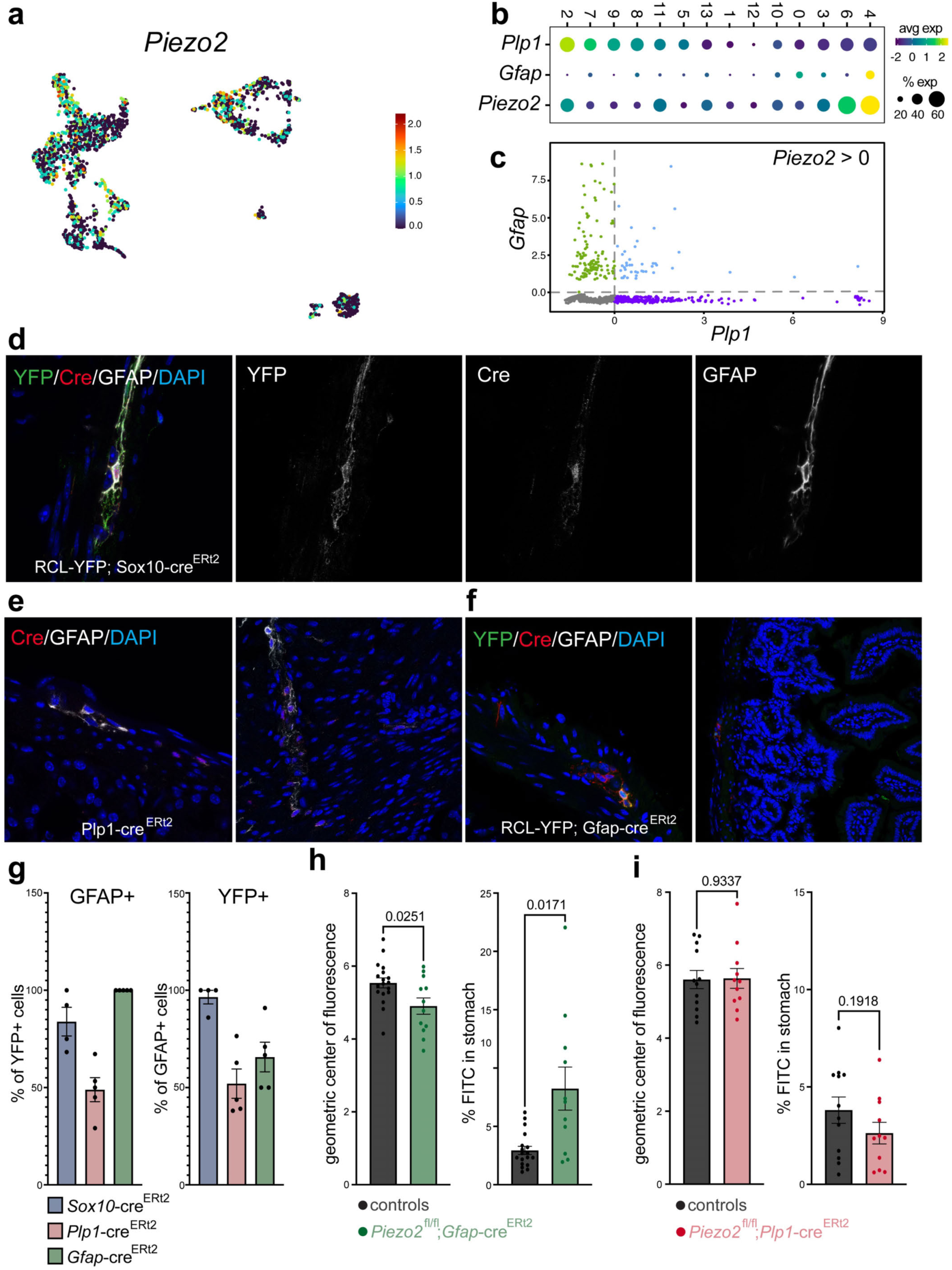
Enteric glial cells exhibit functional specialization. **a**, Expression of *Piezo2* projected across a UMAP of enteric glial cells merged from full thickness duodenum, ileum, and colon. **b**, Dot plot showing expression of *Piezo2, Gfap,* and *Plp1* in enteric glial cells throughout the intestine. The size of each circle indicates percentage of cells in the cluster that express the marker (≥1 UMI) while the color shows the average expression of transcript in cells. **c**, Comparison of expression levels of *Gfap* and *Plp1* in *Piezo2* expressing enteric glial cells > 0. Green dots are cells with expression of *Gfap* but no *Plp1* transcript, purple dots are cells with expression of *Plp1* but no *Gfap* transcript, blue dots are cells that express some level of both *Gfap* and *Plp1,* and gray dots are cells that do not have detectable transcripts of either *Gfap* or *Plp1.* **d**, Representative immunostaining of RCL-YFP; Sox10-cre^ERt2^ mouse duodenal tissue after our induction regimen. Green marks YFP cells where recombination has occurred, red marks Cre, and white marks GFAP+ glial cells. Nuclei are marked in blue by DAPI. **e**, Representative immunostaining of Plp1-cre^ERt2^ mouse duodenal tissue after our induction regimen. Cre is marked in red, GFAP+ glia in white, and nuclei in blue by DAPI. **f**, Representative immunostaining of RCL-YFP; Gfap-cre^ERt2^ mouse duodenal tissue after our induction regimen. Cre is marked in red, GFAP+ glia in white, and nuclei in blue by DAPI. **g**, Targeting efficiency of tamoxifen inducible Cre drivers in our experiments and *Gfap/Plp1* driver overlap. Left, the percent of YFP (targeted) cells that co-stain with GFAP. Center, the percent of GFAP+ cells that are targeted by each driver. Right, percent of YFP (targeted) cells that are negative for GFAP. Dots are individual mice showing mean with SEM. **h**, On left, the geometric center of fluorescence shows the location of the bolus in the small intestine in littermate controls (N=18) and Piezo2^fl/fl^; *Gfap*-cre^ERt2^ mice (N=12). On right, gastric emptying in littermate controls (N=18) and Piezo2^fl/fl^; *Gfap*-cre^ERt2^ mice (N=11). **i**, On left, the geometric center of fluorescence shows the location of the bolus in the small intestine in littermate controls (N=12) and Piezo2^fl/fl^; *Plp1*-cre^ERt2^ mice (N=11). On right, Gastric emptying in littermate controls (N=12) and Piezo2^fl/fl^; *Plp1*-cre^ERt2^ mice (N=11).

## Discussion

Our findings significantly advance our understanding of the unique and functionally specialized roles enteric glial cells play in digestion. By employing an optimized snRNA-seq approach, we were able to delineate the transcriptional and spatial diversity of enteric glia across the intestine. This characterization of enteric glial cells shows that they are not merely peripheral counterparts of other glia, but represent a distinct and transcriptionally unique group of cells with specialized functions crucial for maintaining gastrointestinal physiology.

We found that enteric glia are regionally distinct, with cells in the duodenum and colon possessing distinct molecular signatures. This regional specialization points to the importance of glial cells in modulating local gut environments, potentially influencing everything from nutrient absorption to motility^42,43,75^. In the small intestine, we identified 6 molecularly distinct populations of enteric glia, each exhibiting unique transcriptional profiles and spatial distributions throughout the layers of tissue. This classification underscores the necessity of a comprehensive understanding of glial cell diversity in the gut and the use of approaches to investigate cells throughout every layer of the tissue.

The identification of enteric glial EGC6 cells marked by enriched expression of PIEZO2 expands our understanding of mechanosensitivity within the ENS. These EGC6 cells not only serve as mechanosensors, but also modulate intestinal motility through the release of signaling molecules like purines and polyamines in response to mechanical stimuli. These cells, which we found respond early to force, illustrate a sophisticated network in which glia respond to biomechanical changes and relay information to neighboring cells. The EGC6 enteric glial cells are a molecularly, morphologically, spatially, and functionally unique cell type that integrate mechanical cues and then spread this information. The loss of PIEZO2 in these cells resulted in significant alterations in gut motility, indicating their critical role in maintaining normal gastrointestinal function. Importantly, the *Plp1-*cre^ERt2^ and *Gfap-*cre^ERt2^ drivers, while targeting different populations of cells, do overlap with many cells co-expressing *Gfap* and *Plp1* together. In addition, *Piezo2* transcript was found in both *Gfap+* and *Plp1+* cells. In future studies it will be important to use intersectional genetic technologies to specifically target transcriptionally defined subsets of cells for recombination. In addition, while the *Sox10*-cre^ERt2^ line was characterized in this paper, it is important to note that inducible Cre mouse lines rely on tamoxifen-dependent recombination that can vary between animals, tissues, and cells. Combined with recent findings that peripheral nerves express PIEZO2 and regulate the speed of gut transit^58^ and past studies showing PIEZO2 in enterochromaffin cells^76^ and intrinsic primary afferent neurons^1^, EGC6 cells may coordinate with other systems to provide an elegant stop and go signaling system. These overlapping mechanisms may have been driven to support the complexity of gut motility and organism survival. Our work adds a new layer of complexity to our understanding of how the ENS integrates sensory information to regulate digestive processes.

While glia in the gut are unique from other glia in the body, they express many of the same risk genes involved in neurological conditions. In fact, over 90% of high risk autism spectrum disorder genes are expressed in the enteric nervous system^77,78^. Gut disturbances are unusually common in individuals with diseases ranging from multiple sclerosis, schizophrenia, Parkinson’s disease, and autism^79–81^. In fact, individuals with Parkinson’s disease commonly develop gastroparesis^82^, a defect in gastric emptying. We found evidence that enteric glial cells may play a role in the regulation of gastric emptying. The molecular and functional characterization of enteric glia and their specialized subtypes carries important implications for understanding gut health and disease. Dysfunctional glial signaling is increasingly recognized as a contributing factor in various gut disorders, including inflammatory bowel disease and visceral pain^83–87^. By showing the specific roles of EGC6 in sensing mechanical forces and regulating motility, our findings could pave the way for new therapeutic strategies targeting glial cells to restore normal gut function.

## Methods

### Animals

Animal studies were approved by the Case Western Reserve University School of Medicine Institutional Animal Care and Use Committee. Mice were housed at 22–24°C with a 12 hour light/12 hour dark cycle with standard chow (Lab Diet Pico Lab 5V5R, 14.7% calories from fat, 63.3% calories from carbohydrate, 22.0% calories from protein) and water provided ad libitum unless otherwise indicated. The generation of *Plp1*-eGFP^88^, *Plp1-*cre^ERt289^, *GFAP-*cre^ERt290^, *Sox10-*cre^ERt291^, *Sox10-*cre^92^, *Aldh1l1-*cre^ERt2^; Ai14(RCL-tdT)^93^, Ai95(RCL-GCaMP6f)-D^94^, *Piezo2-*eGFP-IRES-cre^95^, *Rosa26*iDTR^96^, and floxed *Piezo2 (Piezo2^loxP/loxP^)*^95^ mice have been described previously. All mice were of C57Bl/6 background. With the exception of *Plp1-*eGFP mice obtained from Dr. Wendy Macklin, all other strains were obtained from The Jackson Laboratory. Animals were bred for mixed litters to compare phenotypes between littermates co-housed in the same cage. For all analyses, the date of birth was assigned as postnatal day 0. All genotyping was performed by Transnetyx.

### Administration of tamoxifen

For *Piezo2* reduction studies, 4-hydroxytamoxifen was dissolved in ethanol (100 mg/mL) at 65 °C for approximately 10 minutes with intermittent vortexing. Once the 4-hydroxytamoxifen solution was clear, corn oil was added (final concentration 20 mg/mL) and solution was either incubated overnight at 37 °C shaking or for approximately 15 minutes at 65 °C with intermittent vortexing to promote reconstitution. Solution was stored for up to 2 weeks at 4 °C and restricted to three cycles of heating to prevent loss of 4-hydroxytamoxifen activity. 4-hydroxytamoxifen was delivered to *Piezo2^loxP/loxP^*; *Plp1-*cre^ERt2^, *Piezo2^loxP/loxP^*; *GFAP-*cre^ERt2^, and *Piezo2^loxP/loxP^*; *Sox10-*cre^ERt2^ mice on postnatal day 21 (3 weeks) at 75 mg/kg by intraperitoneal injection once daily for 5 days and collected 3 weeks after first injection for analysis. RCL-*GCaMP6f*; *Sox10-*cre^ERt2^ mice were injected between 2 and 4 months at 75 mg/kg by intraperitoneal injection once daily for 5 days. All control mice were littermates, co-housed, and injected with the same dosage and regimen of 4-hydroxytamoxifen. For multicolor lineage tracing, two doses of 7.5 mg/ml tamoxifen by oral gavage were given two days apart from a 50 mg/ml concentration stock solution dissolved in sterile corn oil and ethanol (10%).

### Nuclei isolation protocol for brain tissue

Mouse cortex was removed from the brain of a PBS perfused animal prior to snap freezing and storage at -80°C until nuclei isolation. Nuclei isolation was performed as described^20^.

### CitraPrep nuclei isolation for adult small intestine

Mouse duodenal tissue was dissected from the junction of the stomach to approximately 3cm distal at the turn of the intestine, flushed with PBS to remove luminal contents, and snap frozen in liquid nitrogen.

Snap frozen tissues were stored at -80°C until nuclei isolation. Glass dounce homogenizers were placed on ice (Wheaton Science Products, 357542). We tested over a dozen iterations of extraction methods (dounced, crushed, homogenized), incubation times (5 minutes and 7 minutes), washing conditions (single, double, or triple washed), wash volumes (5mL, 10mL, and 50mL), chelating agent concentrations (10mM, 25mM, 30mM, and 50mM citric acid), RNAse inhibitor concentrations (0.2 U/uL, 0.5 U/uL, and 1 U/uL), and RNAse inhibitor enzyme mixes (SUPERase, RNasin, Protector, and in varying combination). We scored each condition by 28S:18S ribosomal RNA band ratios and percent ambient RNA, naming the top condition *CitraPrep* (**Extended Data Fig. 1**). For *CitraPrep,* frozen tissue was placed into a petri dish on ice and cut into 0.2cm cubed pieces or smaller using clean scissors and forceps in 1mL of CitraHB (0.25M sucrose, 25mM KCl, 5mM MgCl_2_, 20mM Tricine-KOH pH 7.8, 10mM citric acid, 0.5U/uL RNasin (Promega, N2615), 0.5U/uL SUPERase RNase inhibitor (Thermo Fisher, AM2696), 1mM DTT (Sigma, D0632), 0.15mM spermine tetrahydrochloride (Sigma, S1141), 0.5mM spermidine trihydrochloride (Sigma, S2501), and cOmplete, EDTA-free protease inhibitor (Sigma, 11836170001)). Chopped tissue (totaling less than 1.5cm cubed) was transferred to the glass dounce homogenizer on ice and 4mL of CitraHB was added. Tissue was incubated on ice for 7 minutes. Tissue was homogenized 20x with loose pestle before adding 320uL of NP-40 and homogenizing 40x with tight pestle. If tissue does not homogenize, an electric homogenizer with short bursts may be used. Nuclei were observed under the microscope for quality control before proceeding.

Tissue lysate was filtered through a 40um strainer into a 50mL conical tube. Next, 5mL of 50% iodixanol solution (5 volumes OptiPrep and 1 volume of Diluent composed of 150mM KCl, 30mM MgCl_2_, 120mM Tricine-KOH pH 7.8 mixed, plus 10mM citric acid, 0.5U/uL RNasin, 0.5U/uL SUPERase RNase inhibitor, 1mM DTT, 0.15mM spermine, and 0.5mM spermidine) was added to the filtered tissue lysate to make a 25% iodixanol solution and vortexed to mix. The tissue lysate was slowly underlaid with 30% iodixanol solution (30% OptiPrep in CitraHB with 0.5U/uL RNasin, 0.5U/uL SUPERase RNase inhibitor, 1mM DTT, 0.15mM spermine, and 0.5mM spermidine), then slowly underlaid with 40% iodixanol solution (40% OptiPrep in CitraHB with 0.5U/uL RNasin, 0.5U/uL SUPERase RNase inhibitor, 1mM DTT, 0.15mM spermine, and 0.5mM spermidine). Without disturbing the layers, the 50mL tube was weighed and balanced before centrifugation at 10,000g for 18 minutes at 4°C with no brake.

Nuclei were collected from the interface layer with a 1000mL pipette and placed into a new tube with 10mL nuclei wash buffer (1x PBS, 1% BSA, 0.5U/uL RNasin, and 0.5U/uL SUPERase RNase inhibitor) and vortexed before filtering through a 20um filter. Nuclei were pelleted by centrifugation at 500g for 5 minutes at 4°C. Supernatant was removed and tissue was washed a second time. The pellet was resuspended in nuclei wash buffer and nuclei were counted using a hemocytometer.

### Analysis of RNA integrity

RNA isolation was performed as described^97^. RNA quality of samples prepped by CitraPrep or established methods^1,20,21^. was checked using RNA formaldehyde electrophoresis to assess ribosomal RNA band integrity. 1% agarose was prepared by heating water and adding 1x MOPS (Quality Biological, 351-059-101) and 2% formaldehyde in a chemical hood before pouring into a gel unit to solidify. 1-2ug of RNA was processed by adding 10uL of RNA loading dye (60 uL 10x MOPS, 120 uL 37% formaldehyde, 300 uL formamide, 30 uL bromophenol blue dye, 10 uL GelRed) and heating at 95 degrees for 2 minutes to denature RNA before immediate transfer to ice to prevent formaldehyde from evaporating. The gel was loaded and ran at 50-60 V in 1x MOPS buffer in water. The gel was visualized using GelRed with the BioRad GelDoc EZ Imager and the ratio of 28S and 18S bands were determined using ImageJ. Analysis of RNA integrity for MERFISH experiments was done by assessing the DV200 from several sections before and after tissue collection using High Sensitivity RNA TapeStation from Agilent Technologies. We assessed the cDNA size and quantity using High Sensitivity DNA ScreenTape with D5000 on an Agilent TapeStation instrument. Traces with below 400bp average sizes were deemed too degraded with low integrity RNA input, and samples below this threshold were excluded from further analysis.

### Single-nucleus RNA-sequencing

Single-nucleus suspensions for each sample was loaded into a separate well of a Chromium 10X Genomics single cell 3’ library chip as per the manufacturer’s protocol with a modified fragmentation time of 4 minutes (10X Genomics: 3’ GEM Library and Gel Bead Kit v3.1 1000128, Chromium Next GEM Chip G Single Cell Kit 1000127), aiming to recover 15,000 nuclei. All libraries were sequenced paired-end following 10X Genomics guidelines on an Illumina NovaSEQ at the University of Chicago Genomics Facility or at the Case Western Reserve University School of Medicine Genomics Core, aiming to sequence 50,000 reads per nuclei.

### Processing single-nucleus RNA-sequencing

Sequencing data were aligned and counted using CellRanger v7.1.0 followed by CellBender v3 with the number of epochs, fpr, and learning rate optimized for each individual dataset based off ELBO and UMI count curves^98^. This processing step uses a maximum-likelihood inference algorithm that filters ambient RNAs and barcode-swapped reads from the raw count matrix outputted from CellRanger. The filtered dataset was then processed with an nUMI cutoff of >500 and <8000. Nuclei with fewer than 5% mitochondrial gene contamination were retained, albeit nearly all nuclei prepped with *CitraPrep* fell within this boundary. Prior to normalization, cells expressing >2% of *Kcnq1ot1* transcripts, a previously identified marker of low quality cells were removed from the analysis^99^. A total of 111,920 nuclei passed quality control filtering with a mean and median detection of 1763 and 1439 genes per nucleus and a mean and median detection of 7249 and 4647 UMIs per nucleus.

We next used DoubletFinder to selectively remove heterotypic doublets^1,100,101^. First, we generated artificial doublets from existing snRNAseq data before pre-processing the merged artificial-real dataset. We used the PC distance matrix to find the cell’s proportion of artificial k nearest neighbors to then rank and threshold according to the expected number of doublets with pANN threshold set to 0.10 and artificial doublets at 0.25. The 8 mouse datasets were integrated using SCTransform normalization with variable regression set to mitochondrial reads using Seurat 4.0^29–32^ package followed by principal component analysis prior to clustering. We mathematically defined the optimal principal components (PC) to use by calculating the percent variation associated with each. Next, we calculated the cumulative percentage for each PC and determined which exhibits greater than 90% cumulative percent and less than 5% variation associated with the PC. Next the difference between the variation of PC and subsequent PC was calculated where change of percentage of variation is more than 0.1% before finding the smaller of the two values as the optimal value for downstream analyses. Batch effects were removed using Harmony^102^ by grouping variables by original sample identity. Resolution was initially set at 0.5-1 before adjusting until all unsupervised clusters represented transcriptionally unique groups as determined by post-hoc analysis of top 10 enriched transcripts per cluster on heatmap. Clustering was visualized by UMAP and clusters were annotated by inputting the top 50 DEGs per cluster into CellMeSH for probabilistic cell type identification through indexed literature^103^ and further cross referenced to literature. Finally, clusters with more than 50% of the top 50 DEGs associated with mitochondrial or ribosomal RNA were removed as this indicates degradation of RNA. Two clusters (17 and 19) were removed from the final analyses leaving 107, 001 total nuclei.

### Western blotting

Protein lysates from whole duodenum, cortex, and colon were prepared in RIPA lysis buffer (Sigma, R0278) supplemented with cOmplete, EDTA-free Protease inhibitor (Sigma, 11836170001). Protein concentrations were measured using the BCA kit (Thermo Fisher, 23225) and then diluted in 4x Laemmeli buffer (40% glycerol, 8% SDS, 240mM Tris-HCl pH 6.8, 5% b-mercaptoethanol, 12.5mM EDTA, 0.04% bromophenol blue). Lysate was resolved on 4-12% NuPAGE 1.5mm Bis-Tris gel (Thermo Fisher, NP0335BOX) and transferred to PVDF membrane (Thermo Fisher, LC2002). Membranes were blocked with 5% milk in Tris-buffered saline with 1% Tween 20 (TBST) for 1hr, followed by overnight incubation at 4°C with primary antibodies (PDE3A from ProSci 18-159 at 1:500, CDH10 from Novus NBP2-92600 at 1:1000, GHR from Affinity Biotech DF8425 at 1:1000, SORBS2 from Bioss BS-4905R at 1:500, B-ACTIN from Sigma A3854 at 1:10,000). Membranes were washed three times with TBST before incubation for 1hr with either anti-rabbit IgG conjugated to horseradish peroxidase (HRP) or anti-mouse IgG-HRP.

### Tissue sectioning and immunostaining

For sectioning, mouse duodenal tissue was dissected from the junction of the stomach to approximately 3cm distal at the turn of the intestine, flushed with PBS to remove luminal contents, and fixed overnight at 4 °C in 4% paraformaldehyde (PFA). Fixed tissues were washed 3 times in PBS before moving to 30% sucrose in PBS and stored at 4 °C overnight until tissue sank. Tissues were thoroughly wiped clean of sucrose using a Kim wipe, cut into smaller segments, and washed several times in Optimal Cutting Temperature Compound (OCT; Sakura, 25608-930) prior to embedding. Tissue blocks were sections on a Leica CM1950 Cryostat at 15-20 um onto slides and stored at -80 °C.

For immunofluorescent staining, slides were blocked for 30 minutes room temperature with PBS+0.1% Triton X-100 (VWR) with 5% donkey serum (Jackson ImmunoResearch) then incubated overnight at 4°C in primary antibodies including RFP (Chromotek, 5f8-100), GFAP (Agilent DAKO, Z033429-2 and Thermo Fisher 13-0300), GFP (Aves, GFP-1020), NFIA (Sigma, HPA006111), VIM (Biolegend, 919101), B-CATENIN (Thermo Fisher, 71-2700), FIGN (MyBioSource, MBS7048173), NPAS2 (GeneTex, GTX105741), PIEZO2 (Novus, NBP1-78624; Novus NBP1-78538), SOX10 (R&D, AF2864), NRP1 (R&D, AF3870), GRIK3 (Thermo Fisher, MA5-31743), SOX6 (Abcam, ab30455), and PPARd (Thermo Fisher, PA1-823A). Slides were washed, incubated with secondary antibody for 1 hour at room temperature, and washed. All secondary antibodies were purchased from Jackson ImmunoResearch. Slides were mounted in Fluoromount G (SouthernBiotech), covered with coverslips, and sealed with nail polish. Nuclei were stained with DAPI (Invitrogen). Human slides embedded in paraffin were deparaffinized prior to immunostaining by incubating in xylene two times, 1:1 xylene:100% ethanol once, 100% ethanol two times, followed by one time in 95% ethanol, 70% ethanol, and 50% ethanol before washing in running cold water to rehydrate.

### MERFISH library design

To profile gene expression across gastrointestinal tissues and compare colon and duodenal enteric glial cell transcripts, we selected a panel of 140 genes. These genes were included due to differential expression across tissues (24 enriched in colon, 50 in duodenum, 18 in ileum), between enteric glial subtypes identified by snRNAseq (26 enriched in EGC1, 8 enriched in EGC2, 40 enriched in EGC3, 13 in EGC4, 6 in both EGC5 and EGC6, and 29 enriched in EGC6 cells), between spatial transcriptomic clusters (21 enriched in cluster 0, 20 in cluster 1, 1 in cluster 2, 4 in cluster 3, 5 in cluster 4, and 14 in cluster 5), alongside 5 canonical enteric glial markers and 3 canonical pan-enteric neuron markers. MERFISH encoding probes targeting these 140 genes were designed through Vizgen, with all transcripts falling within single gene abundance threshold.

### MERFISH tissue collection and cryosectioning

Upon euthanasia of 6 week old mice, the duodenum and colon were quickly collected, placed on ice in cold 4% v/v paraformaldehyde (PFA; Electron Microscopy Sciences, 15714) in 1x PBS, and flushed immediately with this solution to remove luminal contents. We initially sought to optimize duodenal and colonic Swiss roll preparations for higher tissue coverage, however these attempts were unsuccessful in achieving quality MERFISH data for analysis, thus we focused on cross sections for MERFISH experiments. Tissues were then processed using a described approach^104^. Tissues were incubated for 3 hours in 4% PFA in 1x PBS at 4°C before transferring to 30% w/v sucrose in 4% v/v PFA in 1x PBS at 4°C for 16 hours. Tissues were washed in OCT, placed into RNAse free plastic cryomolds, and immediately frozen in dry ice. Tissues were transferred to - 80°C for storage until further processing. All buffers and materials were prepared with RNAse-free reagents, including disinfection with RNAse decontamination solution and ethanol, and were pre-chilled.

Tissues were equilibrated to -20°C in a Leica CM1950 cryostat for 20 minutes prior to mounting for sectioning. MERSCOPE frozen tissue slides were equilibrated to room temperature in parallel. Tissues were sectioned to trim away flanking tissue, collecting at least 3 tissues before and after into pre-chilled tubes for RNA isolation to measure DV200. To collect tissue onto MERSCOPE slides (Vizgen PN10500001), we sectioned at 10 μm directly onto the room temperature MERSCOPE slide, holding the slide for 10 seconds for adhesion. The MERSCOPE slide with tissue was placed into a clean 60-mm petri dish and placed in the cryostat for 30 minutes for further adhesion. Slides were washed 3x with RNAse free 1x PBS, dried for 1 hour at room temperature, then overlaid with 70% ethanol to permeabilize the tissues. Dishes were secured with parafilm and placed at 4°C until further processing.

### MERFISH sample preparation

The parafilm-sealed dish holding the MERSCOPE slide in 70% ethanol was placed in the MERSCOPE Photobleacher (Vizgen PN10100003) for 5 hours at room temperature for autofluorescence quenching. Following quenching, slides were washed once in 1x PBS before they underwent cell boundary staining. Tissues were blocked at room temperature for 1 hour (Vizgen PN20300100) before primary staining (Vizgen PN20300010) with 1U/uL RNAse inhibitor for 1 hour at room temperature. Finally, tissues were stained with secondary staining solution (Vizgen PN20300011) with 1U/uL RNAse inhibitor for 1 hour at room temperature before washing 3x with 1x PBS and fixation at room temperature for 15 minutes. We next washed in PBS, washed with wash buffer (Vizgen PN20300001), and incubated in formamide wash buffer (Vizgen PN20300002) for 30 minutes at 37°C. After thorough removal of the formamide, MERSCOPE custom gene probes were spotted onto the tissue sections and overlaid with parafilm. Dishes were sealed and incubated at 37°C for 36 hours.

Post encoding probe hybridization, tissues were washed and prepared for gel embedding. First, probe mix was removed and replaced with 5mL of formamide wash buffer to incubate at 47°C for 30 minutes for two iterations. Slides were washed in wash buffer. During the second formamide incubation, gel coverslips (Vizgen, PN30200004) were cleaned with RNAse decontamination spray and ethanol before pre-treating with Gel Slick solution for 10 minutes. After washing the MERSCOPE slide with wash buffer, a gel embedding premix (Vizgen PN20300004) was prepared with 0.5% w/v ammonium persulfate solution and 0.05% N,N,N,N’-tetramethylethylenediamine and the slide was rinsed once in this solution before adding 50uL to the tissue section and covering with the Gel Slick-treated coverslip to evenly embed tissues into the gel. Gel was formed at room temperature for 1.5 hours before removing the coverslip and proceeding with tissue digestion with

1.5U/uL RNAse inhibitor (SUPERasin and Protector) in digestion premix (Vizgen PN20300005) at 37°C for 5 hours. Following digestion, tissues were cleared using SDS based clearing premix (Vizgen PN20300003) and 4U/mL Proteinase K (NEB, P8107) for 15 hours at 37°C before refreshing the solution and waiting 10 hours before switching to clearing premix without Proteinase K and storage at 37°C until imaging.

### MERFISH and Visium spatial transcriptomics analysis

MERFISH imaging was completed using Vizgen MERSCOPE Ultra Spatial Imaging platform. Imaging cartridge was thawed at 37°C for 60 minutes, wiped down to dry before cleaning with RNAse decontamination solution and ethanol. Clearing solution was removed from the tissue slides before washing twice with sample wash buffer. DAPI and PolyT staining was then done for 15 minutes rocking at room temperature protected from light, followed by a 10 minute incubation at room temperature with formamide wash buffer and a final wash in sample prep wash buffer. The MERSCOPE instrument fluidics were washed immediately preceding imaging. MERSCOPE slides were loaded into a clean flow chamber before chamber priming, tissue overview, high magnification imaging, and data acquisition. Following data acquisition, segmentation parameters were selected and images were processed as output. We used a modified Seurat code to load data into R to support parquet files for cell boundary segmentation. Each individual animal and tissue region was processed, normalized using SCTransform, followed by principal component analysis prior to clustering. We mathematically defined the optimal principal components (PC) to use by calculating the percent variation associated with each. Next, we calculated the cumulative percentage for each PC and determined which exhibits greater than 90% cumulative percent and less than 5% variation associated with the PC. Next the difference between the variation of PC and subsequent PC was calculated where change of percentage of variation is more than 0.1% before finding the smaller of the two values as the optimal value for downstream analyses. Individual regions were combined by merging, and batch effects were removed using Harmony^102^ by grouping variables by original sample identity. Resolution was set at 0.1. Clustering was visualized by UMAP and differential gene expression was performed using Wilcoxon Rank Sum test. Visium spatial transcriptomics dataset was analyzed using the standard Visium HD Seurat pipeline. Briefly, data was loaded, normalized, and variable features were determined prior to clustering. Enteric glia were identified via unsupervised clustering and expression of a cohort of glial cell markers before subclustering enteric glia alone to assess spatial distribution and marker expression.

### Whole mount immunofluorescence

Longitudinal muscle and myenteric plexus tissue was isolated as described above and washed one time in PBS before fixing in 4% PFA for 20 minutes at room temperature. Tissue was washed 3 times in PBS before blocking for 30 minutes room temperature with PBS+0.1% Triton X-100 (VWR) with 5% donkey serum (Jackson ImmunoResearch) and incubating overnight at 4°C in primary antibodies including RFP (Chromotek, 5f8-100), GFAP (Agilent DAKO, Z033429-2 and Thermo Fisher 13-0300), PIEZO2 (Novus, NBP1-78624; Novus NBP1-78538), and SOX10 (R&D, AF2864). Tissues were washed, incubated with secondary antibody for 1 hour at room temperature, and washed. All secondary antibodies were purchased from Jackson ImmunoResearch. Nuclei were stained with DAPI (Invitrogen). Imaging was performed on two microscopes: (1) a Leica TCS SP8 gated super-resolution STED confocal laser scanner mounted on an inverted DMI6000 microscope and (2) an inverted Leica DMi8 microscope.

### Longitudinal muscle myenteric plexus preparations

Mouse duodenal tissue was dissected from the junction of the stomach to approximately 3cm distal at the turn of the intestine, flushed with PBS to remove luminal contents, and placed in PBS on ice. Mesentery and pancreatic tissue were removed, then duodenal tissue was cut into 1.0-1.5cm segments. Tissue segments were individually slid onto a borosilicate glass Pasteur pipette and, using pressure from a finger for stability and under microscopy, forceps were used to create a thin nick along the serosa and muscle layers of the duodenum. A cotton swap was wetted with cold PBS and used to wipe along the nick while slowly turning the tissue using the stabilizing finger to peel away the longitudinal muscle and myenteric plexus layer. This region, when done properly, resembles the transparent membrane of an onion. Collected tissue was immediately used for subsequent assays including whole mount immunostaining, mouse primary cell culture, *ex vivo* contractility assays, and calcium imaging.

### Mouse primary cell culture

Longitudinal muscle myenteric plexus layers were collected and placed in a digestive KREBS solution buffer with 1.3 mg/mL collagenase type II (Worthington, LS004176) and 0.3 mg/mL BSA (Sigma, A4161) through modification of established protocols^105,106^. KREBS solution was composed of the following: 121 mM NaCl, 5.9 mM KCl, 2.5 mM CaCl_2_, 1.2 mM MgSO_4_, 1.2 mM NaH_2_PO_4_, 10 mM HEPES, 21.2 mM NaHCO_3_, 1 mM pyruvic acid, 8 mM glucose. All tissues must be collected and placed into the solution on ice within 30 minutes of dissection. The digestive KREBS solution was moved to 37 °C and bubbled with carbogen for 1 hour. Following this hour, solution was quenched with base media and centrifuged at 356 g for 10 minutes at 4 °C. Enteric glial cell media was composed of 1:1 DMEM/F12 to Neurobasal, 1x B27 (ThermoFisher, 17504044), 1x N2max (Fisher, AR009), 10ng/mL GDNF (Stem Cell Technologies, 78058), 20ng/mL FGF2 (R&D Systems, 23-3FB-010M), and 1x penicillin/streptomycin (Thermo Fisher, 15070-063). The remaining pellet was gently resuspended in enteric glial cell media supplemented with 1x CloneR (Stem Cell Technologies, 5889) and 50ug/mL primocin (Invivogen, ant-pm-1), filtered through 100 um filters (Fisher Scientific, 08-771-19), and plated onto 10cm 0.1mg/mL poly-ornithine primed, 10μg/mL laminin coated plates. Media was replaced every other day and cells were split into 96-well plates after reaching confluence or 2 weeks in culture.

For analysis, cells were washed one time in PBS, fixed for 20 minutes in 4% PFA at room temperature, and washed 3 times in PBS prior to blocking. Cells were blocked for 30 minutes room temperature with PBS+0.1% Triton X-100 (VWR) with 5% donkey serum (Jackson ImmunoResearch) then incubated overnight at 4°C in primary antibodies including RFP (Chromotek, 5f8-100), GFAP (Agilent DAKO, Z033429-2 and Thermo Fisher 13-0300), GFP (Aves, GFP-1020), NFIA (Sigma, HPA006111), VIM (Biolegend, 919101), B-CATENIN (Thermo Fisher, 71-2700), FIGN (MyBioSource, MBS7048173), NPAS2 (GeneTex, GTX105741), PIEZO2 (Novus, NBP1-78624; Novus NBP1-78538), SOX10 (R&D, AF2864), NRP1 (R&D, AF3870), GRIK3 (Thermo Fisher, MA5-31743), SOX6 (Abcam, ab30455), and PPARd (Thermo Fisher, PA1-823A). Plates were washed, incubated with secondary antibody for 1 hour at room temperature, and washed. All secondary antibodies were purchased from Jackson ImmunoResearch. Nuclei were stained with DAPI (Invitrogen). Brightfield imaging of primary enteric glial cells was performed on a Leica DMIL LED microscope.

### Quantitative PCR

Total RNA was isolated by phenol-chloroform extraction using TRIzol (Ambion). Genomic DNA was depleted by DNAse treatment (Invitrogen) before reverse transcription using iScript cDNA Synthesis kit (1708891, BioRad). Real-time PCR was performed using TaqMan Fast Advanced qPCR mix (Thermo Fisher, 4444963) in technical quadruplicates using the Applied Biosystems 7300 real-time PCR system and analyzed by computing ΔCq. Genes were normalized to *Hprt1*.

### Biochip fabrication

Biochip microchannel geometry was formed via a laser micro-machined, double-sided adhesive film (3M, 8412KCL) and polymethyl methacrylate (McMaster-Carr, 8560K) components assembled in the following manner: a 1 mm polymethylmethacrylate spacer was sandwiched between two double-sided adhesive films and adhered to a glass slide for contraction imaging or to a cover slip for calcium imaging. At this point, microchannels were rinsed with ethanol and deionized water, then functionalized with N-γ-maleimidobutyryl-oxysuccinimide ester (Fisher Scientific, PI22309) to facilitate Matrigel bonding. For *ex vivo* preparations, we chilled biochips before adding 50 µL of Matrigel solution to each channel. Dissected tissues were carefully laid flat in each channel to facilitate adhesion before incubation at 37 °C with 5% CO_2_ for 10 minutes for curing. Culture media was then gently added to each channel and biochips were incubated at 37 °C with 5% CO_2_ for at least 60 minutes for tissue recovery. Thereafter, a 3.175 mm polymethylmethacrylate block, with inlet and outlet holes was bonded on top of the open biochips to enclose each channel. Assembled microchannels were then perfused with fresh modified KREBS buffer as described above to remove any residual air bubbles, then kept at 37 °C with 5% CO_2_ until use.

### Calcium imaging and quantification in intact tissue

The longitudinal muscle myenteric plexus layer of Ai95(RCL-GCaMP6f)-D mice were prepared as described above. Fluorescent images of the mounted tissues and cells were acquired via an inverted Olympus IX83 microscope at 40X objective. Acquisition was completed with the GFP shutter open acquiring 408 images over a 2 minute period.

Images were analyzed using Fiji and ImageJ plugin Time Series Analyzer ver3. Briefly, the GCamP6f expressed cell regions were defined as regions of interest (ROI) and circled around then obtained the pixel intensity with Time Series Analyzer. Background nonspecific muscle staining/autofluorescence was subtracted from each picture by using the defined region without GCamP6f expression. The sequence of the experiments was (1) background activity (2) with continuous flow at 1 dyn/cm^2^ (rates reported to activate PIEZO2 in other cell types^76^) (3) with Piezo2 inhibitor D-GsMTx4 at 10 μM plus continuous flow (4) washout. The same ROIs were tracked over time to record the response of individual cells over time. XY drifts between each experiment were manually corrected so the ROI targeted the same cells throughout the sequence of experiments.

### Calcium imaging and quantification in dispersed enteric glial cells

Primary cultures were generated as described above. Enteric glial cells were split from poly-ornithine / laminin coated plates onto functionalized intestine-on-a-chip devices modified onto cover slips with no lid attached to allow for cell growth. Immediately before imaging, lids were added to the functionalized chips with cells with fluidics lines attached for shear stress application and media was changed from EGC growth media as described above to KREBS buffer with 1 μM Fluo4, AM (Thermo Fisher, F14201) with 0.02% Pluronic F-127 to disperse the nonpolar AM ester in aqueous media. Enteric glia were incubated in this media for at least 30 minutes at 37°C protected from light before imaging via an inverted Olympus IX83 microscope at 20X objective. Acquisition was completed with the GFP shutter open acquiring 1016 images over 5 minutes.

In contrast to experiments using intact tissue, cells were imaged for 120 seconds at baseline with no stimulation. After 60 seconds and before 120 seconds, the microfluidic clip was removed to allow fluidics to pass through. In some experiments the removal of this clip produced a brief pulse of shear stress that initiated a small early response. At 120 seconds, 10 seconds of continuous shear stress at 1 dyn/cm^2^ was applied before returning to 0 dyn/cm^2^ for the remaining 170 seconds. We binned responses into three groups: baseline (first 120 seconds), stimulation (90 seconds from onset of flow beyond), and recovery (final 90 seconds). Images were analyzed using Fiji and ImageJ plugin Time Series Analyzer ver3. Briefly, the GCamP6f expressed cell regions were defined as regions of interest (ROI) and circled around then obtained the pixel intensity with Time Series Analyzer. Background nonspecific muscle staining/ autofluorescence was subtracted from each picture by using the defined region without GCamP6f expression. We calculated the area under the curve from the ι1F/F in each phase for quantitative analyses. Location of each measurement was annotated for follow up analysis.

After recovery in EGC growth media at 37°C for at least 30 minutes, cells were incubated in KREBS buffer with 1 μM Fluo4, AM (Thermo Fisher, F14201), 0.02% Pluronic F-127, and 10 μM D-GsMTx4 for 30 minutes. Chips were then loaded onto the microscope with a syringe containing KREBS + 10 μM D-GsMTx4 for shear stress stimulation following the same paradigm as described above in the same location as previously analyzed. The same ROIs were tracked over time to record the response of individual cells over time. XY drifts between each experiment were manually corrected so the ROI targeted the same cells throughout the sequence of experiments. Following inhibition analysis, cells were fixed for 20 minutes at room temperature with 4% PFA for immunostaining.

### Lineage tracing of Cre efficiency

For Cre tracing studies to assess targeted cells, 4-hydroxytamoxifen was delivered to *RCL-*YFP*; Gfap-*cre^ERt2^*, RCL-*YFP*; Sox10-*cre^ERt2^, and *Plp1-*cre^ERt2^ mice on postnatal day 21 (3 weeks) at 75 mg/kg by intraperitoneal injection once daily for 5 days and collected 3 weeks after first injection for analysis. Intestinal tissue was collected, fixed in 4% PFA overnight at 4°C, and sectioned for immunostaining for Cre (Cell Signaling, 15036), GFP (Abcam, ab13970), GFAP (DAKO, Z033429-2), and DAPI. Sectioned tissues were imaged on a Leica TCS SP8 gated super-resolution STED confocal laser scanner mounted on an inverted DMI6000 microscope, and targeted cells were quantified by counting cells stained for each marker as a percentage of total cells.

### RNAscope in situ hybridization (ISH)

ISH was performed using the RNAscope Multiplex Fluorescence V2 Assay (ACDBio, cat# 323110) following the manufacturer’s instructions for frozen fixed tissue. The tissue was dehydrated using ethanol and blocked with hydrogen peroxide. Target retrieval was undertaken for 5 minutes and tissue was dried before applying the supplied protease III (ACDBio, cat# 322340). Tissue was then incubated with RNA probes from ACDBio at 40°C for 2 hours. The tissue was placed in a 5x Saline Sodium Citrate buffer overnight and washed the next morning with RNAscope Wash Buffer (ACDBio, cat# 310091). After a series of amplification steps, RNA was hybridized with opal dye fluorophores (Akoya Biosciences). ISH images were captured using a Leica DMi8 fluorescence microscope and Leica DFC7000 T camera at 20x magnification. Mouse-specific RNA probes used were *Piezo2* (ACDBio, cat# 439971) and *GFAP* (ACDBio, cat # 313211-C3).

### Whole gut transit time assay

To measure the amount of time it took contents to fully pass through the gastrointestinal tract, we gavaged mice with 6% carmine red (Sigma, C1022) in methylcellulose (Sigma, 274429). To prepare this reagent, we dissolved 0.5% methylcellulose in hot distilled water prior to mixing 6% carmine red. The solution was used at room temperature but stored at 4°C until use.

Oral gavage of 300uL carmine red solution was performed using a 24-20 gauge feeding needle between 2.5-3.8 cm in length. Before the oral gavage procedure, we measure the distance from the oral cavity to the caudal point of the sternum with the feeding needle on the outside of the restrained animal to mark the distance needed to insert into the esophagus. This distance was marked using a marker. The syringe was loaded with the volume to be administered (10 uL/gram mouse weight). The outside of the needle was wiped to remove any compound on the outside.

Mice were gently scruffed and restrained in an upright position to immobilize the head and neck. The gavage needle was slid into the left side of the animals’ mouth behind the teeth in front of the first molar along the roof of the animals’ mouth. The gavage needle was used to gently tilt the mouse’s head back towards the spine with gentle pressure to allow a straight line from the mouth to the esophagus and stomach, ensuring no resistance. The gavage was passed until the pre-marked line is reached. The solution was slowly injected over the course of 2-3 seconds to minimize fluid coming back up the esophagus.

Mice were housed in upside down empty p1000 pipette tip boxes on metal racks to suspend mice above a clean white paper mat. Mice were not disturbed during the assay other than during the gavage procedure to minimize effects due to handling^107^. A timer was started after gavaging mice and fecal matter was observed immediately upon dropping to the mat for carmine red color. The time from gavage to the first carmine red defecation was documented. All mice were tested with littermate controls also injected with tamoxifen; two litters of mice were used for analyses with a total of 4 control and 14 Piezo2^fl/fl^; *Sox10*-cre^ERt2^ mice. Mouse genotypes were blinded for data collection. There were no data exclusions.

### Gastrointestinal transit time assay

Gastrointestinal motility was analyzed to determine the contribution of enteric glial subtypes to regulation of intestinal physiology. We performed pilot experiments to determine the optimal fasting time prior to starting the assay, in which mice were fasted for different lengths of time and autofluorescence of stomach contents were measured. Mice were fasted for 2-5 hours prior to oral gavage of a 70-kDa FITC labeled dextran solution (100 µL, 25 mg/mL PBS; gavage procedure as described above). Initially, we collected timepoints between 20 and 50 minutes after gavage to determine when to assay experimental mice. All data in the manuscript were collected 30 minutes post gavage.

The gastrointestinal tract was removed from stomach to anus and placed into ice cold PBS to cold shock tissue and inhibit further peristalsis. We carefully cut the mesenteric and pancreatic tissue to straighten the gut and measured the small intestine length from the pyloric sphincter to the cecum. The small intestine was divided into 10 equal sections and each segment, in addition to the stomach and the cecum, was flushed with 2 mL of PBS. The colon tissue was measured and divided into 3 equal segments and flushed with 2mL of PBS. The flushed contents were centrifuged at 500 rpm for 10 minutes and 200 ul of the supernatant from each section was placed in a 96 well plate. The quantification of the fluorescent signal in the supernatant from each segment was determined utilizing a Synergy neo 2 multi-well fluorescence plate reader from BioTek/Agilent (excitation 545 nm and emission 590 nm). The distribution of the fluorescent signal in the intestinal segments was mapped and used to calculate the rate of gastric emptying and the geometric center of fluorescence (GCF). GCF was determined by calculating the fraction of fluorescence per segment multiplied by the segment number (1-15) and adding all segments together before dividing by 15. This was then multiplied by 10, with GCF ranging from 1 to 10 with a higher number indicating a faster motility and shorter intestinal transit time. Gastric emptying was calculated as: [(total fluorescence – fluorescence in stomach)/(total fluorescence)] ×100. All mice were tested with littermate controls also injected with tamoxifen with multiple litters included. Mouse genotypes were blinded for data collection.

### Fecal matter composition

Fecal water content was measured by weighing feces in a 1.5 mL tube with the cap on immediately after collection from the animal before opening the cap and heat drying the sample at 42 °C for 48 hours. After 48 hours, sample was capped and weighed before measuring the weight of the tube. Percent water content was calculated as (pre drying weight – tube weight) - (post drying weight – tube weight) / (pre drying weight – tube weight) * 100. All mice were tested with littermate controls also injected with tamoxifen with multiple litters included. Mouse genotypes were blinded for data collection. There were no data exclusions.

### Epithelial barrier integrity assays

Mice were fasted for 2-5 hours prior to oral gavage of 4-kDa (Sigma-Aldrich, 600 mg/kg body weight, 80 mg/mL PBS) by gavage. The 4-kDa fluorescent dextran is a non-digestible dextran conjugated with dye that can transit through the gastrointestinal tract and passively cross the intestinal epithelium. The use of this dye allows for tracking of peristalsis and gut motility while also allowing for a readout of epithelial barrier integrity. All data in the manuscript were collected 30 minutes post gavage.

Mice were euthanized and blood was collected directly from the heart to ensure contents were not diluted by intraperitoneal fluids. Plasma was collected by centrifugation at 2000g for 5 minutes. Equal volumes of serum were loaded into a 96-well microplate in duplicate and read by spectrophoto fluorometry with an excitation of 485nm and emission of 528nm using as standard serially diluted FITC-dextran.

### *Ex vivo* motility assays

Longitudinal muscle myenteric plexus preparations from 6 week old mice were dissected fresh and washed one time in cold Matrigel before placing in 50uL of a 50% KREBS buffer: Matrigel solution in channels of chilled biochips and processed as described above. Biochips were incubated at 37 °C with 5% CO_2_ for 1 hour prior to analysis to allow tissue to recover. Tissues were screened for quality and only tissues exhibiting spontaneous contractions were used for contractility assays under stimulation. Images were taken every 250 ms via a BX51WI Olympus microscope equipped with Retiga EXi CCDcamera (QImaging) using microManager and ImageJ applications. Acquisition spanned for 30 seconds in brightfield at baseline and with 1 dyn/cm^2^ flow (above threshold) or at baseline.

### Total neurotransmitter levels from the tunica muscularis

Levels of acetylcholine and nitric oxide were measured from primary mouse tissues. The longitudinal muscle myenteric plexus was dissected from the proximal small intestine as described above and immediately homogenized in RIPA lysis buffer (Sigma, R0278) with cOmplete, EDTA-free Protease inhibitor (Sigma, 11836170001). The protein content was measured using a BCA assay (Thermo Fisher, 23225) to normalize neurotransmitter levels to protein content of the tissue samples. The level of acetylcholine was determined using the Choline/Acetylcholine Quantification Kit (Sigma, MAK056) and measuring the difference between the level of total choline and free choline. Total nitrite/nitrate was measured as a proxy for nitric oxide concentration using the Nitric Oxide Assay Kit (Abcam, ab65327), with 5 hours of incubation with the enzyme cofactor to permit more than 99% conversion of nitric oxide to nitrite.

### Cell stretching bioreactor assay

Enteric glial cells were cultured as described above. Cells were plated into poly-ornithine / laminin coated BioFlex culture plates (Flexcell, BF-3001U-Case) and allowed to grow to 85% confluency. Media was changed from EGC growth media to KREBS buffer after washing cells gently 3x with pre-warmed KREBS buffer. Enteric glia incubated for 10 minutes in KREBS buffer before collecting conditioned media for a baseline measurement, then changed to fresh KREBS buffer. After 10 minutes of incubation to mirror baseline, bioreactor plates were stretched using 5psi of air for distension and conditioned media was collected 30 seconds later and 10 minutes later for untargeted metabolomics. For nitric oxide, ATP, and LDH assays the same paradigm was used but focused on 30 seconds post-stretch for conditioned media analysis.

### Bulk RNA-sequencing and analysis

The longitudinal muscle myenteric plexus was dissected from the proximal small intestine as described above, washed in cold 1x PBS, and immediately homogenized in TRIzol. Total RNA was isolated by phenol-chloroform extraction. Genomic DNA was depleted by DNAse treatment (Invitrogen) before sending to Novogene for library preparation and sequencing on the Illumina NovaSeq X with paired-end 150bp reads with a read-depth of at least 20 million reads per sample. Reads were mapped to the mm10 genome using kallisto v0.46.1^108^. Normalized expression and differential gene expression were then generated using DESeq2 v1.32.0^109^.

### Untargeted and parallel reaction monitoring metabolomics via LCMS

Two different LCMS experiments were performed on conditioned media from our cell stretching bioreactor assay. The first was an untargeted metabolomics analysis with reverse phase C18 chromatography. The second was to assess specific metabolites including adenosine triphosphate (ATP), acetylcholine (ACh), and 4-aminobutanoate (GABA), determining the relative abundance of these samples across samples.

10mM each of unlabeled ATP, ACh, and GABA was prepared for direct injection into the mass spectrometry (MS) analysis followed by column separation analysis (LCMS) to extract the mass spectra of the compounds. After collection of conditioned media, samples were spun at 10,000g for 5 minutes and 100uL of supernatants were collected, mixed with 600 µL of chilled methanol: water (4:1) containing heavy-labeled internal standards, and vortexed. In addition, 10mM of cocktail of unlabeled ACh and GABA was spiked into a different set of samples. The samples were then centrifuged at 10,000xg for 15 min at 4°C. The supernatants were dried overnight in a speed vacuum and dried extracts were re-suspended in 95:5 water/acetonitrile (%v/v). Each sample was then randomized and subjected to LC-MS analysis. 2 µL from each sample was taken and pooled and this pooled QC sample was analyzed every 10th injection. The internal standards and observed ions are reported in **Supplementary Table 7**. MS2 level data were collected on representative control and treated samples in this study.

Reverse phase chromatography was performed by injecting 6 µL of each sample on a Thermo Accucore Vanquish C18 column with dimensions 100 x 2.1 mm, 1.5 µm particle size (Thermo P/N 27101-102130) at 60°C coupled to a Thermo Vanquish UHPLC by gradient elution where mobile phase A is 0.1% formic acid in water and mobile phase B is 0.1% formic acid in acetonitrile. The Orbitrap Q Exactive HF was operated in positive and negative electrospray ionization modes in different LC-MS runs over a mass range of 56-850 Da using full MS at 120,000 resolution. Data dependent acquisitions (DDA) on the pooled representative QC samples include MS full scans at a resolution of 120,000 and HCD MS/MS scans taken on the top 5 most abundant ions at a resolution of 30,000 with dynamic exclusion. The resolution of the MS2 scans were taken at a stepped NCE energy of 10.0, 20.0 and 30.0.

### LCMS Data Processing and Annotation

Reifycs Analysis Base File (ABF) Converter Software was used to convert all raw data files to ABF file formats. Data were then processed using MSDIAL 1 (v.4.92) for feature detection, identification and alignment using parameters optimized for data acquired on an Orbitrap mass spectrometer. MS1 and MS2 were set to profile mode in both positive and negative ionization modes. Peak detection of MS1 and MS2 spectra were set to tolerances of 0.01 Da and 0.025 Da, respectively, over a mass range of 56-850 m/z with minimum peak width and height of 5 and 1,000,000.

Metabolites identifications were done using publicly available libraries from MassBank of North America (MoNA) containing 13,303 unique compounds (positive mode) 2 and MSDIAL Metabolomics MSP Spectral Kit 3 containing 12,879 unique compounds (negative mode) with 80% identification cut-off score. Before the untargeted metabolomics analysis, three unlabeled standards (ACh, GABA and ATP) were analyzed by either direct infusion injection into the mass spectrometer and/or with prior LC separation. Only mass spectra of ACh and GABA were recovered. Following these analyses, a parallel reaction monitoring (PRM) method was set up to analyze these standards and media samples spiked with or without the standards to extract the MS/MS spectra information. Chromatograms were plotted for two transition for both ACh and GABA and while GABA was not identified in these samples, ACh was positively identified across all the media samples including the KREBS blank sample. The chromatograms for ACh were integrated and the resulting peak areas were normalized to the peak area for two of the heavy label internal standards, Betaine-d9 and Picolinic acid-d4.

Spectral features from MSDIAL processed data were further analyzed via MetaboAnalyst 5.0 (www.metaboanalyst.ca) 4. Zero/missing values were replaced by 1/5 of the minimum peak height over all samples, normalized by sum normalization, log 10 transformed and autoscaled prior to downstream statistical analysis. A raw p-value less than 0.05 was chosen for significance. All p-values were corrected via the Benjamini-Hochberg procedure for false discovery rate (FDR) with a threshold of 0.1. Fold change analysis was performed. Multivariate principal component analysis (PCA) and hierarchical clustering were performed for understanding metabolite variation and expression patterns between groups. Pathway enrichment analysis was performed to identify top 25 altered pathways for putatively identified metabolites in the dataset. In this analysis the Pathway-associated metabolite sets (SMPDB) database, which consists of 99 pathways from the Small Molecule Pathway Database 5 was used to map the putatively identified ions to various pathways.

### Targeted conditioned media assays

Conditioned media from stretch bioreactor assays was spun down at 10,000g for 5 minutes and supernatants were collected for downstream analyses. For nitric oxide assays, a fluorometric assay kit was used to measure nitrate and nitrite in the samples, allowing for total nitric oxide measurement (Abcam, ab65327). Live ATP analyses were both performed using the RealTime-Glo Extracellular ATP Assay kit (Promega, GA5010). For ACh and GABA by LCMS, media was collected from baseline (10 minutes) and stretch (10 minutes rest, 30 seconds after stretch) and conditioned media was centrifuged before supernatant was assessed for ATP. LDH levels were assessed using the LDH-Glo Cytotoxicity Assay (Promega, J2380).

### Enteric neuron and smooth muscle primary cultures

Enteric neuron and smooth muscle primary cultures were performed as described^110^. Briefly, longitudinal muscle myenteric plexus preparations from the small intestine of 6 week old mice were dissected fresh and placed into cold KREBS buffer (121 mM NaCl, 5.9 mM KCl, 2.5 mM CaCl_2_, 1.2 mM MgSO_4_, 1.2 mM NaH_2_PO_4_, 10 mM HEPES, 21.2 mM NaHCO_3_, 1 mM pyruvic acid, 8 mM glucose). Samples were washed two times in KREBS buffer after centrifugation at 4°C for 5 minutes at 350g.

Longitudinal muscle myenteric plexus preparations were resuspended in either neuronal digestion buffer (13mg collagenase II and 3mg BSA in 10mL RPMI1640) or smooth muscle digestion buffer (14mg collagenase II, 3mg Dispase II, 10ug/mL DNase I in 5mL RPMI1640), placed in a petri dish, and chopped with a sterile scalpel before placing at 37°C in a carbogen bubbler system. Tissues in enteric neuron digestion buffer incubated for 1 hour, while tissues in smooth muscle digestion buffer incubated for 10 minutes. For neuron cultures, cells were centrifuged at 350g for 8 minutes at 4°C. The pellet was resuspended in 5mLs of pre-warmed 0.05% Trypsin-EDTA solution for 7 minutes at 37°C. After quenching and centrifugation at 350g for 8 minutes, cells were resuspended in neuron media (Neurobasal A, 1x B27, 2mM L-Glutamine, 1x penicillin/streptomycin, 1mM sodium pyruvate, 1% FBS, and 10ng/mL GDNF), filtered through a 100 micron nylon mesh filter, and plated onto laminin coated plates. For smooth muscle cells, pellet was triturated prior to centrifugation at 350g for 8 minutes at 4°C before resuspending in smooth muscle media (DMEM, 2mM L-glutamine, 1x penicillin/streptomycin, 1mM sodium pyruvate, and 10% FBS) and plating on laminin coated plates. Both smooth muscle cells and enteric neurons were fed daily and imaged within 1 week.

### Gliotransmitter analysis

Compounds identified from LCMS to be PIEZO2 and force dependent were identified and filtered for biological relevance. Spermidine (Sigma, S2501) was dissolved in sterile water at 10mM and used at 10μM. Aspartic acid (Sigma, A5474) was dissolved in water with NaOH at 10mM and used at 10μM. BzATP (Cayman Chemical, 15577) was dissolved in sterile water at 10mM and used at 100μM. Linoleic acid (Sigma, L5900) was dissolved in sterile water at 10mM and used at 10μM.

Enteric neuron and smooth muscle media was changed from growth media as described above to KREBS buffer, washed gently 3 times, then loaded with KREBS plus 1 μM Fluo4, AM (Thermo Fisher, F14201) with 0.02% Pluronic F-127 to disperse the nonpolar AM ester. Cells were incubated in this media for at least 30 minutes at 37°C protected from light before imaging via an Olympus FV3000 confocal microscope with a Tokai Hit 5% CO2 stage top incubator. Cells were imaged for 120 seconds at baseline with no stimulation, before slowly dispensing candidate gliotransmitters into the culture media using syringe pump system. Images continued for a total of 5 minutes. Images were analyzed using Fiji and ImageJ plugin Time Series Analyzer ver3. Briefly, the cell regions were defined as regions of interest (ROI) and circled around then obtained the pixel intensity with Time Series Analyzer. Background nonspecific muscle staining/ autofluorescence was subtracted from each picture by using the defined region without Fluo4, AM expression.

### Statistical analysis

P values were calculated as indicated in figure legends using two-sided Welch’s t-test, one way Welch’s ANOVA, or two-way Welch’s ANOVA with multiple comparisons as indicated in the figure legends. Data are presented as mean ± SEM. N represents number of biological replicates, n represents number of independent experimental samples. Individual biological replicates are shown as dots on all plots.

## Supporting information

Auxiliary Code Files

## Reporting summary

Further information on research design is available in the Reporting Summary linked to this article.

## Data availability

All processed datasets generated from this work are included as Supplementary Tables, while Fastq files and processed sequencing datasets are available at the Gene Expression Omnibus (GEO) under accession number GSE285136, with snRNAseq of cortical tissue in GSE232703 and MERSCOPE data in GSE308108. The following publicly available datasets were used in this manuscript: GSE103892 for spinal cord, GSE172204 for ventromedial hypothalamus, GSE182098 for vagal, sural, sciatic, and peroneal nerves, GSE175421 for sensory and sympathetic ganglia, GSE132044 for cortex, GSE156905 for distal colon myenteric plexus, GSE229322 for sorted Sox10-cre; INTACT cells from the colon, GSE149524 for sorted Baf53b-cre; RCL-tdTomato cells from the myenteric plexus of the small intestine, SRP135960 for sorted Wnt1-cre; RCL-tdTomato cells from the myenteric plexus, GSE182506 for tunica muscularis from the duodenum, GSE237713 for tunica muscularis from the small intestine throughout development, SCP1038 for enteric nervous system cells from the colon and ileum, GSE227742 for small intestinal spatial transcriptomics, and GSE153202 for enteric neurons from the colon and ileum.

## Code availability

Data analysis was performed with publicly available packages by following available tutorials and vignettes. Annotated code is available upon request. See Auxiliary Files 1-4.

## Acknowledgments

This research was supported by grants from the Howard Hughes Medical Institute Hanna H. Gray Fellowship (MAS), The New York Stem Cell Foundation Druckenmiller Fellowship (MAS), National Institutes of Health R35NS116842 (PJT), National Institutes of Health R01CA160356 (PJT), National Institutes of Health F31NS124282 (EFC), National Institutes of Health T32NS077888 (EFC, JJZ), National Institutes of Health F30HD096784 (KCA), National Institutes of Health TL1TR002549 (WJW), National Institutes of Health T32GM152319 (WJW), National Institutes of Health T32GM007250 (KCL, YX, KCA, EFC, WJW, JJZ), Simons Foundation SFARI978897 (PJT), Simons Foundation AN-SURFiN-00010569 (NSA), National Multiple Sclerosis Society TA-2105-37619 (BLLC), and the National Institute of General Medical Sciences, R25-GM075207 (CPT). Additional support was provided by the Small Molecule Drug Development, Genomics, and Light Microscopy and Imaging core facilities of the Case Western Reserve University Comprehensive Cancer Center (P30CA043703), access to multiphoton imaging in the lab of Gabriel Victora, I. Ampong in the Proteomics and Metabolomics Shared Laboratory Resources Center at Cleveland Clinic Foundation, and M. Drumm and The Research Institute for Children’s Health. Philanthropic support was generously contributed by sTF5 Care and the Long, Goodman, and Geller families. We thank K. Lovrenert, N. Datko, A. Vadivel, M. Madhavan, N. Abercrombie, A. Kerr, A. Sturno, A. Lawless, I. Barnett, F. Luna-Polanco, C. Maysonet, A. Scavuzzo, K. Cruz, A. Kumar, K. Keeran, A. Letai, and all other Tesar lab members for helpful discussions and technical assistance. We thank U. Gurkan for intestine-on-a-chip resources and C. Lilliehook for editorial support.

## Author contributions

Conceptualization, MAS and PJT; design and methodology: MAS and PJT; snRNA-seq experiments and analyses: MAS; Tissue processing, immunostaining, cell culture, microscopy, and quantification: MAS, KCL, IKS, JSR, AT, AKS, KCA, BA; MERFISH and Visium spatial transcriptomics: MAS, CPT, AP; Mouse management: MAS, KCL, IKS, AT, BWA, NSA, AKS, and JSR; Biochip generation: WJW; Intestine-on-a-chip experiments and quantifications: MAS, WJW, JJZ, CPT, and YMH; RNAscope: AMH, BLLC; Bioreactor assays: MAS, KCL, EFC; Phenotyping: MAS, KCL, JSR, AT, IKS, HES, EFC, BWA, NSA, CPT, AMH, YX, and YMH; Funding acquisition: MAS and PJT; Project administration: MAS and PJT; Supervision: MAS and PJT; Writing – original draft: MAS; Writing – review & editing: MAS and PJT with input and approval from all authors.

## Competing interests

M.A.S. and P.J.T. are listed as inventors on pending patent claims filed by Case Western Reserve University covering methods for high degradation snRNA-seq (*CitraPrep*). M.A.S., K.C.L., and P.J.T. are listed as inventors on pending patent claims filed by Case Western Reserve University covering methods for scalable enteric glial cell culture. The other authors declare no competing interests related to this work.

**Supplementary Table 1. A proximal-to-distal atlas of enteric glial cells.**

---

Tab 1) Differential expression of annotated cell types from whole duodenal tissue.

Tab 2) Differential expression of unsupervised clusters from whole duodenal tissue.

Tab 3) Differential expression of non-epithelial annotated unsupervised subclusters from duodenum.

Tab 4) Differential expression of unsupervised clusters from merged enteric glial cell datasets.

**Supplementary Table 2. Differentially expressed genes across glia in the body.**

Tab 1) Differential expression of unsupervised clusters from glia throughout the body.

Tab 2) Differential expression of glial cells comparing across tissue of origin.

Tab 3) Differential expression of glial cells comparing by cell type.

**Supplementary Table 3. Regionally enriched transcriptional profiles of glia in the duodenum and colon.**

Tab 1) Differential expression of unsupervised clusters from enteric glia in the duodenum, ileum, and colon.

Tab 2) Differential expression of enteric glial cells compared by tissue of origin.

Tab 3) MERFISH analysis of 140 transcripts across duodenum and colon.

Tab 4) MERFISH differential expression analysis from all cells. Tab 5) MERFISH probe design.

**Supplementary Table 4. Molecular diversity of the enteric nervous system.**

Tab 1) Differential expression of unsupervised clusters from the duodenal enteric nervous system.

Tab 2) Comparison of marker genes from spatial transcriptomics unsupervised clusters with single nucleus RNA-sequencing unsupervised clusters.

**Supplementary Table 5. Functional interaction network of cells in the adult duodenum.**

Tab 1) Ligand-receptor pairs to and from all cells across the duodenum.

Tab 2) Ligand-receptor pairs from enteric glial subtypes to all other cells.

Tab 3) Ligand-receptor pairs from EGC6 to all other cells.

**Supplementary Table 6. Bulk RNA-seq of LMMP tissue from Piezo2 Sox10-creERt2 depleted and littermate controls.**

Tab 1) DESeq2 for differential expression in the whole tunica muscularis from Piezo2 Sox10-creERt2 depletion and littermate controls.

Tab 2) FPKM from each individual biological replicate.

**Supplementary Table 7. Untargeted metabolomics. Related to Figure 5**.

Tab 1) All metabolite features including unknowns mapped by LC-MS.

Tab 2) Metabolite features and levels across all samples including a KREBS blank.

Tab 3) Metabolite features from control timecourse with spiked ACh and GABA standards. Tab 4) Quality control statistics.

Tab 5) Fold change results of before and after stretch Piezo2 Sox10-creERt2 depletion compared to control.

Tab 6) Volcano analysis results of Piezo2 Sox10-creERt2 depletion compared to control, at raw p value or FDR < 0.05.

Tab 7) Orthogonal least square discriminant analysis variables important in the projection, Piezo2 Sox10-creERt2 depletion compared to control.

Tab 8) Orthogonal least square discriminant analysis variables important in the projection, control force vs baseline.

Tab 9) Orthogonal least square discriminant analysis variables important in the projection, Piezo2 Sox10-creERt2 depletion force vs baseline.

Tab 10) Parameters for data pre-processing for metabolomics data.

**Supplementary Table 8. Molecular diversity of enteric neurons in the tunica muscularis.**

Tab 1) Differential expression of enteric neuron clusters.

**Supplementary Video 1.**

Calcium imaging in GCaMP6f; *Sox10*-cre^ERt2^ mouse intestine-on-a-chip. Flash of light indicates the onset of flow as media washes over tissue. Two cells respond to mechanosensation after onset of flow.

**Supplementary Video 2.**

Calcium imaging in enteric glial cells plated into chip devices with Fluo4AM dye. Video is 300 seconds condensed into 20 frames per second, covering baseline (120 seconds), stimulation (10 seconds of 1 dyn/cm^2^ shear stress), and recovery (170 seconds).

**Supplementary Video 3.**

Calcium imaging in enteric glial cells plated into chip devices with Fluo4AM dye. Video shows cells immediately upon 1 dyn/cm^2^ shear stress to illustrate calcium spreading from PIEZO2+/SOX10+/GFAP+ early responders to late responders.

**Extended Data Fig. 1.**
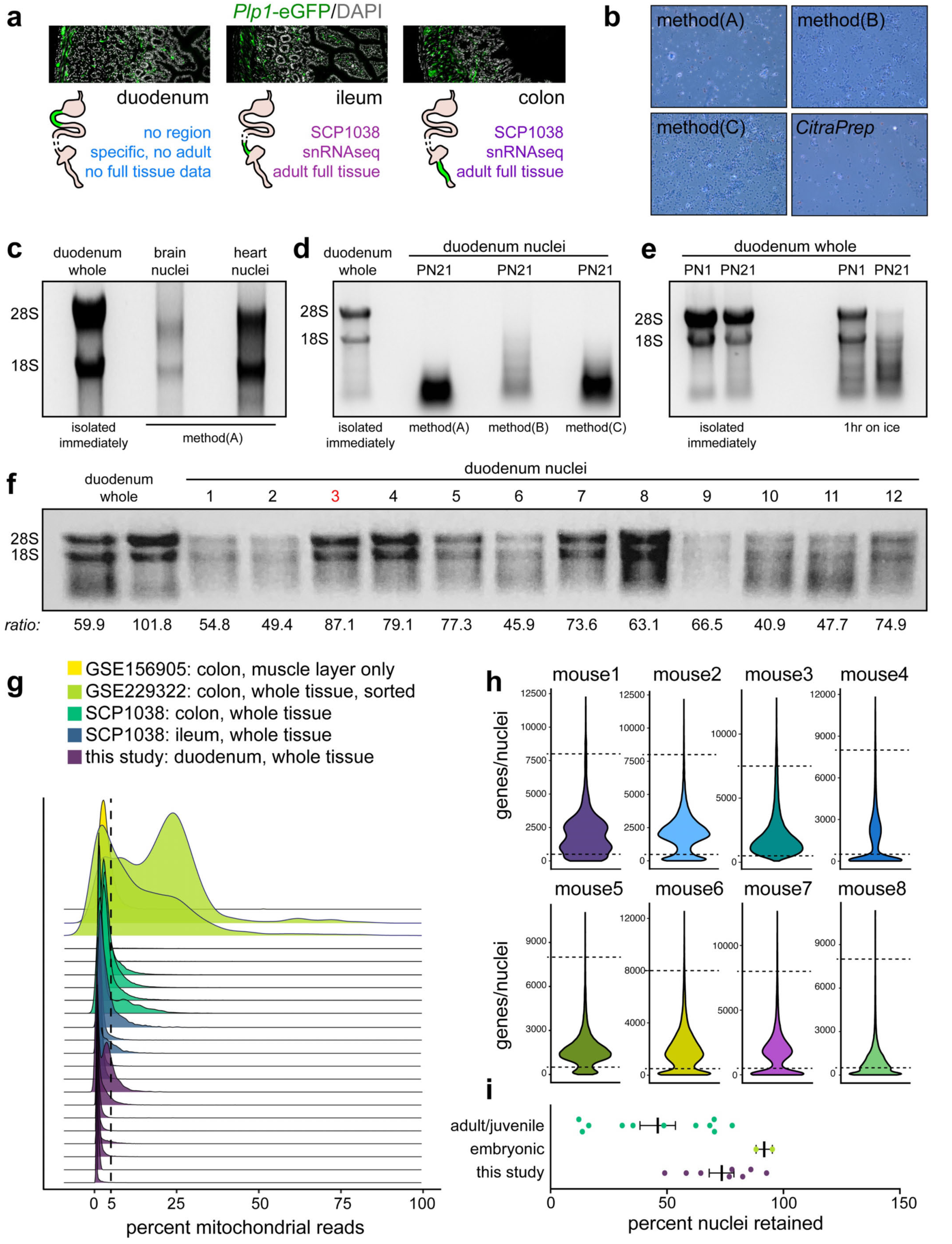
Optimization of *CitraPrep* for single nuclei RNA-seq of high degradation tissue. **a**, Immunostaining of enteric glial cells marked by *Plp1-*eGFP in a cross section of the duodenum, ileum, and colon, showing cells present in every layer of tissue. **b**, Brightfield images of isolated nuclei stained with 0.4% Trypan blue using three established methods (A)^20^, (B)^1^, and (C)^21^, and *CitraPrep.* **c**, RNA formaldehyde gel showing ribosomal RNA 28S and 18S bands in bulk postnatal day 21 duodenal tissue isolated immediately compared to nuclei preparations from the cortex and the heart using an established method^20^. **d**, RNA formaldehyde gel showing ribosomal RNA 28S and 18S bands in bulk postnatal day 21 (PN21) duodenal tissue isolated immediately compared to nuclei preparations from the duodenum using 3 different established methods (A)^20^, (B)^1^, and (C)^21^. **e**, RNA formaldehyde gel showing ribosomal RNA 28S and 18S bands in bulk PN1 and PN21 duodenal tissue isolated immediately compared to snap frozen tissue that incubated undisturbed for 1 hour (1hr) on ice prior to isolation. **f**, RNA formaldehyde gel showing ribosomal RNA 28S and 18S bands in bulk PN1 and PN21 duodenal tissue isolated immediately compared to iterations of nuclei preparation chemistry. Ratio of 28S to 18S band is shown below to score each method. The 10mM citric acid, 7 minute incubation time resulted in the best results by 28S:18S ratio at 87.1. **g**, Distribution of mitochondrial reads sequenced in each individual animal compared to publicly available ENS datasets from other gut regions. Cutoffs for filtering for further analysis were set at 5% and shown by a dashed line. **h**, Violin plots showing genes/nuclei in each individual sample. Dashed lines indicate thresholds for filtering. **i**, Comparison of our datasets with publicly available embryonic intestine and publicly available adult/juvenile gastrointestinal tissue datasets. Bold line is mean with SEM. Each dot is an individual sample with percent nuclei retained after filtering for high quality nuclei (mitochondria <5%, genes between 500 and 8000 per nuclei).

**Extended Data Fig. 2.**
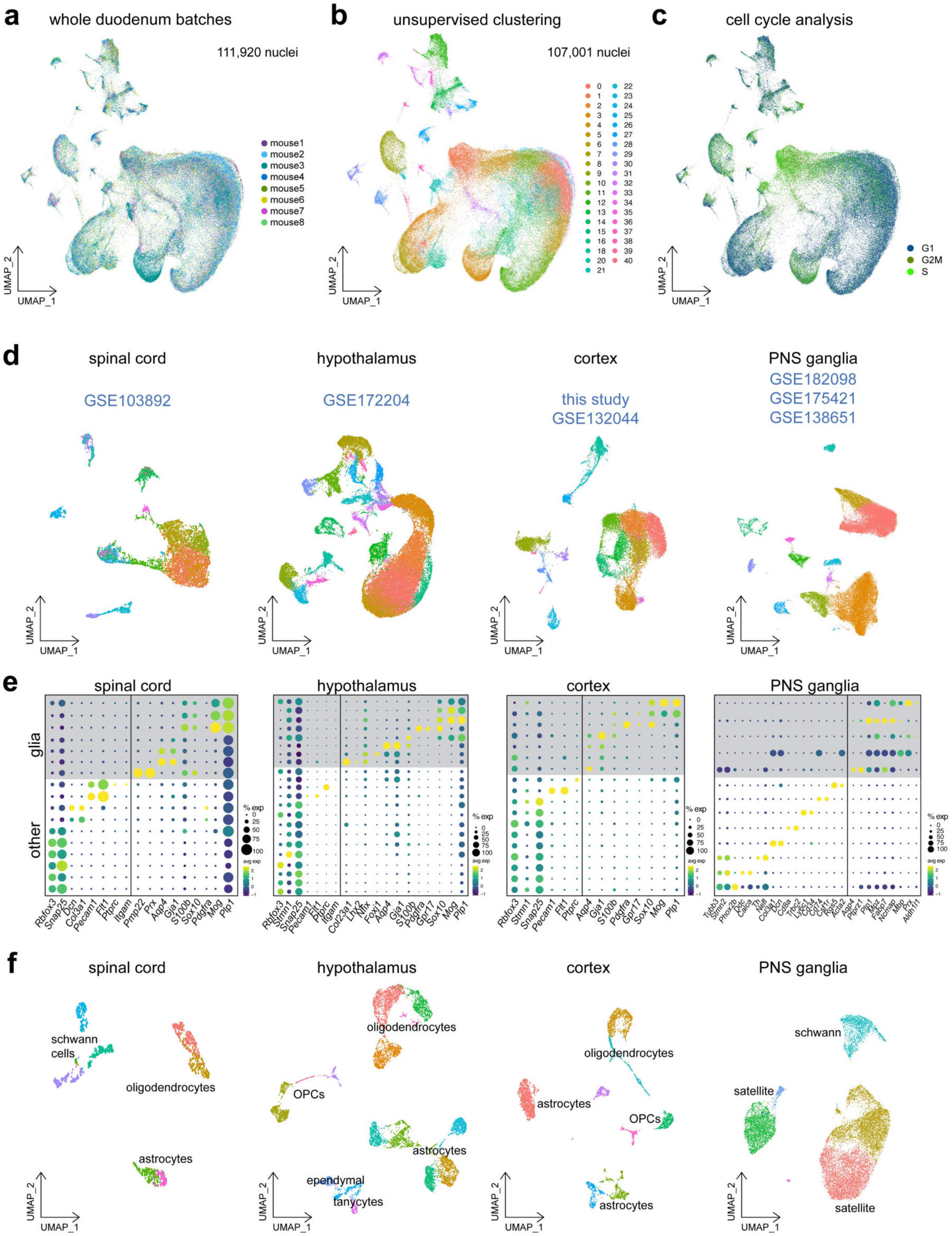
Blueprints of the duodenum, spinal cord, hypothalamus, cortex, and peripheral ganglia. **a**, snRNAseq of duodenal nuclei (N=8 mice, n=111,920 nuclei) visualized by UMAP and colored by sample identity. **b**, UMAP of duodenal nuclei after removal of two low quality clusters leaving 107,001 high quality nuclear transcriptomes colored by unsupervised clusters. **c**, Cell cycle scoring of duodenal nuclei based on expression of G1, G2/M, and S phase markers projected onto UMAP. All data are in Supplementary Table 1. **d**, snRNAseq of tissues from studies visualized by UMAP and colored by unsupervised clustering. **e**, Cell type markers shown by dot plot across unsupervised clusters. Transcripts are shown in each column and annotated below with unsupervised clusters shown in rows. Clusters with expression of glial markers and low expression of other markers (neurons, endothelial cells, microglia) are shown with gray boxes and selected for subclustering. The size of each circle indicates percentage of cells in the cluster that express the marker (≥1 UMI) while the color shows the average expression of transcript in cells. **f**, Subclustered glial cell types from (c), with unsupervised clusters labeled with cell type identity. These nuclei were used in Fig. 1f-h.

**Extended Data Fig. 3.**
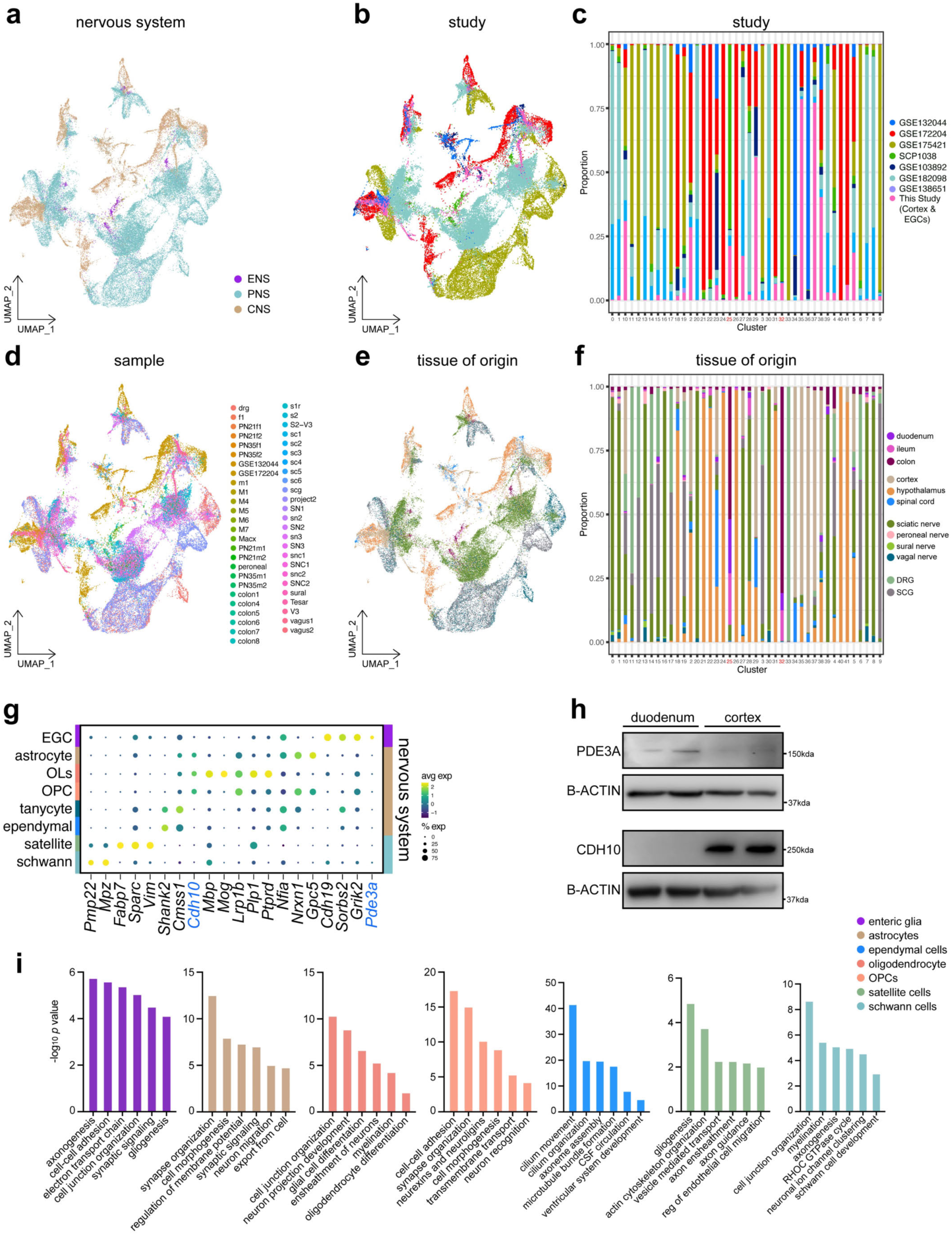
Comparison of glial cells throughout the body. **a** to **f**, Glial cells from Fig. 1f visualized by UMAP and colored by nervous system division (**b**), study (**b-c**), sample (**d**), and tissue of origin (**e-f**). Proportion mapping is shown in for study in (**c**) and for tissue of origin in (**f**). **g**, Dot plot showing expression of select differentially expressed genes across cell types. Cell types are shown in rows with nervous system depicted by colors on the right matching (**a**), with cell type annotated and colored at the left. Column show transcripts with validated differential expression highlighted in blue. The size of each circle indicates percentage of cells in the cluster that express the marker (≥1 UMI) while the color shows the average expression of transcript in cells. All data are in Supplementary Table 2. **h**, Western blot of select differentially expressed genes PDE3A and CDH10 in whole duodenum and whole cortex protein lysate showing differential protein expression. Each lane is a biological replicate. **i**, Gene ontology analysis of top 100 enriched transcripts of each glial cell type compared to other glia in the body using Metascape.

**Extended Data Fig. 4.**
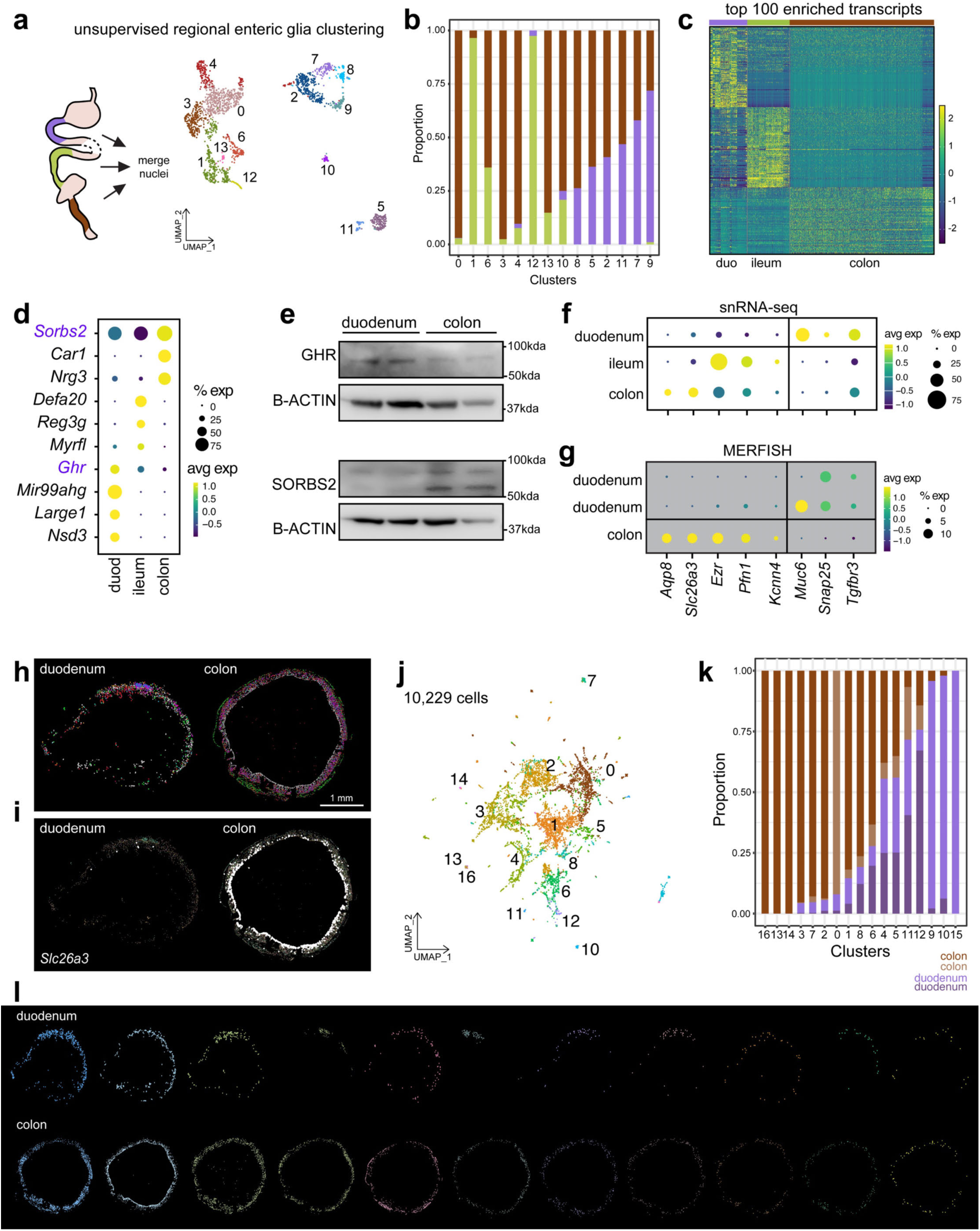
Regional analysis of enteric glial cells. **a**, On left, schematic of study design. On right, snRNAseq of glial cells spanning the foregut, midgut, and hindgut colored by unsupervised clusters and visualized by UMAP (N=14 mice, n=2,257 nuclei). **b**, Proportion mapping of tissue origin to unsupervised clusters from snRNA-seq. Tissue types are shown by color with key shown to the right. Proportion of cells are shown on the y-axis while columns binned in the x-axis show unsupervised clusters. **c**, Top 100 differentially expressed transcripts enriched between tissue regions. Individual transcripts are shown in rows, tissue origins are clustered together in columns with color annotation above. All data are in Supplementary Table 3. **d**, Dot plot showing expression of select differentially expressed genes across tissues. The size of each circle indicates percentage of cells in the cluster that express the marker (≥1 UMI) while the color shows the average expression of transcript in cells. **e**, Western blot of select differentially expressed genes GHR and SORBS2 in whole duodenum and whole colon protein lysate showing differential protein expression. Each lane is a biological replicate. **f-g**, Dot plot showing tissue enriched (**f**) snRNA-seq and MERFISH (**g**) marker expression (columns) in tissues (rows). The size of each circle indicates percentage of cells in the cluster that express the marker (≥1 UMI) while the color shows the average expression of transcript in cells. **h**, Single molecule spatial transcriptomics of 140 differentially expressed RNAs measured with MERFISH in the mouse duodenum and colon at 6 weeks of age. Both tissues are derived from the same animal. Dots represent unsupervised clusters. Scale bar is 1mm. **i**, Spatial distribution of *Slc26a3* RNA measured with MERFISH in the mouse duodenum and colon. **j**, Combined UMAP of two biological replicates and four tissues profiling 10,229 cells. **k**, Proportion mapping of tissue origin to unsupervised clusters from MERFISH from both biological replicates. Tissue types are shown by color from two animals with key shown to the bottom. Proportion of cells are shown on the y-axis while columns binned in the x-axis show unsupervised clusters. **l**, Spatial distribution of unsupervised clusters identified via MERFISH in the duodenum and colon. **g**, UMAP showing tissue identity of nuclei from MERFISH.

**Extended Data Fig. 5.**
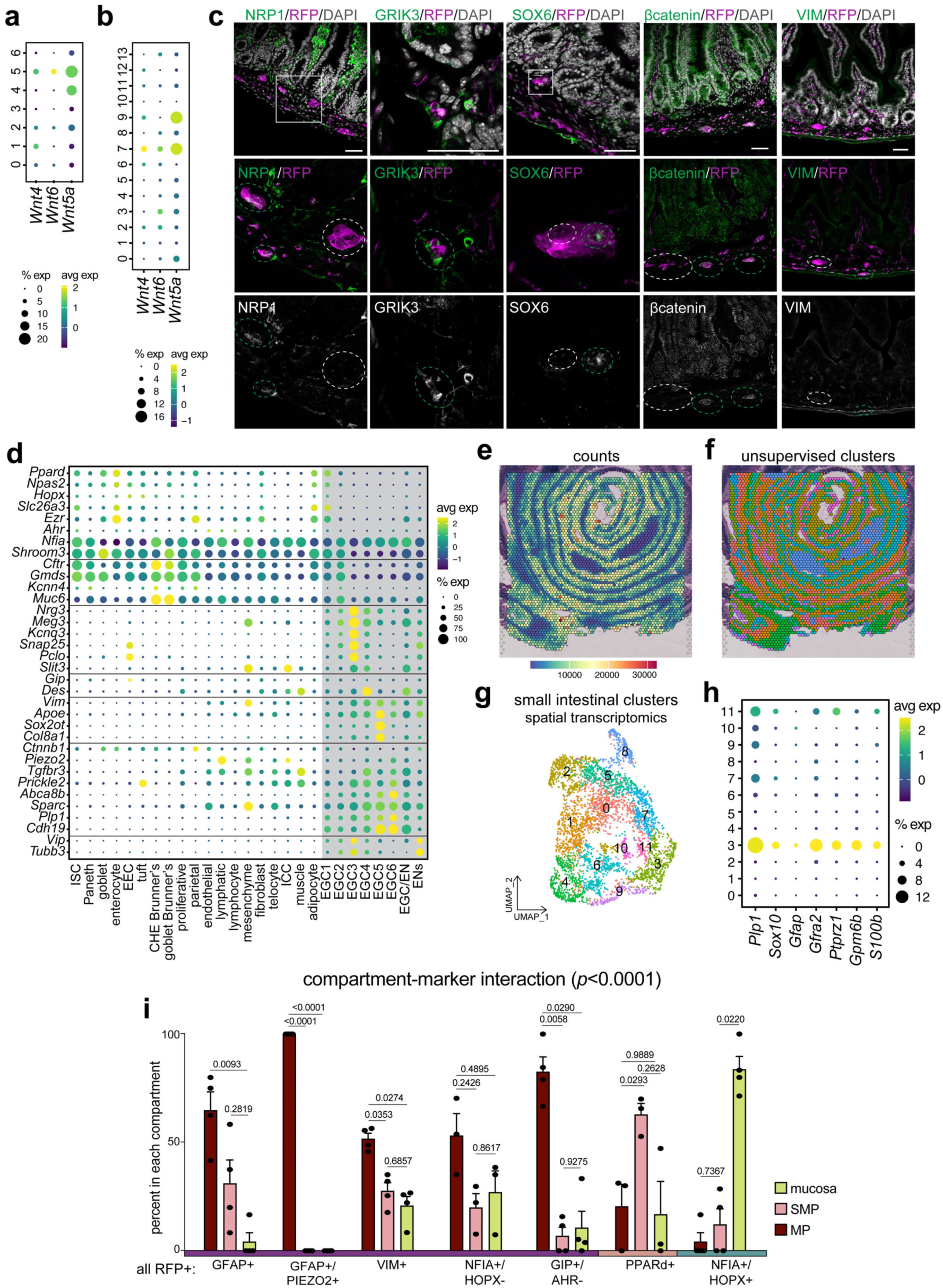
Enteric glia are diverse. **a-b**, Dot plot showing expression of select differentially expressed genes across tissues. Expression of select Expression of *Wnt* ligands reported in a previous study^45^ in enteric nervous system clusters throughout the gut in (**a**) and in duodenum in (**b**). Unsupervised clusters from each tissue are shown in columns, rows show transcripts. The size of each circle indicates percentage of cells in the cluster that express the marker (≥1 UMI) while the color shows the average expression of transcript in cells. **c**, Immunostaining showing enteric glia subtype markers expressed only in subsets of cells imaged on confocal showing z stacks. RFP marks tdTomato from *Sox10-*cre; RCL-tdT mice. DAPI marks nuclei in white. White boxes show area that is enlarged below in separate channels. Green dashed ovals show enteric nervous system cells co-expressing the marker of interest, while white dashed ovals show enteric nervous system cells not expressing the marker of interest. Scale bar = 50um. **d**, Dot plot showing expression of select enteric glial cell genes across duodenal cell types. The size of each circle indicates percentage of cells in the cluster that express the marker (≥1 UMI) while the color shows the average expression of transcript in cells. **e**, Spatial transcriptomics of the small intestine. Each dot is a barcoded region and sequencing depth is shown via counts. **f**, Unsupervised clusters as shown in Extended Data Fig. 5e projected onto tissue. **g**, UMAP of unsupervised clusters from all small intestinal dots shown in Extended Data Fig. 5e-f. **h**, Dot plot showing expression of select enteric glial cell genes across unsupervised clusters. The size of each circle indicates percentage of cells in the cluster that express the marker (≥1 UMI) while the color shows the average expression of transcript in cells. **i**, Quantification of the spatial distribution of enteric glial cell subtype enriched markers in different intestinal compartments. P values from two-way ANOVA with multiple comparisons. Plots show mean with SEM; biological replicates shown as individual dots.

**Extended Data Fig. 6.**
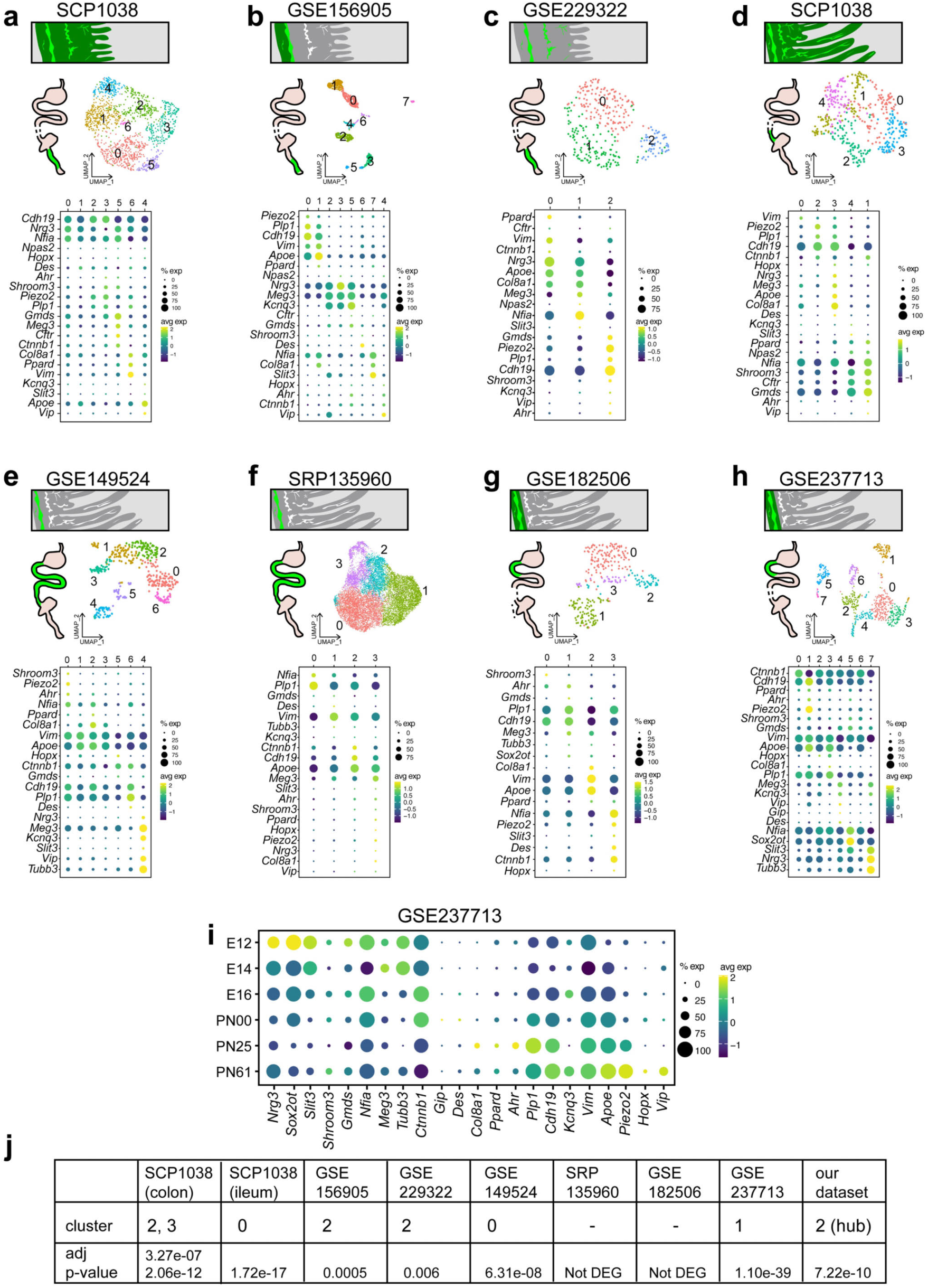
EGC6 cell signature is present across all myenteric datasets, while submucosal EGC1 signature is absent. **a-h**, Schematic of gastrointestinal tissue profiled with cells and regions captured highlighted in green below the study accession number. UMAP plot shows unsupervised clustering of subsetted glial cells. Dot plot showing expression of differentially expressed genes from Fig. 2 across studies. Unsupervised clusters from each tissue are shown in columns, rows show transcripts. The size of each circle indicates percentage of cells in the cluster that express the marker (≥1 UMI) while the color shows the average expression of transcript in cells. **i**, Dot plot showing expression of differentially expressed genes from Fig. 2 throughout development. Timepoints shown in rows with transcripts in columns. E=embryonic, PN=postnatal. The size of each circle indicates percentage of cells in the cluster that express the marker (≥1 UMI) while the color shows the average expression of transcript in cells. **j**, Table showing *Piezo2* enrichment across publicly available datasets, including which clusters *Piezo2* is differentially expressed in and at what adjusted p-value from the Wilcoxon Rank Sum test.

**Extended Data Fig. 7.**
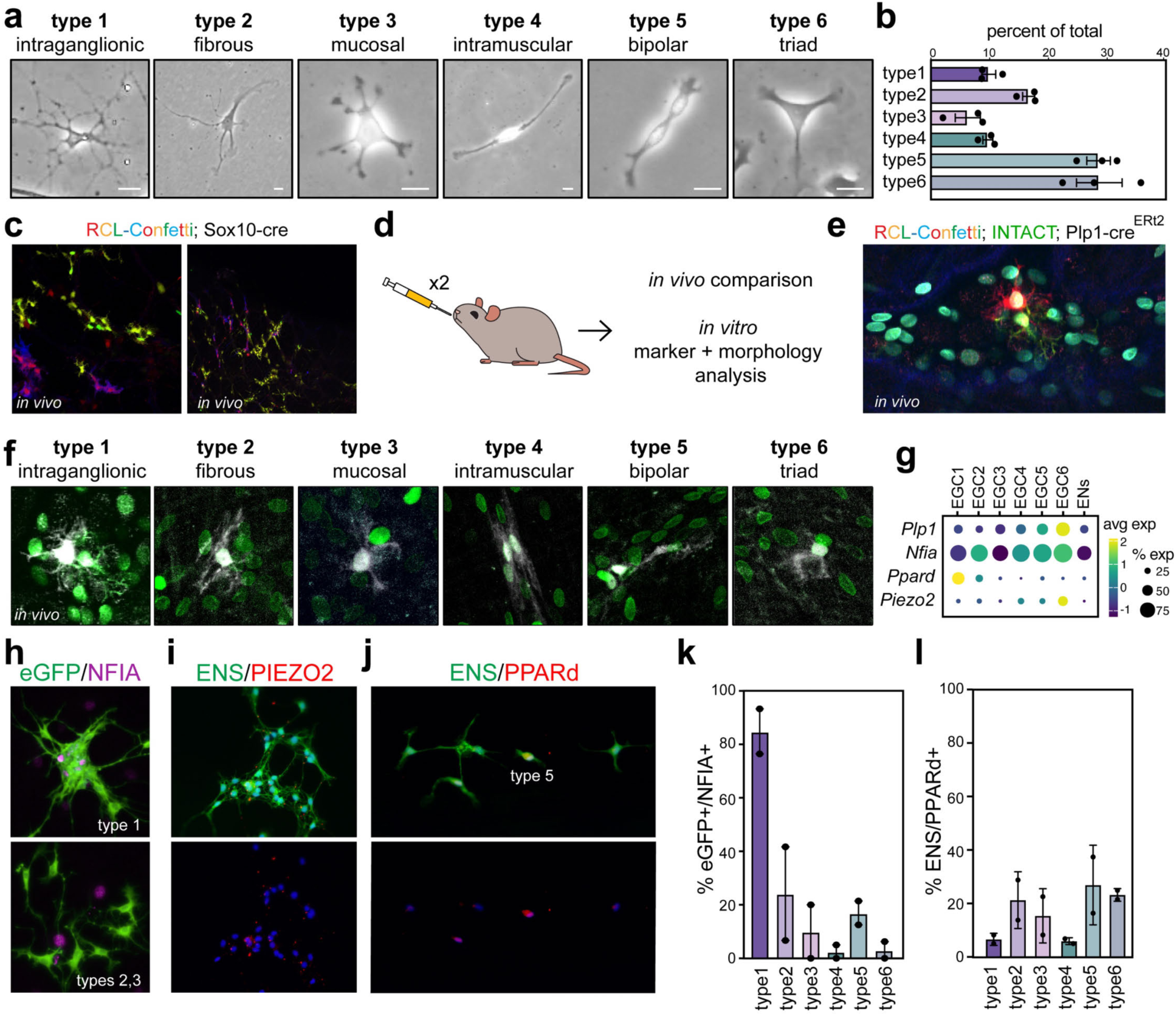
Enteric glia are morphologically diverse. **a**, Representative images of enteric glial cells in culture capturing 6 different morphologies. Scale bars = 20 um. **b**, Quantification of enteric glia morphologies in culture (N=3 mice) as shown in Fig. 3e. **c**, Representative images of RCL-Confetti; Sox10-cre mouse duodenal tissue in 3D after optical clearing. **d**, Schematic of tamoxifen dosing for inducble Cre mouse lines for *in vivo* and *in vitro* morphology comparison. **e**, Representative image of full colorimetric two-photon image in the RCL-Confetti; Plp1-cre^ERt2^; INTACT mouse myenteric plexus. **f**, Sparse *in vivo* multicolor labeling of enteric glial cells by two-photon imaging. Sparsely labeled enteric glia are shown in white with Plp1-cre^ERt2^ labeled INTACT nuclei shown in green. **g**, Dot plot showing transcript levels of genes used for subtype morphology assessment. Rows show transcripts while columns show enteric glial subtypes. The size of each circle indicates percentage of cells in the cluster that express the marker (≥1 UMI) while the color shows the average expression of transcript in cells. **h**, Representative immunostaining of cultured enteric glial cells with different morphologies and marker expression. *Plp1-*eGFP is shown in green while NFIA is shown in magenta. **i**, Representative immunostaining of cultured enteric glial cells. RCL-tdTomato; Sox10-cre is shown in green marking the ENS while PIEZO2 is shown in red. DAPI marks nuclei in blue. Separate channel is shown below. **j**, Representative immunostaining of cultured enteric glial cells. RCL-tdTomato; Sox10-cre is shown in green marking the ENS while PPARd is shown in red. DAPI marks nuclei in blue. Separate channel is shown below. **k**, Quantification of NFIA/eGFP+ marking EGC6 cells in each morphological group shown as percentage. Each dot represents a biological replicate. P values show one-way ANOVA with unpaired t and Welch’s correction. **l**, Quantification of PPARd/ RCL-tdTomato; Sox10-cre+ marking EGC1 in each morphological group shown as percentage. Each dot represents a biological replicate. P values show one-way ANOVA with unpaired t and Welch’s correction.

**Extended Data Fig. 8.**
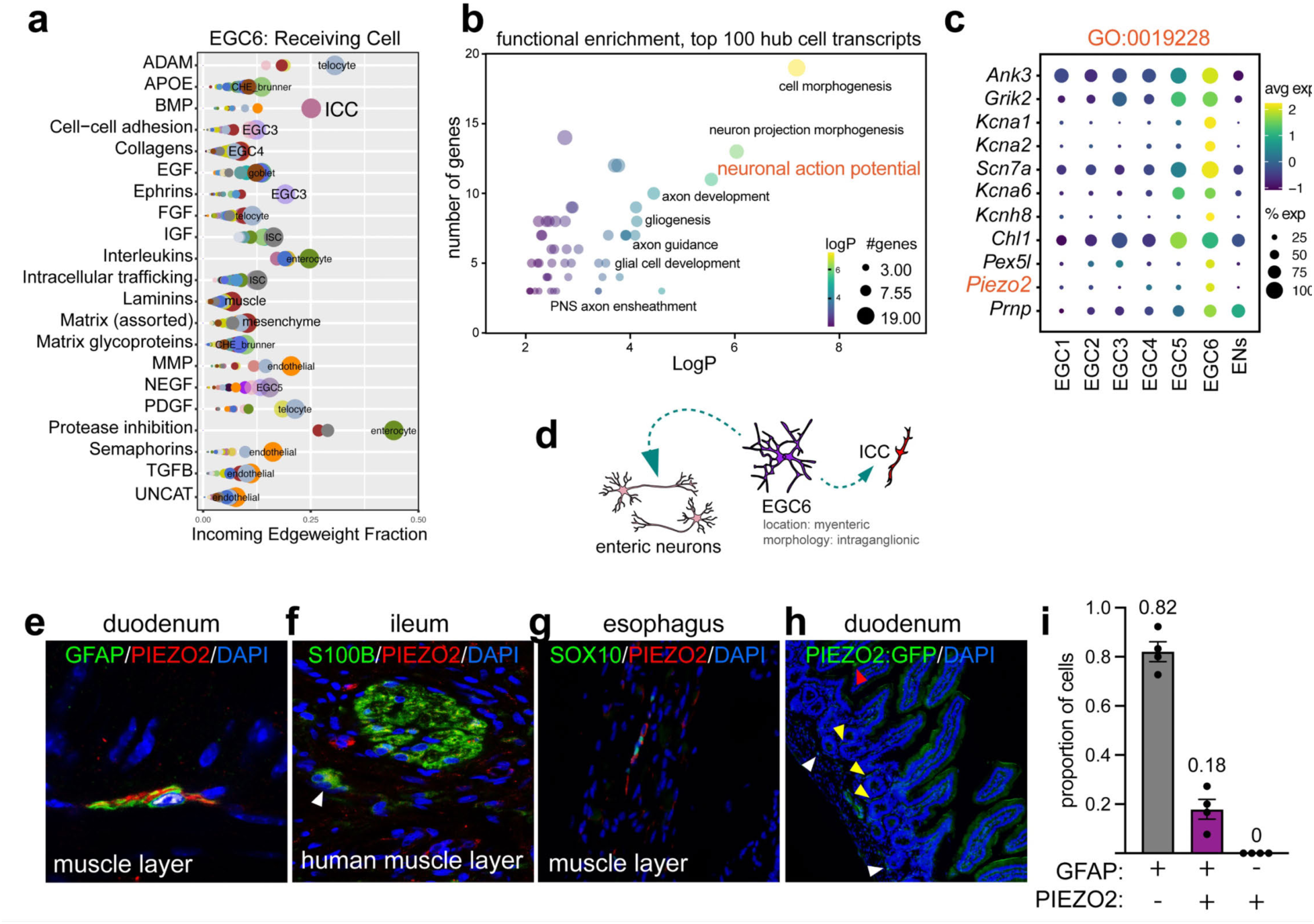
Subtype-specific enteric glia functional interaction network. **a,** Interactome analyses of putative ligand-receptor pairs using the FANTOM5 database (*p*<0.00001, percent>0.2). Signaling classes are grouped in rows with enriched cell types shown across the x-axis. Datasets are provided in Supplementary Table 5. **b**, Bubble plot showing gene ontology analysis of the top 100 enriched transcripts in EGC6 compared to other cells in the ENS. Dot size shows number of genes while color shows P value. **c**, Dot plot showing transcript levels of genes associated with neuronal action potential in duodenal enteric glia subtypes. Rows show transcripts while columns show enteric glial subtypes. The size of each circle indicates percentage of cells in the cluster that express the marker (≥1 UMI) while the color shows the average expression of transcript in cells. **d**, Depiction of enteric nervous system and signaling relationships. EGC6 cells are dominated by intraganglionic morphologies in the myenteric plexus. **e**, Longitudinal muscle myenteric plexus preparation stained for SOX10, PIEZO2, and GFAP with DAPI shown in blue. **f**, Human enteric ganglia in the ileum muscle layer with enteric glia in green and PIEZO2 in red with DAPI showing nuclei in blue. **g**, Immunostaining of PIEZO2 in enteric glia in the mouse esophagus. **h**, Expression of PIEZO2:GFP with DAPI shown in blue in the duodenum. Red arrow shows enterochromaffin cells in the epithelium positive for PIEZO2:GFP where it is known to be expressed. Yellow arrows show basement membrane cells beneath the epithelium with mesenchymal morphology. White arrows show ganglia like morphology with PIEZO2:GFP positivity. **i**, Quantification of PIEZO2+ cells and GFAP+ cells. Each dot shows a biological replicate.

**Extended Data Fig. 9.**
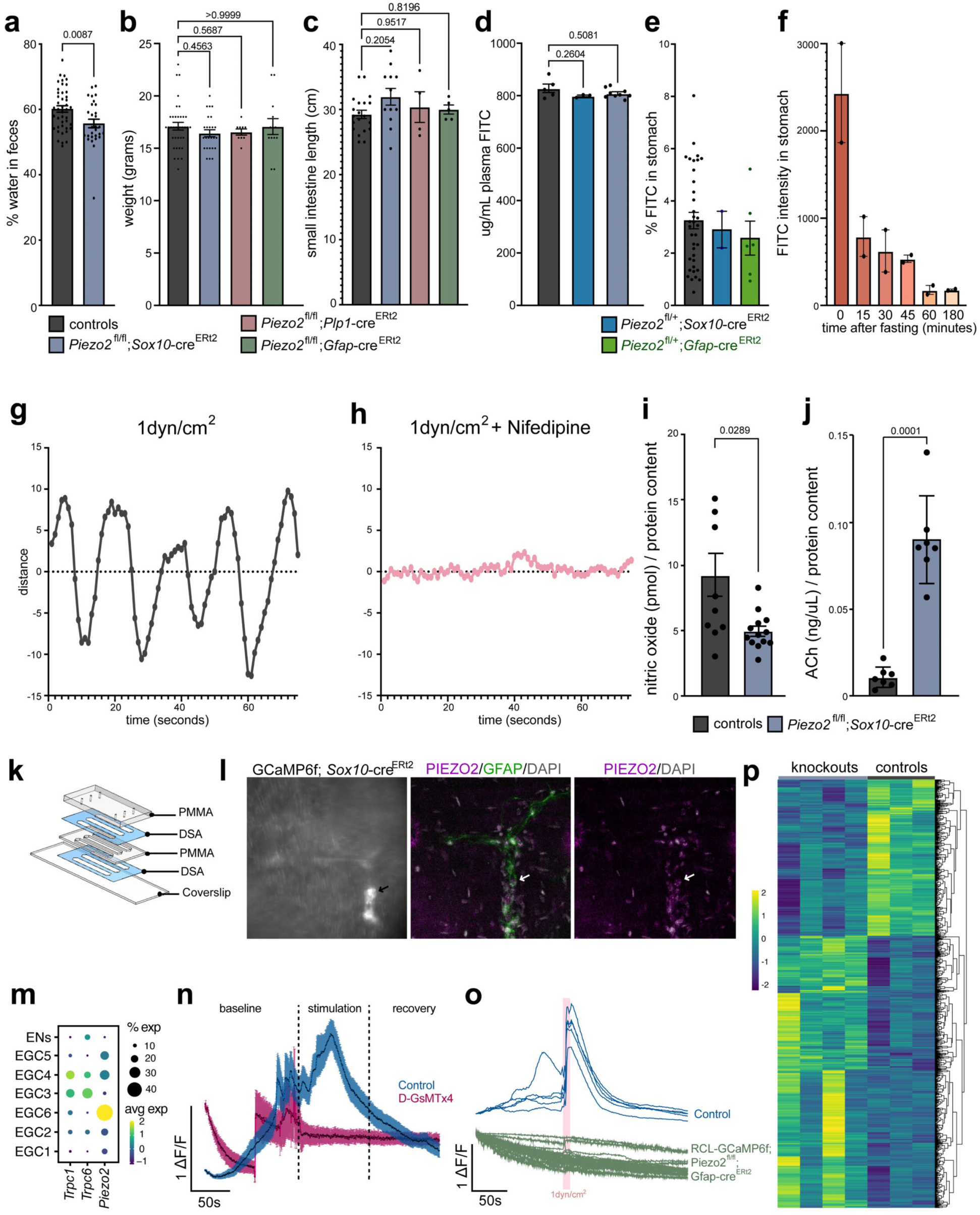
Intestinal physiology after deletion of PIEZO2 from myenteric glia. **a**, Percent fecal water composition. P values from Welch’s unpaired t-test. Plots show mean with SEM; biological replicates shown as individual dots. **b-d,** At six weeks of age, the weight **(b)**, small intestinal length (**c**), and epithelial permeability **(d)** of mice. P values from one-way ANOVA with multiple comparisons. Plots show mean with SEM; biological replicates shown as individual dots. **e,** Gastric emptying in mice with one functional copy of *Piezo2*. **f**, Pilot study determining fasting length for emptying of chow from stomach. **g,** Rhythmic oscillatory contractions in mice after stimulation with shear stress at 1dyn/cm^2^. **h,** Loss of rhythmic oscillations in mice treated with 3uM Nifedipine to block voltage gated calcium channels. **i-j,** Levels of nitric oxide **(i)** and acetylcholine (**j),** in the longitudinal muscle myenteric plexus relative to total protein content. P values from Welch’s unpaired t-test. Plots show mean with SEM; biological replicates shown as individual dots. **k,** Assembled view of the intestine-on-a-chip. Channels are 4 mm wide, 24 mm long, and 1.1 mm in height. Exploded view of the intestine-on-a-chip depicting the individual layers of polymethylmethacrylate (PMMA), double-sided adhesive (DSA) film, and glass. Top PMMA layer is 3.175 mm, DSA layers are 50 µm, and middle PMMA layer is 1 mm. Two types of glass were used; for calcium imaging, cover slips were used for the glass layer while standard microscope slides were used for contraction imaging. **l,** Left: still image from live imaging GCaMP6f upon stimulation with 1dyn/cm^2^ shear stress in GCaMP6f; *Sox10-*cre^ERt2^ mouse longitudinal muscle myenteric plexus. Middle and right: immunostaining of tissue shown on left for PIEZO2 with GFAP marking a subset of enteric glia and DAPI marking nuclei in white. Arrow points to mechanosensory cell co-expressing PIEZO2 with GFAP. **m,** Dot plot showing transcript levels of D-GsMTx4 targets *Trpc1, Trpc6*, and *Piezo2.* The size of each circle indicates percentage of cells in the cluster that express the marker (≥1 UMI) while the color shows the average expression of transcript in cells. **n,** Average traces from calcium imaging of enteric glial cells in culture at baseline for 120 seconds, after stimulation for 10 seconds with 1 dyn/cm^2^ shear stress and the following 80 seconds, and at recovery for 90 seconds in KREBS buffer. After 30 minutes of rest, the cells were treated with 10μM D-GsMTx4 in KREBS buffer for 10 minutes before repeating the paradigm with 10 seconds of shear stress at 1 dyn/cm^2^ with 10μM D-GsMTx4. Dots show the mean individual data points with SEM. n=18 cells. **o,** Individual traces from calcium imaging of enteric glial cells in culture before, during, and after stimulation with shear stress at 1 dyn/cm^2^ for 10 seconds in control animals (n=5) or animals with the genetically encoded calcium indicator GCaMP6f expressed in cells with Gfap-cre^ERt2^ driven knockout of PIEZO2 (n=18). **p,** Heatmap showing differentially expressed genes from the tunica muscularis of littermate control and Piezo2^fl*/fl*^; *Sox10-*cre^ERt2^ mice via bulk RNA-sequencing. Datasets are provided in Supplementary Table 6. For all plots, each dot represents an individual animal.

**Extended Data Fig. 10.**
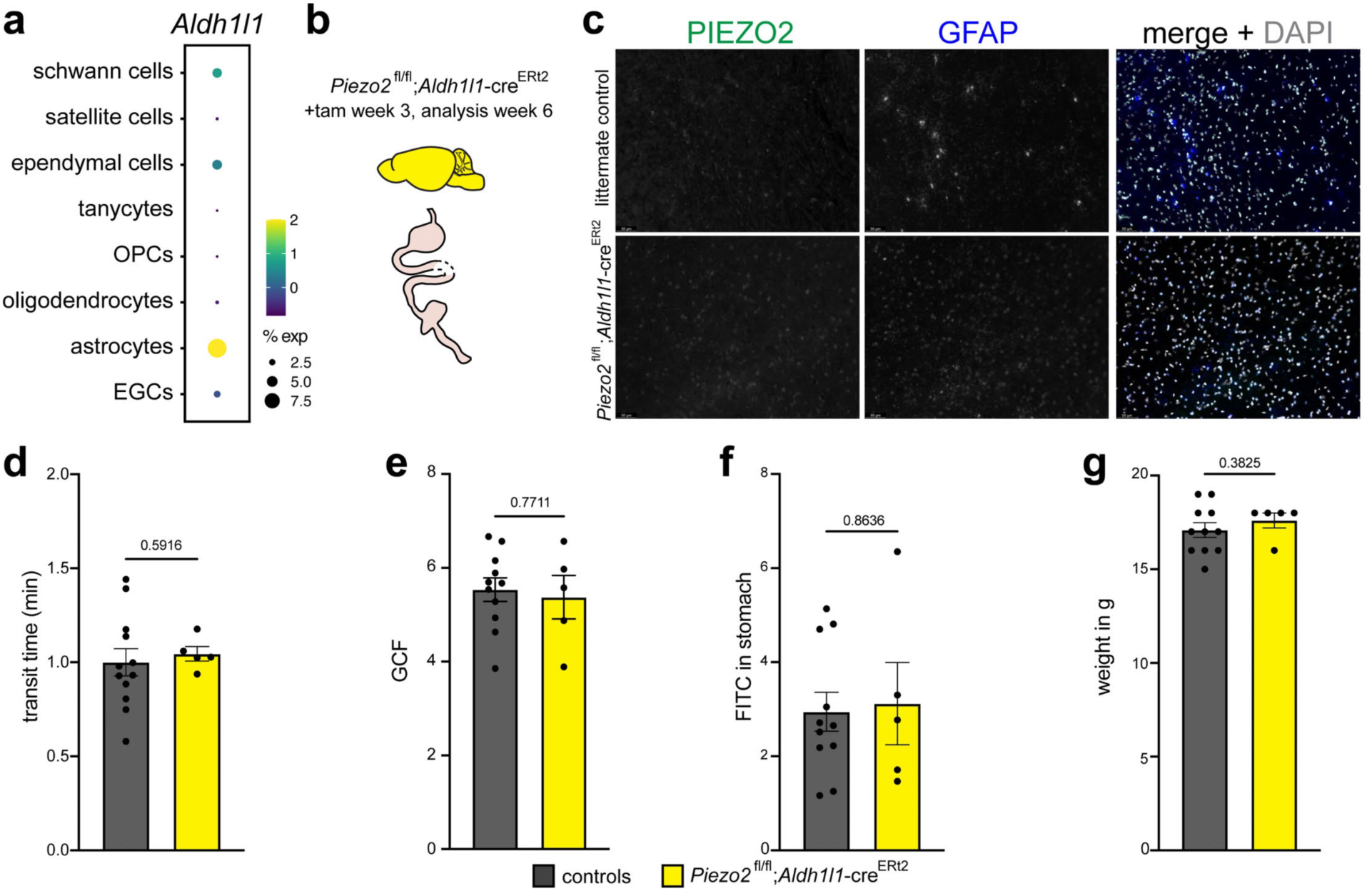
Deleting PIEZO2 from astrocytes does not impact gut motility. **a**, Dot plot showing expression of *Aldh1l1,* in different glial populations throughout the body as shown in Extended Data Fig. 2. The size of each circle indicates percentage of cells in the cluster that express the marker (≥1 UMI) while the color shows the average expression of transcript in cells. **b**, Schematic showing targeting of *Aldh1l1*+ cells for *Piezo2* knockout. **c**, RNAscope in the cortex of Piezo2^fl/fl^; *Aldh1l1-*cre^ERt2^ mice and littermate control. Astrocytes are. Marked in blue by *Gfap*. Scale bar = 50 um. **d**, Total gastrointestinal transit time measuring time from gavage of carmine red dye to the appearance of red stool in littermate controls (N=12) and Piezo2^fl/fl^; *Aldh1l1*-cre^ERt2^ mice (N=5). **e**, The geometric center of fluorescence shows the location of the bolus in the small intestine in littermate controls (N=11) and Piezo2^fl/fl^; *Aldh1l1*-cre^ERt2^ mice (N=5). **f**, Gastric emptying in littermate controls (N=11) and Piezo2^fl/fl^; *Aldh1l1*-cre^ERt2^ mice (N=5). **g**, Body weight of littermate controls (N=11) and Piezo2^fl/fl^; *Aldh1l1*-cre^ERt2^ mice (N=5). All error bars show SEM with individual mice shown as dots. P values by Welch’s t-test.

