## Supplementary material for "Specialized enteric glial cells coordinate intestinal motility": Auxiliary Code Files: Auxiliary File 1.R

```
library(Connectome)
###connectome

CreateConnectome 

#takes as input a Seurat 3.0 object and uses the active identity slot 
#to define nodes for network analysis. 
###The output of this function is an edgelist connecting pairs of nodes via 
#specific ligand-receptor mechanisms, with many edge attributes allowing 
#downstream processing and filtration.

connectome <- CreateConnectome(duod2, LR.database = "fantom5", 
                               species="mouse", assay="SCT")


FilterConnectome 
#is a streamlined way of reducing the above edgelist (which is generally large) 
#to a smaller subset of edges more likely to be of biological and statistical 
#interest. Identical functionality can be achieved with the base subset() function

sigconnectome <- FilterConnectome(connectome = connectome, 
                                  min.pct=0.5, max.p = 0.000001)

EGC1 <- FilterConnectome(connectome = connectome, 
                                  min.pct=0.2, max.p = 0.00001, sources.include = c("EGC1"))


NetworkPlot 
#provides a simple way to visualize networks of interest. It is a wrapper for 
#the R package igraph, and allows filtration on all arguments which can be 
#passed to 
FilterConnectome.

NetworkPlot(EGC1)

Centrality 
#creates a paired centrality plot allowing identification of cell types 
#dominating production or reception of specific modes of signaling. 
#Allows filtration on all arguments which can be passed to 
FilterConnectome.

Centrality(EGC1)

SignalScatter 
#aids in identification of top cell-cell signaling vectors within a network 
#of interest.

SignalScatter(sigconnectome, "Abca1")
SignalScatter(EGC1, genes.viz)

CellCellScatter 
#aids in identification of top signaling mechanisms between a specified 
#source and target cell of interest.

CellCellScatter(sigconnectome, sources.include = "EGC1", targets.include = "muscle")

CircosPlot 
#replaces the hive plots displayed in the original manuscript. 
#CircosPlot is a versatile plotting function which includes many adaptable 
#parameters and yields easy-to interpret quantitative graphs using the 
#R package circlize. Allows filtration on all arguments which can be passed to 
FilterConnectome. #Both edgeweights can be displayed, and the edges 
#can be colored by either source or target cell. Useful for visualizing 
#niche-networks and mechanism interactomes.


CircosPlot(EGC1, cols.use=grid.col)

EdgeDotPlot 
#is an elegant way to visualize both edgeweights (signal strength and signal 
#specificity) at the same time, for a sub-system of interest. It also shows 
#how highly weighted edges in one population compare to 
#lowly-weighted ones in another, which can be useful for answering key 
#scientific questions.

EdgeDotPlot(EGC1)+scale_color_viridis_c()


####
#CircosPlot(sigconnectome, cols.use=grid.col, min.z=1, big.gap = 1, small.gap = 0.1)
#Pre-filter edges:  5593
#Post-filter edges:  1466


#CircosPlot(sigconnectome, cols.use=grid.col, min.z=2)
#Pre-filter edges:  5593
#Post-filter edges:  665
```
