## Supplementary material for "Specialized enteric glial cells coordinate intestinal motility": Auxiliary Code Files: Auxiliary File 2.R

```
###allglia


#load packages

library(biomaRt)
library(dplyr)
library(celda)
library(Seurat)
library(plotly)
library(e1071)
library(sctransform)
library(viridis)
library(harmony)
library(ggplot2)


###load objects

##########PNS

PNS <- merge(sciatic, pns_scwann, add.cell.ids=NULL, project="merge")

PNS <- SCTransform(PNS, vars.to.regress = "percent.mt", assay = "SCT", verbose=FALSE, return.only.var.genes = FALSE)
PNS <- RunPCA(PNS, features = VariableFeatures(object = PNS))

stdv <- PNS[["pca"]]@stdev
sum.stdv <- sum(PNS[["pca"]]@stdev)
percent.stdv <- (stdv / sum.stdv) * 100

cumulative <- cumsum(percent.stdv)

co1 <- which(cumulative > 90 & percent.stdv < 5)[1]
co1

co2 <- sort(which(
  (percent.stdv[1:length(percent.stdv) - 1] - 
     percent.stdv[2:length(percent.stdv)]) > 0.1), 
  decreasing = T)[1] + 1

co2

min.pc <- min(co1, co2)
min.pc


plot_df <- data.frame(pct = percent.stdv, 
                      cumu = cumulative, 
                      rank = 1:length(percent.stdv))

ggplot(plot_df, aes(cumulative, percent.stdv, label = rank, color = rank > min.pc)) + 
  geom_text() + 
  geom_vline(xintercept = 90, color = "grey") + 
  geom_hline(yintercept = min(percent.stdv[percent.stdv > 5]), color = "grey") +
  theme_bw()


PNS <- RunHarmony(PNS, group.by.vars="orig.ident", assay.use="SCT")
PNS <- RunUMAP(PNS, dims=1:min.pc, reduction="harmony")
PNS <- FindNeighbors(PNS, dims = 1:min.pc, assay="SCT", reduction="harmony")
PNS <- FindClusters(PNS, resolution = 0.5)

DimPlot(PNS, reduction = "umap", label=T)
DimPlot(PNS, reduction = "umap", group.by="origin")
DimPlot(PNS, reduction = "umap", group.by="celltype")

PNS_glia <- merge(satelliteschwann, spinalcordglia, add.cell.ids=NULL, project="merge")

PNS_glia <- SCTransform(PNS_glia, vars.to.regress = "percent.mt", assay = "SCT", verbose=FALSE, return.only.var.genes = FALSE)
PNS_glia <- RunPCA(PNS_glia, features = VariableFeatures(object = PNS_glia))

stdv <- PNS_glia[["pca"]]@stdev
sum.stdv <- sum(PNS_glia[["pca"]]@stdev)
percent.stdv <- (stdv / sum.stdv) * 100

cumulative <- cumsum(percent.stdv)

co1 <- which(cumulative > 90 & percent.stdv < 5)[1]
co1

co2 <- sort(which(
  (percent.stdv[1:length(percent.stdv) - 1] - 
     percent.stdv[2:length(percent.stdv)]) > 0.1), 
  decreasing = T)[1] + 1

co2

min.pc <- min(co1, co2)
min.pc


plot_df <- data.frame(pct = percent.stdv, 
                      cumu = cumulative, 
                      rank = 1:length(percent.stdv))

ggplot(plot_df, aes(cumulative, percent.stdv, label = rank, color = rank > min.pc)) + 
  geom_text() + 
  geom_vline(xintercept = 90, color = "grey") + 
  geom_hline(yintercept = min(percent.stdv[percent.stdv > 5]), color = "grey") +
  theme_bw()


PNS_glia <- RunHarmony(PNS_glia, group.by.vars="orig.ident", assay.use="SCT")
PNS_glia <- RunUMAP(PNS_glia, dims=1:min.pc, reduction="harmony")
PNS_glia <- FindNeighbors(PNS_glia, dims = 1:min.pc, assay="SCT", reduction="harmony")
PNS_glia <- FindClusters(PNS_glia, resolution = 0.5)

DimPlot(PNS_glia, reduction = "umap", label=T)
DimPlot(PNS_glia, reduction = "umap", group.by="origin")
DimPlot(PNS_glia, reduction = "umap", group.by="celltype")


###################here
PNSglia1 <- merge(PNS_glia, vagalglia, add.cell.ids=NULL, project="merge")

PNSglia <- merge(PNSglia1, PNS, add.cell.ids=NULL, project="merge")


PNSglia <- SCTransform(PNSglia, vars.to.regress = "percent.mt", assay = "SCT", verbose=FALSE, return.only.var.genes = FALSE)
PNSglia <- RunPCA(PNSglia, features = VariableFeatures(object = PNSglia))

stdv <- PNSglia[["pca"]]@stdev
sum.stdv <- sum(PNSglia[["pca"]]@stdev)
percent.stdv <- (stdv / sum.stdv) * 100

cumulative <- cumsum(percent.stdv)

co1 <- which(cumulative > 90 & percent.stdv < 5)[1]
co1

co2 <- sort(which(
  (percent.stdv[1:length(percent.stdv) - 1] - 
     percent.stdv[2:length(percent.stdv)]) > 0.1), 
  decreasing = T)[1] + 1

co2

min.pc <- min(co1, co2)
min.pc


plot_df <- data.frame(pct = percent.stdv, 
                      cumu = cumulative, 
                      rank = 1:length(percent.stdv))

ggplot(plot_df, aes(cumulative, percent.stdv, label = rank, color = rank > min.pc)) + 
  geom_text() + 
  geom_vline(xintercept = 90, color = "grey") + 
  geom_hline(yintercept = min(percent.stdv[percent.stdv > 5]), color = "grey") +
  theme_bw()


PNSglia <- RunHarmony(PNSglia, group.by.vars="orig.ident", assay.use="SCT")
PNSglia <- RunUMAP(PNSglia, dims=1:min.pc, reduction="harmony")
PNSglia <- FindNeighbors(PNSglia, dims = 1:min.pc, assay="SCT", reduction="harmony")
PNSglia <- FindClusters(PNSglia, resolution = 0.5)

DimPlot(PNSglia, reduction = "umap", label=T)
DimPlot(PNSglia, reduction = "umap", group.by="origin")
DimPlot(PNSglia, reduction = "umap", group.by="celltype")


saveRDS(PNSglia, file="PNS_12302022.rds")


##########################cortex

cortex <- merge(cortexglia, cortexglia_GSE132044, add.cell.ids=NULL, project="merge")


cortex <- SCTransform(cortex, vars.to.regress = "percent.mt", assay = "SCT", verbose=FALSE, return.only.var.genes = FALSE)
cortex <- RunPCA(cortex, features = VariableFeatures(object = cortex))

stdv <- cortex[["pca"]]@stdev
sum.stdv <- sum(cortex[["pca"]]@stdev)
percent.stdv <- (stdv / sum.stdv) * 100

cumulative <- cumsum(percent.stdv)

co1 <- which(cumulative > 90 & percent.stdv < 5)[1]
co1

co2 <- sort(which(
  (percent.stdv[1:length(percent.stdv) - 1] - 
     percent.stdv[2:length(percent.stdv)]) > 0.1), 
  decreasing = T)[1] + 1

co2

min.pc <- min(co1, co2)
min.pc


plot_df <- data.frame(pct = percent.stdv, 
                      cumu = cumulative, 
                      rank = 1:length(percent.stdv))

ggplot(plot_df, aes(cumulative, percent.stdv, label = rank, color = rank > min.pc)) + 
  geom_text() + 
  geom_vline(xintercept = 90, color = "grey") + 
  geom_hline(yintercept = min(percent.stdv[percent.stdv > 5]), color = "grey") +
  theme_bw()


cortex <- RunHarmony(cortex, group.by.vars="orig.ident", assay.use="SCT")
cortex <- RunUMAP(cortex, dims=1:min.pc, reduction="harmony")
cortex <- FindNeighbors(cortex, dims = 1:min.pc, assay="SCT", reduction="harmony")
cortex <- FindClusters(cortex, resolution = 0.5)

DimPlot(cortex, reduction = "umap", label=T)
DimPlot(cortex, reduction = "umap", group.by="origin")
DimPlot(cortex, reduction = "umap", group.by="celltype")

saveRDS(cortex, file="cortex_12272022.rds")


##########################colon

colon <- merge(colonmucosalglia, colonMPglia, add.cell.ids=NULL, project="merge")


colon <- SCTransform(colon, vars.to.regress = "percent.mt", assay = "SCT", verbose=FALSE, return.only.var.genes = FALSE)
colon <- RunPCA(colon, features = VariableFeatures(object = colon))

stdv <- colon[["pca"]]@stdev
sum.stdv <- sum(colon[["pca"]]@stdev)
percent.stdv <- (stdv / sum.stdv) * 100

cumulative <- cumsum(percent.stdv)

co1 <- which(cumulative > 90 & percent.stdv < 5)[1]
co1

co2 <- sort(which(
  (percent.stdv[1:length(percent.stdv) - 1] - 
     percent.stdv[2:length(percent.stdv)]) > 0.1), 
  decreasing = T)[1] + 1

co2

min.pc <- min(co1, co2)
min.pc


plot_df <- data.frame(pct = percent.stdv, 
                      cumu = cumulative, 
                      rank = 1:length(percent.stdv))

ggplot(plot_df, aes(cumulative, percent.stdv, label = rank, color = rank > min.pc)) + 
  geom_text() + 
  geom_vline(xintercept = 90, color = "grey") + 
  geom_hline(yintercept = min(percent.stdv[percent.stdv > 5]), color = "grey") +
  theme_bw()


colon <- RunHarmony(colon, group.by.vars="orig.ident", assay.use="SCT")
colon <- RunUMAP(colon, dims=1:min.pc, reduction="harmony")
colon <- FindNeighbors(colon, dims = 1:min.pc, assay="SCT", reduction="harmony")
colon <- FindClusters(colon, resolution = 0.5)

DimPlot(colon, reduction = "umap", label=T)
DimPlot(colon, reduction = "umap", group.by="origin")
DimPlot(colon, reduction = "umap", group.by="celltype")

saveRDS(colon, file="colon_12272022.rds")
##########################duoEGCs, VMHglia, PNS_glia, colon, cortex

gut <- merge(duoEGCs, colon, add.cell.ids=NULL, project="merge")

###sctransform
gut <- SCTransform(gut, vars.to.regress = "percent.mt", assay = "SCT", verbose=FALSE, return.only.var.genes = FALSE)


# Determine percent of variation associated with each 
gut <- RunPCA(gut, features = VariableFeatures(object = gut))

stdv <- gut[["pca"]]@stdev
sum.stdv <- sum(gut[["pca"]]@stdev)
percent.stdv <- (stdv / sum.stdv) * 100

# Calculate cumulative percents for each PC
cumulative <- cumsum(percent.stdv)

# Determine which PC exhibits cumulative percent greater than 90% and
# and % variation associated with the PC as less than 5
co1 <- which(cumulative > 90 & percent.stdv < 5)[1]
co1

# Determine the difference between variation of PC and subsequent PC
co2 <- sort(which(
  (percent.stdv[1:length(percent.stdv) - 1] - 
     percent.stdv[2:length(percent.stdv)]) > 0.1), 
  decreasing = T)[1] + 1

# last point where change of % of variation is more than 0.1%.
co2

# Minimum of the two calculation
min.pc <- min(co1, co2)
min.pc


# Create a dataframe with values
plot_df <- data.frame(pct = percent.stdv, 
                      cumu = cumulative, 
                      rank = 1:length(percent.stdv))

# Elbow plot to visualize 
ggplot(plot_df, aes(cumulative, percent.stdv, label = rank, color = rank > min.pc)) + 
  geom_text() + 
  geom_vline(xintercept = 90, color = "grey") + 
  geom_hline(yintercept = min(percent.stdv[percent.stdv > 5]), color = "grey") +
  theme_bw()


##harmony
gut <- RunHarmony(gut, group.by.vars="orig.ident", assay.use="SCT")
gut <- RunUMAP(gut, dims=1:min.pc, reduction="harmony")
gut <- FindNeighbors(gut, dims = 1:min.pc, assay="SCT", reduction="harmony")
gut <- FindClusters(gut, resolution = 0.5)

DimPlot(gut, reduction = "umap", label=T)
DimPlot(gut, reduction="umap", group.by="origin")
DimPlot(gut, reduction = "umap", group.by="origin", cols=c('colonMP0'='firebrick4', 'colonMP1'='darkorange3', 
                                                                    'colonMP2'='sienna', 'colonMP3'='saddlebrown', 'colonMP4'='coral1', 'colonMP5'='salmon', 'female_duo1'='purple', 
                                                                    'female_duo2'='orchid', 'male_duo1'='slateblue', 'male_duo2'='plum3', 'colonmucosa0'='khaki3', 'colonmucosa1'='navajowhite', 
                                                                    'colonmucosa2'='wheat2', 'colonmucosa3'='darkgoldenrod3', 'colonmucosa4'='gold3'))

DimPlot(gut, reduction="umap", group.by="tissue")
DimPlot(gut, reduction = "umap", group.by="tissue", cols=c('colonMP'='saddlebrown', 'duo0'='purple', 
                                                           'duo1'='plum1', 'duo2'='slateblue', 'duo3'='deepskyblue3', 'duo4'='dodgerblue4', 'duo5'='hotpink', 'colonmucosa'='gold3'))
DimPlot(gut, reduction = "umap", group.by="celltype")


###############
pt <- table(Idents(gut), gut$tissue)
pt <- as.data.frame(pt)
pt$Var1 <- as.character(pt$Var1)

ggplot(pt, aes(x=Var1, y=Freq, fill=Var2)) + theme_bw(base_size=15) + geom_col(position="fill", width=0.5) + xlab("Sample") + ylab("Proportion") + theme(legend.title=element_blank()) + scale_fill_manual(values = c('colonMP'='saddlebrown', 'duo0'='purple', 
                                                                                                                                                                                                                      'duo1'='purple', 'duo2'='purple', 'duo3'='purple', 'duo4'='purple', 'duo5'='purple', 'colonmucosa'='gold3'))
  
  
saveRDS(gut, file="gutglia_12272022.rds")


###################################
duo_vmh <- merge(duoEGCs, VMHglia, add.cell.ids=NULL, project="merge")
two <- merge(PNSglia, cortex, add.cell.ids=NULL, project="merge")
globalglia <- merge(duo_vmh, two, add.cell.ids=NULL, project="merge")


###sctransform
globalglia <- SCTransform(globalglia, vars.to.regress = "percent.mt", assay = "SCT", verbose=FALSE, return.only.var.genes = FALSE)


# Determine percent of variation associated with each 
globalglia <- RunPCA(globalglia, features = VariableFeatures(object = globalglia))

stdv <- globalglia[["pca"]]@stdev
sum.stdv <- sum(globalglia[["pca"]]@stdev)
percent.stdv <- (stdv / sum.stdv) * 100

# Calculate cumulative percents for each PC
cumulative <- cumsum(percent.stdv)

# Determine which PC exhibits cumulative percent greater than 90% and
# and % variation associated with the PC as less than 5
co1 <- which(cumulative > 90 & percent.stdv < 5)[1]
co1

# Determine the difference between variation of PC and subsequent PC
co2 <- sort(which(
  (percent.stdv[1:length(percent.stdv) - 1] - 
     percent.stdv[2:length(percent.stdv)]) > 0.1), 
  decreasing = T)[1] + 1

# last point where change of % of variation is more than 0.1%.
co2

# Minimum of the two calculation
min.pc <- min(co1, co2)
min.pc


# Create a dataframe with values
plot_df <- data.frame(pct = percent.stdv, 
                      cumu = cumulative, 
                      rank = 1:length(percent.stdv))

# Elbow plot to visualize 
ggplot(plot_df, aes(cumulative, percent.stdv, label = rank, color = rank > min.pc)) + 
  geom_text() + 
  geom_vline(xintercept = 90, color = "grey") + 
  geom_hline(yintercept = min(percent.stdv[percent.stdv > 5]), color = "grey") +
  theme_bw()


##harmony
globalglia <- RunHarmony(globalglia, group.by.vars="orig.ident", assay.use="SCT")
globalglia <- RunUMAP(globalglia, dims=1:min.pc, reduction="harmony")
globalglia <- FindNeighbors(globalglia, dims = 1:min.pc, assay="SCT", reduction="harmony")
globalglia <- FindClusters(globalglia, resolution = 0.2)

DimPlot(globalglia, reduction = "umap", label=T, group.by="celltype")


globalglia1 <- merge(globalglia, vagalglia, add.cell.ids=NULL, project="merge")


###sctransform
globalglia1 <- SCTransform(globalglia1, vars.to.regress = "percent.mt", assay = "SCT", verbose=FALSE, return.only.var.genes = FALSE)


# Determine percent of variation associated with each 
globalglia1 <- RunPCA(globalglia1, features = VariableFeatures(object = globalglia1))

stdv <- globalglia1[["pca"]]@stdev
sum.stdv <- sum(globalglia1[["pca"]]@stdev)
percent.stdv <- (stdv / sum.stdv) * 100

# Calculate cumulative percents for each PC
cumulative <- cumsum(percent.stdv)

# Determine which PC exhibits cumulative percent greater than 90% and
# and % variation associated with the PC as less than 5
co1 <- which(cumulative > 90 & percent.stdv < 5)[1]
co1

# Determine the difference between variation of PC and subsequent PC
co2 <- sort(which(
  (percent.stdv[1:length(percent.stdv) - 1] - 
     percent.stdv[2:length(percent.stdv)]) > 0.1), 
  decreasing = T)[1] + 1

# last point where change of % of variation is more than 0.1%.
co2

# Minimum of the two calculation
min.pc <- min(co1, co2)
min.pc


# Create a dataframe with values
plot_df <- data.frame(pct = percent.stdv, 
                      cumu = cumulative, 
                      rank = 1:length(percent.stdv))

# Elbow plot to visualize 
ggplot(plot_df, aes(cumulative, percent.stdv, label = rank, color = rank > min.pc)) + 
  geom_text() + 
  geom_vline(xintercept = 90, color = "grey") + 
  geom_hline(yintercept = min(percent.stdv[percent.stdv > 5]), color = "grey") +
  theme_bw()


##harmony
globalglia1 <- RunHarmony(globalglia1, group.by.vars="orig.ident", assay.use="SCT")
globalglia1 <- RunUMAP(globalglia1, dims=1:min.pc, reduction="harmony")
globalglia1 <- FindNeighbors(globalglia1, dims = 1:min.pc, assay="SCT", reduction="harmony")
DimPlot(globalglia1, reduction = "umap", label=T, group.by="celltype")


DimPlot(globalglia1, reduction = "umap", label=T, group.by="celltype")
DimPlot(globalglia1, reduction = "umap", label=T, group.by="tissue")
DimPlot(globalglia1, reduction = "umap", label=T, group.by="origin")
DimPlot(globalglia1, reduction = "umap", label=T, group.by="orig.ident")

DimPlot(globalglia, reduction = "umap", label=T)


DimPlot(globalglia1, reduction = "umap", label=T, group.by="celltype", cols=c("astrocyte"="burlywood3", "EGC"="purple", "ependymal"="dodgerblue", "OL"="salmon", "OPC"="lightsalmon", "satellite"="darkseagreen", "schwann"="cadetblue3", "tanycyte"="deepskyblue4"))

DimPlot(globalglia1, reduction = "umap", label=T, group.by="tissue", cols=c("cortex"="bisque3", "duo0"="purple", "duo1"="purple", "duo2"="purple", 
                                                                          "duo3"="purple", "duo4"="purple", "duo5"="purple", "peroneal"="lightpink",
                                                                          "scg"="thistle4", "drg"="darkseagreen", "sciatic"="olivedrab", "Sciatic"="olivedrab", "spinalcord"="burlywood3", "sural"="yellowgreen", "vagal"="plum3", "vagus"="plum3", "VMH"="tan2"))

DimPlot(globalglia, reduction = "umap", cols=c('0'='dodgerblue4', '1'='skyblue3', '2'='lightskyblue2', '3'='aquamarine3', 
                                                                    '4'='deepskyblue3', '5'='cadetblue', '6'= 'dodgerblue3', '7'='darkseagreen', '8'='olivedrab3', 
                                                                    '9'='palegreen3', '10'='darkolivegreen4', '11'='chartreuse4', '11'='honeydew4',   
                                                                    '12'='seashell4', '13'='khaki3', '14'='burlywood4', '15'='darkslateblue', 
                                                                    '16'='deepskyblue2', '17'='purple', '18'='cornflowerblue', '19'='cyan4', 
                                                                    '20'='yellow3', '21'='forestgreen', '22'='palegreen', '23'='springgreen3', '24'='royalblue', '25'='blue'))

DimPlot(globalglia, reduction = "umap", group.by="origin", label=T)
DimPlot(globalglia, reduction = "umap", group.by="origin", cols=c('female_duo1'='purple', 'female_duo2'='orchid', 
                                                                    'male_duo1'='slateblue', 'male_duo2'='plum3', 
                                                                    'GSE132044_cortex'='burlywood3', 'GSE182098'='green4', 'PNS_SC_GSE182098'='olivedrab', 
                                                                    'sc1'='seagreen3', 'sc2'='seagreen3', 'sc3'='seagreen3', 'sc4'='seagreen3', 'sc5'='seagreen3',
                                                                    'sc6'='seagreen3', 'sc7'='seagreen3', 'sc8'='seagreen3', 'sc9'='seagreen3', 'Tesar_cortex'='lightpink2',
                                                                    'VMH'='indianred3'))


DimPlot(globalglia, reduction = "umap", group.by="tissue", label=T)
DimPlot(globalglia, reduction = "umap", group.by="tissue", cols=c('duo0'='purple', 'duo1'='purple', 'duo2'='purple', 'duo3'='purple', 'duo4'='purple', 'duo5'='purple', 
                                                                    'peroneal'='steelblue2', 'sciatic'='olivedrab', 
                                                                    'Sciatic'='olivedrab', 'spinalcord'='seagreen3', 'sural'='blue', 
                                                                    'vagus'='black', 'VMH'='indianred3', 'cortex'='burlywood3'))

DimPlot(globalglia, reduction = "umap", group.by="tissue", cols=c('duo0'='purple', 'duo1'='plum1', 'duo2'='slateblue', 'duo3'='deepskyblue3', 'duo4'='dodgerblue4', 'duo5'='hotpink', 
                                                                'peroneal'='steelblue2', 'sciatic'='olivedrab', 
                                                                'Sciatic'='olivedrab', 'spinalcord'='seagreen3', 'sural'='blue', 
                                                                'vagus'='black', 'VMH'='indianred3', 'cortex'='burlywood3'))

                                                           

DimPlot(globalglia, reduction = "umap", group.by="celltype", label=T)
DimPlot(globalglia, reduction = "umap", group.by="celltype", cols=c('astrocyte'='steelblue3', 'EGC'='purple', 
                                                                    'OPC'='dodgerblue4', 'OL'='cadetblue3', 
                                                                    'schwann'='olivedrab'))


###############
pt <- table(Idents(globalglia), globalglia$origin)
pt <- as.data.frame(pt)
pt$Var1 <- as.character(pt$Var1)

ggplot(pt, aes(x=Var1, y=Freq, fill=Var2)) + theme_bw(base_size=15) + geom_col(position="fill", width=0.5) + xlab("Sample") + ylab("Proportion") + theme(legend.title=element_blank()) + scale_fill_manual(values = c('female_duo1'='purple', 'female_duo2'='orchid', 
                                                                                                                                                                                                                      'male_duo1'='slateblue', 'male_duo2'='plum3', 
                                                                                                                                                                                                                      'GSE132044_cortex'='burlywood3', 'GSE182098'='green4', 'PNS_SC_GSE182098'='olivedrab', 
                                                                                                                                                                                                                      'sc1'='seagreen3', 'sc2'='seagreen3', 'sc3'='seagreen3', 'sc4'='seagreen3', 'sc5'='seagreen3',
                                                                                                                                                                                                                      'sc6'='seagreen3', 'sc7'='seagreen3', 'sc8'='seagreen3', 'sc9'='seagreen3', 'Tesar_cortex'='lightpink2',
                                                                                                                                                                                                                      'VMH'='indianred3'))

saveRDS(globalglia1, file="globalglia.rds")

UMAPPlot(globalglia, label=T, 
         cols=c('0'='dodgerblue4', '1'='skyblue3', '2'='lightskyblue2', '3'='aquamarine3', 
                '4'='deepskyblue3', '5'='cadetblue', '6'= 'dodgerblue3', '7'='darkseagreen', '8'='olivedrab3', 
                '9'='palegreen3', '10'='darkolivegreen4', '11'='chartreuse4', '11'='hallgliaydew4',   
                '12'='seashell4', '13'='khaki3', '14'='springgreen3', '15'='purple', 
                '16'='deepskyblue', '17'='darkslateblue', '18'='cornflowerblue', '19'='cyan4', 
                '20'='yellow3', '21'='forestgreen', '22'='palegreen'))


#################################

##colonglia, PNS glia, CNS glia, duodenal glia
allglia <- merge(globalglia, colon, add.cell.ids=NULL, project="merge")


###sctransform
allglia <- SCTransform(allglia, vars.to.regress = "percent.mt", assay = "SCT", verbose=FALSE, return.only.var.genes = FALSE)


# Determine percent of variation associated with each 
allglia <- RunPCA(allglia, features = VariableFeatures(object = allglia))

stdv <- allglia[["pca"]]@stdev
sum.stdv <- sum(allglia[["pca"]]@stdev)
percent.stdv <- (stdv / sum.stdv) * 100

# Calculate cumulative percents for each PC
cumulative <- cumsum(percent.stdv)

# Determine which PC exhibits cumulative percent greater than 90% and
# and % variation associated with the PC as less than 5
co1 <- which(cumulative > 90 & percent.stdv < 5)[1]
co1

# Determine the difference between variation of PC and subsequent PC
co2 <- sort(which(
  (percent.stdv[1:length(percent.stdv) - 1] - 
     percent.stdv[2:length(percent.stdv)]) > 0.1), 
  decreasing = T)[1] + 1

# last point where change of % of variation is more than 0.1%.
co2

# Minimum of the two calculation
min.pc <- min(co1, co2)
min.pc


# Create a dataframe with values
plot_df <- data.frame(pct = percent.stdv, 
                      cumu = cumulative, 
                      rank = 1:length(percent.stdv))

# Elbow plot to visualize 
ggplot(plot_df, aes(cumulative, percent.stdv, label = rank, color = rank > min.pc)) + 
  geom_text() + 
  geom_vline(xintercept = 90, color = "grey") + 
  geom_hline(yintercept = min(percent.stdv[percent.stdv > 5]), color = "grey") +
  theme_bw()


##harmony
allglia <- RunHarmony(allglia, group.by.vars="orig.ident", assay.use="SCT")
allglia <- RunUMAP(allglia, dims=1:min.pc, reduction="harmony")
allglia <- FindNeighbors(allglia, dims = 1:min.pc, assay="SCT", reduction="harmony")
allglia <- FindClusters(allglia, resolution = 0.5)

DimPlot(allglia, reduction = "umap", label=T)
DimPlot(allglia, reduction = "umap", group.by="orig.ident", cols=c('brain'='dodgerblue4', 'femalemouse_duodenum'='purple', 
                                                                    'GSE172204'='skyblue3', 'malehuman_duodenum'='purple', 
                                                                    'malevehiclemouse_duodenum'='purple', 'schwann'='cadetblue', 'SeuratProject'='darkseagreen', 
                                                                    'spinalcordglia'='olivedrab3', 'vehicle_duo_female'='purple'))
DimPlot(allglia, reduction = "umap", group.by="celltype")


###############
pt <- table(Idents(allglia), allglia$orig.ident)
pt <- as.data.frame(pt)
pt$Var1 <- as.character(pt$Var1)

ggplot(pt, aes(x=Var1, y=Freq, fill=Var2)) + theme_bw(base_size=15) + geom_col(position="fill", width=0.5) + xlab("Sample") + ylab("Proportion") + theme(legend.title=element_blank())
ggplot(pt, aes(x=Var1, y=Freq, fill=Var2)) + theme_bw(base_size=15) + geom_col(position="fill", width=0.5) + xlab("Sample") + ylab("Proportion") + theme(legend.title=element_blank()) + scale_fill_manual(values = c('0'='saddlebrown', '1'='saddlebrown', '2'='saddlebrown', '3'='saddlebrown', '4'= 'saddlebrown', 'mucosacolon'='dodgerblue3', 'femalemouse_duodenum'='thistle3', 'malehuman_duodenum'='orchid2', 'malevehiclemouse_duodenum'='mediumpurple', 'vehicle_duo_female'='darkviolet'))

saveRDS(allglia, file="allglia_12302022.rds")

UMAPPlot(allglia, label=T, 
         cols=c('0'='dodgerblue4', '1'='skyblue3', '2'='lightskyblue2', '3'='aquamarine3', 
                '4'='deepskyblue3', '5'='cadetblue', '6'= 'dodgerblue3', '7'='darkseagreen', '8'='olivedrab3', 
                '9'='palegreen3', '10'='darkolivegreen4', '11'='chartreuse4', '11'='hallgliaydew4',   
                '12'='seashell4', '13'='khaki3', '14'='springgreen3', '15'='purple', 
                '16'='deepskyblue', '17'='darkslateblue', '18'='cornflowerblue', '19'='cyan4', 
                '20'='yellow3', '21'='forestgreen', '22'='palegreen'))
```
