## Supplementary material for "Specialized enteric glial cells coordinate intestinal motility": Auxiliary Code Files: Auxiliary File 3.R

```
library(pheatmap)
library(monocle3)
library(dplyr)
library(ggplot2)


###convert Seurat to cds
data <- as(as.matrix(ENS@assays$RNA@data), 'sparseMatrix')

fData <- data.frame(gene_short_name = row.names(data), row.names = row.names(data))
fd <- new('AnnotatedDataFrame', data = fData)
monocle_cds <- new_cell_data_set(data,
                                cell_metadata  = ENS@meta.data,
                                gene_metadata  = fData)

###preprocess data 
monocle_cds <- preprocess_cds(monocle_cds, num_dim=100)

##look at the fraction of variation explained by each PC
plot_pc_variance_explained(monocle_cds)

monocle_cds <- reduce_dimension(monocle_cds)
plot_cells(monocle_cds)
plot_cells(monocle_cds, color_cells_by="orig.ident", cell_size = 1)
plot_cells(monocle_cds, color_cells_by="seurat_clusters",  cell_size = 1)
plot_cells(monocle_cds, color_cells_by="orig.ident", label_cell_groups=FALSE)


###batch correct
monocle_cds <- align_cds(monocle_cds, num_dim = 100, alignment_group = "orig.ident")


##reduce dimensions
monocle_cds <- reduce_dimension(monocle_cds)
plot_cells(monocle_cds, color_cells_by="orig.ident", label_cell_groups=FALSE)

###cluster cells
monocle_cds <- cluster_cells(monocle_cds, resolution=0.04)
plot_cells(monocle_cds, cell_size = 1)


###predict start and end points
plot_cells(monocle_cds, genes=c("Plp1", "Nfia", "Rbfox3", "Gfap"))


###find DEGs per cluster
marker_test_res <- top_markers(monocle_cds, group_cells_by="cluster", 
                               reference_cells=1000, cores=8)
top_specific_markers <- marker_test_res %>%
  filter(fraction_expressing >= 0.10) %>%
  group_by(cell_group) %>%
  top_n(100, pseudo_R2)

top_specific_marker_ids <- unique(top_specific_markers %>% pull(gene_id))

plot_genes_by_group(monocle_cds,
                    top_specific_marker_ids,
                    group_cells_by="cluster",
                    ordering_type="maximal_on_diag",
                    max.size=2)


plot_cells(monocle_cds)
plot_cells(monocle_cds, genes=c("Plp1", "Nfia", "Rbfox3", "Gfap"))
monocle_cds <- learn_graph(monocle_cds)
monocle_cds <- order_cells(monocle_cds)
plot_cells(monocle_cds)


plot_cells(monocle_cds,
           color_cells_by = "pseudotime",
           label_cell_groups=FALSE,
           label_leaves=FALSE,
           label_branch_points=FALSE,
           graph_label_size=1.5, cell_size = 1)

###DEG
gois <- c("Gfap", "Ptprz1", "Piezo2", "Nfia", "Plp1", "Rbfox3", "Ppard", "Vim", "Yap1", "Sox6", "Robo2", "Ghr", "Nrp2", "Nrp1", "Gap43", "S100b")
cds_subset <- monocle_cds[rowData(monocle_cds)$gene_short_name %in% gois,]

gene_fits <- fit_models(cds_subset, model_formula_str = "~cluster")

fit_coefs2 <- coefficient_table(gene_fits)

fit_coefs2 %>% filter (q_value < 0.05) %>%
  select(gene_short_name, term, q_value, estimate)

plot_genes_violin(cds_subset, group_cells_by="seurat_clusters", ncol=2) +
  theme(axis.text.x=element_text(angle=45, hjust=1))


pr_graph_test_res <- graph_test(monocle_cds, neighbor_graph="knn", cores=8)
pr_deg_ids <- row.names(subset(pr_graph_test_res, q_value < 0.05))

####map gene modules - resolution can be changed to lower or raise modules called
gene_module_df <- find_gene_modules(cds_object[pr_deg_ids,], resolution=0.0005)


cell_group_df <- tibble::tibble(cell=row.names(colData(monocle_cds)), 
                                cell_group=clusters(monocle_cds)[colnames(monocle_cds)])

agg_mat <- aggregate_gene_expression(monocle_cds, gene_module_df, cell_group_df)
row.names(agg_mat) <- stringr::str_c("Module", row.names(agg_mat))
colnames(agg_mat) <- stringr::str_c("cluster", colnames(agg_mat))


pheatmap::pheatmap(agg_mat, cluster_rows=TRUE, cluster_cols=TRUE,
                   scale="column", clustering_method="ward.D2",
                   fontsize=6, color=viridis(100))


plot_cells(monocle_cds, 
           genes=gene_module_df,
           group_cells_by="cluster",
           color_cells_by="cluster",
           show_trajectory_graph=FALSE, cell_size = 1)

plot_cells(monocle_cds, 
           genes=gene_module_df %>% filter(module %in% c(8, 28, 33, 161)),
           group_cells_by="cluster",
           color_cells_by="pseudotime",
           show_trajectory_graph=FALSE, cell_size = 1)

plot_cells(monocle_cds,
           color_cells_by = "cluster",
           label_groups_by_cluster=FALSE,
           label_leaves=FALSE,
           label_branch_points=FALSE)

ciliated_cds_pr_test_res <- graph_test(monocle_cds, neighbor_graph="principal_graph", cores=4)
pr_deg_ids <- row.names(subset(ciliated_cds_pr_test_res, q_value < 0.05))


plot_cells(monocle_cds, genes=c("Dpp6", "Nrg3", "Epha6", "Slc13a1", "Ephx2", "Abcd3", "Pls1", "Ireb1", "Lipa", "Adipor2"),
           show_trajectory_graph=FALSE,
           label_cell_groups=FALSE,
           label_leaves=FALSE, cell_size = 1)

genes <- c("Piezo2", "Ano1", "Piezo1", "Kcnq5", "Ghr", "Adipor2", "Gip", "Glp2r")

cds_subset <- monocle_cds[rowData(monocle_cds)$gene_short_name %in% genes]
plot_genes_in_pseudotime(cds_subset, color_cells_by = "seurat_clusters")

####choose start branch
EN <- choose_cells(monocle_cds)


genes <- c("Slc13a1", "Gata4", "Slc2a2", "Ddc", "Ppard", "Acox1", "Aldh3a2", "Slc27a2", "Abcd3", "Ephx2", "Kcnq1", "Slc5a1", "Ireb2", "Pls1", "Ezr", "Ephx2", "Lipa")

cds_subset <- EN[rowData(EN)$gene_short_name %in% genes]
plot_genes_in_pseudotime(cds_subset, color_cells_by = "seurat_clusters")


subset_pr_test_res <- graph_test(EN, neighbor_graph="principal_graph", cores=4)
pr_deg_ids <- row.names(subset(subset_pr_test_res, q_value < 0.05))
gene_module_df <- find_gene_modules(EN[pr_deg_ids,], resolution=0.04)


agg_mat <- aggregate_gene_expression(EN, gene_module_df)
module_dendro <- hclust(dist(agg_mat))
gene_module_df$module <- factor(gene_module_df$module, 
                                levels = row.names(agg_mat)[module_dendro$order])

plot_cells(EN,
           genes=gene_module_df,
           label_cell_groups=FALSE,
           show_trajectory_graph=FALSE)


save_monocle_objects(cds=monocle_cds, directory_path='ENS_monocle', comment='whole duodenum ENS 47 53 clusters cds. Stored 2022-04-18.')

plot_clusters(monocle_cds, clustering, drawSummary = TRUE, sumFun = mean_cl_boot,
              ncol = NULL, nrow = NULL, row_samples = NULL, callout_ids = NULL)


#####proportionmapping

pt <- table(Idents(EGCs_merged), EGCs_merged$orig.ident)
pt <- as.data.frame(pt)
pt$Var1 <- as.character(pt$Var1)

ggplot(pt, aes(x=Var1, y=Freq, fill=Var2)) + theme_bw(base_size=15) + geom_col(position="fill", width=0.5) + xlab("Sample") + ylab("Proportion") + theme(legend.title=element_blank())


ENS$new_column <- paste(ENS$orig.ident,ENS$seurat_clusters)
ENS
```
