## Supplementary material for "Specialized enteric glial cells coordinate intestinal motility": Auxiliary Code Files: Auxiliary File 4.R

```
##install dependencies

if (!requireNamespace("BiocManager", quietly = TRUE))
  +     install.packages("BiocManager")
BiocManager::install("DropletUtils")


if (!requireNamespace("BiocManager", quietly = TRUE))
  +     install.packages("BiocManager")
BiocManager::install("scater")


##load packages

library(DropletUtils)
library(biomaRt)
library(dplyr)
library(scater)
library(celda)
library(Seurat)
library(plotly)
library(e1071)
library(sctransform)
library(viridis)
library(gplots)
library(ggbiplot2)

###load data

mouseduo <- Read10X_h5("....cellbender_filtered.h5")
mouseduo <- CreateSeuratObject(mouseduo, min.cells=3, project="malemouse_duodenum")


###remove low quality cells
mouseduo[["kcnq1ot1"]] <- PercentageFeatureSet(object = mouseduo, pattern = "Kcnq1ot1")
head(mouseduo@meta.data) 
VlnPlot(object = mouseduo, features = c("nFeature_RNA", "nCount_RNA", "kcnq1ot1"), ncol = 3)

###remove mitochondrial reads
mouseduo[["percent.mt"]] <- PercentageFeatureSet(object = mouseduo, pattern = "^mt-")
head(mouseduo@meta.data) 
VlnPlot(object = mouseduo, features = c("nFeature_RNA", "nCount_RNA", "percent.mt"), ncol = 3)

plot1 <- FeatureScatter(mouseduo, feature1 = "nCount_RNA", feature2 = "kcnq1ot1")
plot2 <- FeatureScatter(mouseduo, feature1 = "nCount_RNA", feature2 = "nFeature_RNA")
CombinePlots(plots = list(plot1, plot2))
FeatureScatter(object = mouseduo, feature1 = "nCount_RNA", feature2 = "nFeature_RNA")

str(mouseduo)
VlnPlot(object = mouseduo, features = c("nFeature_RNA"), group.by = c('orig.ident'))

mouseduo


###subset out 
mouseduo <- subset(mouseduo, nFeature_RNA < 8000)
mouseduo <- subset(mouseduo, nFeature_RNA > 500)
mouseduo <- subset(mouseduo, kcnq1ot1 < 2)
mouseduo <- subset(mouseduo, percent.mt < 5)
VlnPlot(object = mouseduo, features = c("nFeature_RNA", "nCount_RNA", "kcnq1ot1"), ncol = 3)


#post processing cell number
mouseduo


###sctransform
mouseduo <- SCTransform(mouseduo, vars.to.regress = "percent.mt", return.only.var.genes = FALSE)

  
##pca
mouseduo <- RunPCA(mouseduo, features = VariableFeatures(object = mouseduo))


# Determine percent of variation associated with each PC
stdv <- mouseduo[["pca"]]@stdev
sum.stdv <- sum(mouseduo[["pca"]]@stdev)
percent.stdv <- (stdv / sum.stdv) * 100

# Calculate cumulative percents for each PC
cumulative <- cumsum(percent.stdv)

# Determine which PC exhibits cumulative percent greater than 90% and
# and % variation associated with the PC as less than 5
co1 <- which(cumulative > 90 & percent.stdv < 5)[1]
co1

# Determine the difference between variation of PC and subsequent PC
co2 <- sort(which(
  (percent.stdv[1:length(percent.stdv) - 1] - 
     percent.stdv[2:length(percent.stdv)]) > 0.1), 
  decreasing = T)[1] + 1

# last point where change of % of variation is more than 0.1%.
co2

# Minimum of the two calculation
min.pc <- min(co1, co2)
min.pc


# Create a dataframe with values
plot_df <- data.frame(pct = percent.stdv, 
                      cumu = cumulative, 
                      rank = 1:length(percent.stdv))

# Elbow plot to visualize 
ggplot(plot_df, aes(cumulative, percent.stdv, label = rank, color = rank > min.pc)) + 
  geom_text() + 
  geom_vline(xintercept = 90, color = "grey") + 
  geom_hline(yintercept = min(percent.stdv[percent.stdv > 5]), color = "grey") +
  theme_bw()


###umap
mouseduo <- RunUMAP(mouseduo, dims = 1:min.pc)
mouseduo <- FindNeighbors(mouseduo, dims = 1:min.pc)
mouseduo <- FindClusters(mouseduo, resolution = 1)

DimPlot(mouseduo, reduction = "umap", pt.size=1)


mouseduo

##SAVE
saveRDS(mouseduo, file = "mouseduo.rds")


####MERGE

duo <- merge(mouseduo, female_duo, add.cell.ids=c("3", "female"), project="vehicle_data")

duo

###sctransform
duo <- SCTransform(duo, vars.to.regress = "percent.mt", verbose=FALSE, return.only.var.genes = FALSE)

##pca
duo <- RunPCA(duo, features = VariableFeatures(object = duo))

# Determine percent of variation associated with each PC
stdv <- duo[["pca"]]@stdev
sum.stdv <- sum(duo[["pca"]]@stdev)
percent.stdv <- (stdv / sum.stdv) * 100

# Calculate cumulative percents for each PC
cumulative <- cumsum(percent.stdv)

# Determine which PC exhibits cumulative percent greater than 90% and
# and % variation associated with the PC as less than 5
co1 <- which(cumulative > 90 & percent.stdv < 5)[1]
co1

# Determine the difference between variation of PC and subsequent PC
co2 <- sort(which(
  (percent.stdv[1:length(percent.stdv) - 1] - 
     percent.stdv[2:length(percent.stdv)]) > 0.1), 
  decreasing = T)[1] + 1

# last point where change of % of variation is more than 0.1%.
co2

# Minimum of the two calculation
min.pc <- min(co1, co2)
min.pc


# Create a dataframe with values
plot_df <- data.frame(pct = percent.stdv, 
                      cumu = cumulative, 
                      rank = 1:length(percent.stdv))

# Elbow plot to visualize 
ggplot(plot_df, aes(cumulative, percent.stdv, label = rank, color = rank > min.pc)) + 
  geom_text() + 
  geom_vline(xintercept = 90, color = "grey") + 
  geom_hline(yintercept = min(percent.stdv[percent.stdv > 5]), color = "grey") +
  theme_bw()

##harmony

duo <- RunHarmony(duo, group.by.vars="orig.ident", assay.use="SCT")
duo <- RunUMAP(duo, reduction = "harmony", dims=1:min.pc)
duo <- FindNeighbors(duo, dims = 1:min.pc, reduction="harmony")
duo <- FindClusters(duo, resolution = 1)


DimPlot(duo, reduction = "umap", pt.size=0.8)
DimPlot(duo, reduction = "umap", pt.size=0.8, group.by="orig.ident")


# find markers for every cluster compared to all remaining cells, report only the positive ones
mouseduofinal.markers <- FindAllMarkers(duo, only.pos = TRUE)
mouseduofinal.markers %>% group_by(cluster) %>% top_n(2, avg_logFC)

########
mouseduofinal.markers %>% group_by(cluster) %>% top_n(10, avg_logFC) ->top10

DoHeatmap(duo, features = top10$gene) + scale_fill_viridis(option="viridis")
genes.viz=c("Plp1", "S100b", "Sox10", "Gfap", "Rbfo3")
DotPlot(duo, assay=NULL, genes.viz)
FeaturePlot(duo, genes.viz)

write.table(mouseduofinal.markers, file = "mouseduo_markers.txt", sep = "\t")


##SAVE
saveRDS(duo, file = "mouseduo_final_broad.rds")


##### cell cycle
# read in a list of cell cycle markers, from Tirosh et al, 2015
cc.genes <- readLines(con = "/Users/marissascavuzzo/Downloads/regev_lab_cell_cycle_genes.txt")

# segregate this list into markers of G2/M phase and markers of S
# phase
s.genes <- cc.genes[1:43]
g2m.genes <- cc.genes[44:98]


duo <- CellCycleScoring(object = duo, s.features = s.genes, g2m.features = g2m.genes, 
                                   set.ident = FALSE)

DimPlot(duo, reduction = "umap")
DimPlot(duo, reduction = "umap", group.by="Phase", cols=c("green", "blue", "gray"))


###proportion

library(ggplot2)
pt <- table(Idents(duo), duo$Phase)
pt <- as.data.frame(pt)
pt$Var1 <- as.character(pt$Var1)

ggplot(pt, aes(x=Var1, y=Freq, fill=Var2)) + theme_bw(base_size=15) + geom_col(position="fill", width=0.5) + xlab("Sample") + ylab("Proportion") + theme(legend.title=element_blank())
```
